# BehaviorScope-X: reusing pose-trained visual representations for full-video ethology

**DOI:** 10.64898/2026.07.02.735695

**Authors:** Farhan Augustine, Virginia Murray

## Abstract

Pose-estimation pipelines usually export keypoint coordinates and discard the intermediate visual representations learned to localize animals in a specific assay. We asked whether those discarded representations can be reused for full-video ethology. BehaviorScope-X tests this idea by treating a trained pose checkpoint as both a keypoint estimator and a reusable visual encoder: the pose model is run once to cache detections, keypoints, pose-derived social geometry, and frozen intermediate descriptors, after which compact temporal classifiers are trained on cached multimodal windows. Across MARS resident-intruder videos, cached pose-trained descriptors and pose-derived geometry provided complementary evidence for behavior decoding, recovering sustained behavioral episodes and local sequence structure while revealing a main limitation in dense short-bout regions. The same cache-and-classify design generalized across pose routes, including a MobileNetV3 backbone and a DeepLabCut SuperAnimal HRNet-W32 checkpoint, showing that standard pose workflows can expose behavior-relevant visual descriptors without giving up their keypoint-estimation role. We further tested the approach in Fly-v-Fly aggression, extending the analysis to a second species and shorter behavioral time scale, where sub-second events and annotation-boundary uncertainty limited strict bout recovery. End-to-end profiling showed that the workflow can operate near-real-time or real-time on consumer hardware. Together, these experiments support amortized pose vision as a practical strategy for reusing assay-trained pose models as stable sources of visual and geometric evidence for scalable behavioral analysis.

**Author summary:** Pose-estimation models are usually used to convert animal video into body landmarks, while the same model’s internal visual representations are discarded. We ask whether the model that estimates pose can also provide visual features for behavior analysis. Our workflow runs a trained pose model once, caches its landmarks, pose-derived interaction geometry, and internal visual features, and trains compact behavior classifiers on that cache. Across multiple pose backbones and pose-estimation workflows, these shared pose-and-visual signals supported full-video ethogram recovery without training a separate video network. This turns pose training into a reusable source of assay-specific visual and geometric evidence for scalable behavioral analysis.

**Highlights:**

- A pose-estimation checkpoint is reused as both keypoint estimator and visual encoder.
- A single pose-inference pass caches keypoints, social geometry, and intermediate visual descriptors.
- Cached pose-derived visual and geometric evidence improves full-video behavior decoding beyond pose-derived features alone.
- BehaviorScope-X turns existing pose workflows into reusable full-video behavior-analysis pipelines.

## 1. Introduction

Quantifying animal behavior is a central problem in modern neuroscience, because behavior is the observable output through which neural circuits, disease models, genetic perturbations, and interventions are ultimately interpreted (Anderson and Perona, 2014; Datta, Anderson, Branson, Perona and Leifer, 2019; Pereira, Shaevitz and Murthy, 2020). Over the last decade, the field has made this problem more tractable by separating video-based behavior analysis into two linked stages. First, pose-estimation systems recover experimenter-defined body landmarks from video. Second, downstream models use those coordinates, or features engineered from them, to classify behavior. The pose-first division of labor has been enormously productive. Tools such as DeepLabCut and SLEAP have made it practical to train assay-specific keypoint models at scale (Mathis, Mamidanna, Cury, Abe, Murthy, Mathis and Bethge, 2018; Pereira, Tabris, Matsliah, Turner, Li, Ravindranath, Papadoyannis, Normand, Deutsch, Wang, McKenzie-Smith et al., 2022; Lauer, Zhou, Ye, Menegas, Schneider, Nath, Rahman et al., 2022), while MARS, JAABA, SimBA, JABS, DeepOF, B-SOiD, A-SOiD, VAME, BehaviorAtlas, Keypoint-MoSeq, and related systems have shown how pose-derived measurements can support supervised annotation, active learning, standardized phenotyping, unsupervised syllable discovery, and dynamical modeling (Segalin, Williams, Karigo, Hui, Zelikowski, Sun, Perona, Anderson and Kennedy, 2021c; Kabra, Robie, Rivera-Alba, Branson and Branson, 2013; Goodwin, Choong, Hwang, Pitts, Bloom et al., 2024; Choudhary, Geuther, Sproule, Beane, Kohar, Trapszo and Kumar, 2025; Miranda, Bordes, Pütz, Schmidt and Müller-Myhsok, 2023; Hsu and Yttri, 2021; Tillmann, Hsu, Schwarz and Yttri, 2024; Luxem, Mocellin, Fuhrmann et al., 2022; Huang, Han, Chen, Pan, Zhao, Yi, Li, Liu, Wei, Wang et al., 2021; Weinreb, Pearl, Lin, Osman, Zhang et al., 2024). The durability of pose-based behavior analysis comes from a practical strength: keypoints are compact enough to store, interpretable enough to inspect, and structured enough to reuse across many downstream analyses.

However, that same strength also creates a subtle bottleneck. Once video has been converted into keypoint tables, downstream behavior models often stop revisiting the pixels. In many cases this is desirable, because coordinates reduce nuisance variation and provide a body-centered description of the animal. At the same time, a trained pose model is more than a coordinate exporter. To localize body parts under the lighting, occlusion, contact, background, and animal-appearance conditions of a particular assay, the model must learn internal visual representations of that assay. Yet standard pose pipelines usually discard those intermediate representations at the moment they export final coordinates. The central question motivating this study is whether that discarded pose-trained representation contains behaviorally useful information for full-video ethology.

The question matters because many behaviors are not fully described by sparse landmark geometry. In social assays, labels such as attack, investigation, and mount can depend on contact configuration, body overlap, partial occlusion, local appearance, and scene context in addition to instantaneous skeleton position. Recent systems increasingly recognize this limitation by moving beyond coordinate-only pipelines. Video-centric methods such as FERAL and TRACE use visual streams to map video directly to behavior. Hybrid systems such as LookAgain combine keypoints with selective video evidence. In parallel, emerging vision-language approaches such as BehaviorVLM ask whether pretrained multimodal models can support behavioral understanding with reduced task-specific fine-tuning (Bohnslav, Wimalasena, Clausing, Dai et al., 2021; Harris, Finn, Kieseler, Maechler and Tse, 2023; Hu, Ferrario, Maitland et al., 2023; Skovorodnikov, Zhao, Buck, Kay, Frank, Koger, Costelloe, Couzin and Razzauti, 2025; Shi, Zhang, Wang, Zhang, Tao and Zhang, 2026; Li, Ke, Wang, Li and Wu, 2026; Ke, Li, Pradhan, Markowitz and Wu, 2026). Together, these approaches show that behavior analysis is moving toward hybrid representations that combine pose, visual evidence, temporal context, and semantic structure.

BehaviorScope-X works from the same direction but uses a different source of visual evidence: the representation already learned during pose training. We use *amortized pose vision* to describe this principle, in which an assay-trained pose checkpoint is treated as both a keypoint estimator and a reusable source of pose-trained visual evidence for behavior analysis. The scientific question is whether intermediate descriptors, explicit self-pose geometry, and pairwise social geometry retain behavior-discriminative information after coordinate export. The operational consequence is that downstream temporal classifiers can reuse the pose-training investment without training or running a separate raw-video behavior backbone. The representational-operational separation makes the principle testable across pose-model families.

IntegraPose previously unified pose estimation and frame-level behavior classification within a single YOLO multi-task head, with bout-level readouts computed from the frame-wise behavior sequence during postprocessing (Augustine, O’Sullivan, Murray, Ogura, Lin and Singer, 2025). In contrast, the present workflow freezes a completed pose checkpoint and trains temporal sequence classifiers on cached multimodal windows whose bout counts, durations, fragmentation, and transitions can be interpreted biologically. In the present study, YOLO-pose provided the primary ablation substrate, MobileNetV3 tested native backbone dependence, and DeepLabCut SuperAnimal HRNet-W32 tested whether an established DeepLabCut workflow could be made coordinate-and-descriptor reusable.

The upstream-downstream design is evaluated through three computational-biology questions. First, does a validated pose checkpoint contain reusable visual and geometric information for continuous behavior decoding beyond the final coordinates normally exported from pose pipelines? Second, which components of that representation matter for ethogram inference: animal-centered visual evidence, group context, self-pose geometry, pairwise social geometry, temporal decoder capacity, or their interaction? Third, under what conditions does the strategy fail: shorter behavioral time scales, a changed pose backbone, or uncertain manual-annotation bout boundaries? To address these questions, we performed eight linked analyses: (i) validation of the upstream YOLO-pose checkpoint as a reusable representation source; (ii) full-video behavior decoding in held-out MARS resident-intruder recordings; (iii) stream-level ablation of visual, self-pose, and social-geometry features; (iv) analysis of continuous ethogram recovery, local sequence structure, and temporal operating envelopes; (v) static/PCA negative-control testing with Random Forest and XGBoost baselines using window-level inputs; (vi) backbone-dependence and descriptor-provenance testing across YOLO/SPPF, MobileNetV3, and DLC-HRNet, including the DeepLabCut top-down workflow; (vii) short-bout cross-assay stress testing in Fly-v-Fly aggression with annotation-aware evaluation; and (viii) end-to-end throughput benchmarking of the complete video-to-ethogram deployment path. The analyses first evaluate whether cached pose-trained representations support held-out ethogram recovery, then define how that reuse changes across backbones, pose ecosystems, temporal scales, and deployment settings.

## 2. Materials and methods

We designed the BehaviorScope-X workflow as a sequential, multi-stage pipeline that separates upstream pose-trained representation extraction from downstream behavior-classifier training. The upstream pose model is treated as a frozen, assay-adapted representation source, while downstream behavior models are controlled classifiers of workflow-specific caches. The cache-and-classify design lets stream content, temporal decoder family, model capacity, and species-specific temporal scale vary without changing the pose checkpoint, held-out video units, or feature provenance. The Methods follow the cache-and-classify pipeline: datasets and held-out units, video preprocessing and feature preparation, pose-backbone validation, input stream construction, temporal classifiers, classical baselines, DeepLabCut workflow testing, Fly-v-Fly stress testing, postprocessing, evaluation metrics, ethogram analyses, deployment-throughput benchmarking, and the open-source application. The organizing principle is that each comparison changes one downstream question while preserving the provenance of the pose-trained representation being reused. Fig 1 summarizes this workflow and the evidence streams carried forward into temporal behavior decoding.

**Figure 1:**
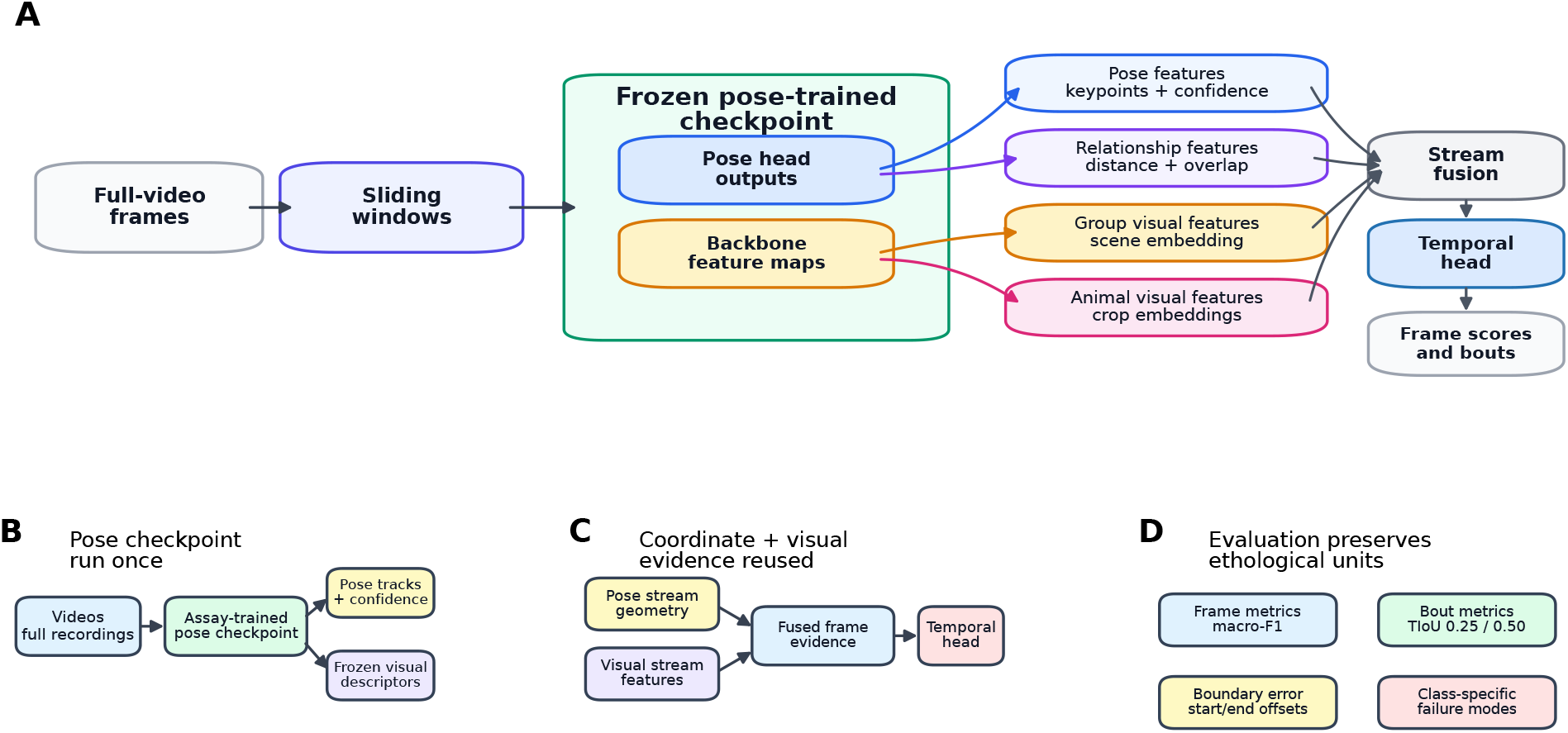
BehaviorScope-X amortized pose vision workflow. (A) Full-video frames are converted into sliding windows and passed through a frozen pose-trained checkpoint. Pose-head outputs provide keypoints, confidence values, and relationship features, while backbone feature maps provide group-scene and per-animal visual descriptors. The synchronized streams are fused and decoded by a temporal head into frame scores and behavior bouts. (B) The pose checkpoint is used to build reusable pose tracks and frozen visual descriptors. (C) Coordinate and visual evidence are combined into fused frame-level evidence before temporal decoding. (D) Evaluation preserves ethological units by reporting frame metrics, bout metrics, boundary errors, and class-specific failure modes.

### 2.1. Datasets and held-out evaluation units

We used the public MARS resident-intruder mouse social-behavior dataset as the primary validation domain for amortized pose vision with a single domain-adapted pose model (Segalin et al., 2021c; Segalin, Williams, Karigo, Hui, Zelikowski, Sun, Perona, Anderson and Kennedy, 2021b,a). The MARS dataset provides full-length interacting-mouse videos, human frame-level annotations, and a social-behavior label set that requires distinguishing attack, investigation, mount, and other behavior. The MARS split used here contained 57 training videos and 33 validation videos. We reserved 30 additional videos for held-out evaluation: 10 from `test_1` and 20 from `test_2`. For ground-truth discovery, we excluded annotations identified as model predictions and retained only human behavior annotations. During discovery, `.annot` files whose filenames contained `pred` or `prediction` were excluded unless explicitly allowed, and behavior/action annotation files were preferred among the remaining human annotation files. This kept the primary MARS evaluation anchored to human-labeled biological videos rather than model-generated labels. After this filtering, 28 of the 30 held-out videos retained accepted human annotations and define the denominator for all reported frame-level and bout-level metrics; the remaining two carried only prediction-derived annotation files.

While MARS served as the primary domain for validating amortized pose vision, we also evaluated a secondary dataset with distinct biological constraints. Fly-v-Fly was used to extend the analysis to cross-assay stress testing and a shorter behavioral time scale (Eyjolfsdottir, Branson, Burgos-Artizzu, Hoopfer, Schor, Anderson and Perona, 2014). The Fly-v-Fly analysis used a public, scope-limited subset of 10 aggression movies with available video, action, feature, and track files. The primary behavior annotations were used for training, validation, and the primary held-out evaluation, with movies 6–10 reserved for behavior-label evaluation. For movie 6, the available secondary human annotation was used for annotation-aware analyses and human-human reference comparisons; model training and checkpoint selection used the primary annotations. As in the MARS held-out analysis, Fly-v-Fly evaluation used a fixed held-out window cache and a fixed frozen YOLO/SPPF feature cache rather than rerunning pose-feature extraction separately for each temporal model. The held-out design changed species, body geometry, behavior repertoire, and bout-duration scale while preserving the same broad modeling idea of pose-derived information, visual information, cached reuse, and temporal classification.

All training and held-out evaluation settings were controlled through a single configuration file before execution. The configuration defined dataset locations, model families, feature streams, temporal-window lengths, optimization settings, postprocessing rules, and held-out evaluation splits. For reproducibility, we saved a frozen copy of this configuration, the full execution plan, per-stage logs, resource measurements, and model-specific configuration summaries for each execution. The study used one fixed experimental matrix rather than a collection of model-specific settings selected after observing performance (Table 1).

**Table 1.**
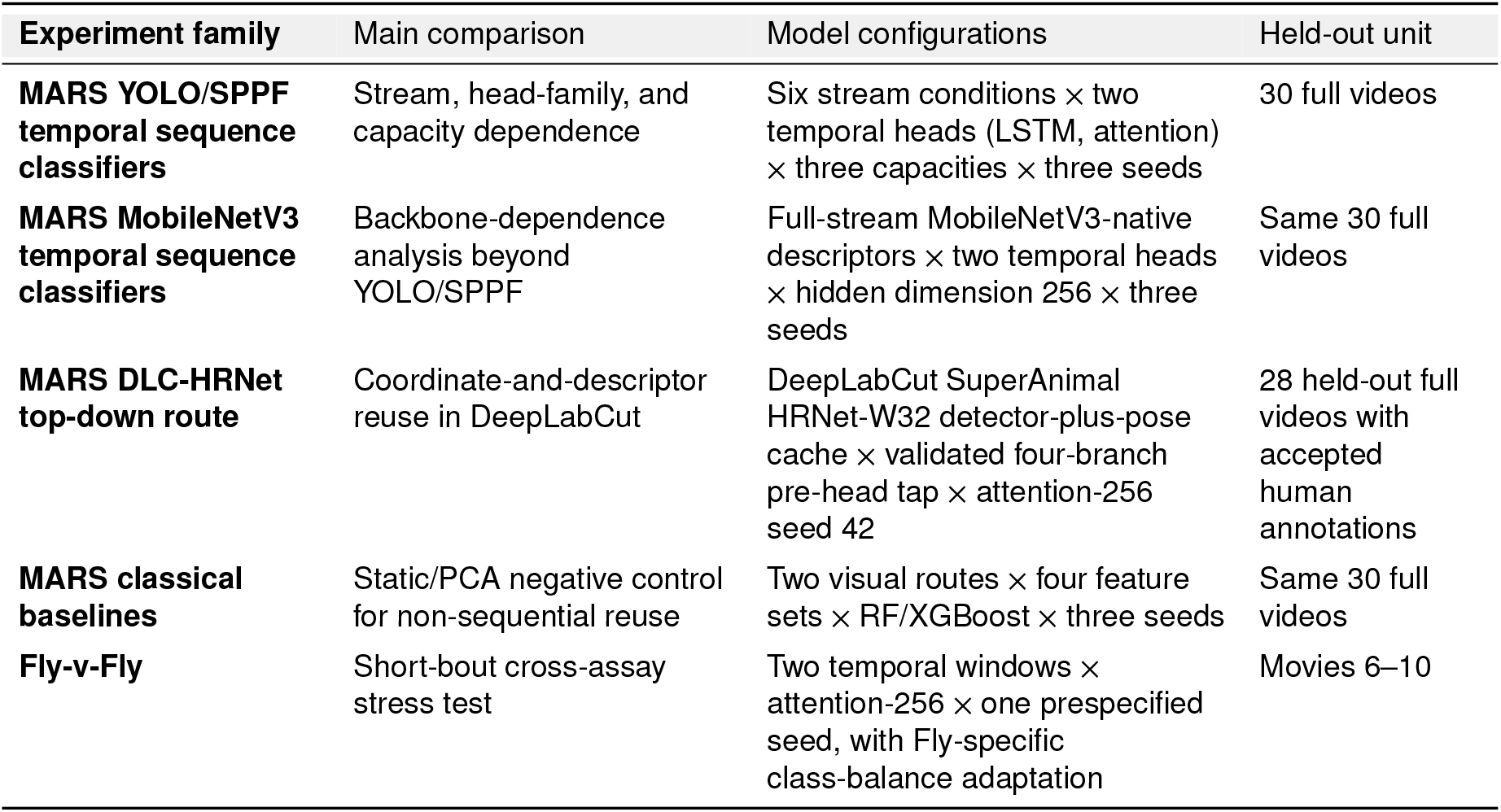
Final BehaviorScope-X experimental matrix. All models were specified by a frozen configuration before execution. The MARS YOLO/SPPF matrix tests stream content, temporal-decoder family, capacity, and seed variability under a shared protocol for evaluating held-out full videos. The MobileNetV3 analysis tests backbone dependence within a native non-YOLO pose route. The DLC-HRNet route instantiates the same cache-and-classify principle in a DeepLabCut SuperAnimal top-down workflow with a different inference style, HRNet feature geometry, and validated pre-head tap. The classical baselines act as a static/PCA negative control for non-sequential reuse by testing Random Forest and XGBoost models under matched temporal access and dimension-reduced visual descriptors. The Fly-v-Fly analysis tests behavior recognition at a shorter temporal scale with annotation-aware handling.

| Experiment family | Main comparison | Model configurations | Held-out unit |
| --- | --- | --- | --- |
| **MARS YOLO/SPPF temporal sequence classifiers** | Stream, head-family, and capacity dependence | Six stream conditions × two temporal heads (LSTM, attention) × three capacities × three seeds | 30 full videos |
| **MARS MobileNetV3 temporal sequence classifiers** | Backbone-dependence analysis beyond YOLO/SPPF | Full-stream MobileNetV3-native descriptors × two temporal heads × hidden dimension 256 × three seeds | Same 30 full videos |
| **MARS DLC-HRNet top-down route** | Coordinate-and-descriptor reuse in DeepLabCut | DeepLabCut SuperAnimal HRNet-W32 detector-plus-pose cache × validated four-branch pre-head tap × attention-256 seed 42 | 28 held-out full videos with accepted human annotations |
| **MARS classical baselines** | Static/PCA negative control for non-sequential reuse | Two visual routes × four feature sets × RF/XGBoost × three seeds | Same 30 full videos |
| **Fly-v-Fly** | Short-bout cross-assay stress test | Two temporal windows × attention-256 × one prespecified seed, with Fly-specific class-balance adaptation | Movies 6–10 |

### 2.2. Video preprocessing and feature preparation

The preprocessing stage converted each video into standardized pose, crop, and visual-feature outputs that served as reusable inputs for downstream behavior modeling. The standardized preprocessing allowed behavior classifiers to be compared while holding frame order, pose checkpoint, crop definitions, and held-out split boundaries fixed.

We converted MARS source videos from NorPix `.seq` files to `.mp4` copies before YOLO-pose inference, using a target frame rate of 30 frames/s, the `mp4v` codec, and up to three retries with a 2-s delay between retries. Standardized frame access keeps downstream differences interpretable as model and representation effects rather than video-decoding artifacts.

For MARS, pose and visual summaries were prepared from the training and validation splits, while `test_1` and `test_2` were reserved for held-out evaluation. We ran the MARS YOLO-pose checkpoint trained on the assay domain over the training video library once, recording its animal detections, seven keypoints per animal, group and per-animal crops, and intermediate visual summaries for behavior-classifier training. For held-out evaluation, the same YOLO-pose checkpoint generated pose and visual streams for the held-out videos, but those held-out summaries were prepared separately from the training summaries. Held-out test features were never reused during training. For the MobileNetV3 cross-backbone analysis, the same MARS window manifests and pose-derived geometry were reused, but visual descriptors were extracted from the frozen MobileNetV3 pose checkpoint into a separate run-specific cache. Thus, the MARS comparisons used matched biological units while preserving route-specific feature-cache provenance.

### 2.3. YOLO-pose backbone trained on the assay domain and visual feature tap

BehaviorScope-X used a YOLO-pose checkpoint trained on the assay domain as the shared upstream perception model. For MARS, this checkpoint was a YOLO26n-pose nano model implemented with the Ultralytics training framework (Ultralytics, 2023). The checkpoint produced animal detections and keypoints, and the same frozen network produced visual descriptors for the behavior classifier. The MARS pose model used seven keypoints per animal (`[7, 3]` keypoint shape), two animals per frame, 224-pixel crops, a pose-confidence threshold of 0.2, an animal crop scale factor of 4.0, and a group crop scale factor of 8.0 (Fig 3). The checkpoint defined the common detection, keypoint, crop, and visual-feature source for the primary MARS reuse experiments.

The MARS pose-training set was constructed from 90 indexed training and validation videos. Candidate frames were sampled from annotated behavior bouts using an intra-bout stride of 6 frames. The candidate pool contained 37,129 investigation frames, 8,808 mount frames, and 7,763 attack frames. To balance pose training across behavior contexts, we targeted 10,000 frames per behavior class, sampling 10,000 investigation frames and retaining all available mount and attack candidates. The sampling procedure produced 26,571 selected frames, which were split 80/20 into 21,256 training frames and 5,315 validation frames. The YOLO26n-pose nano model was then trained for 50 epochs on this MARS pose dataset. The training and validation loss curves, together with detection and pose mAP trajectories, confirmed that the checkpoint converged before being used as a feature extractor (Fig 2).

**Figure 2:**
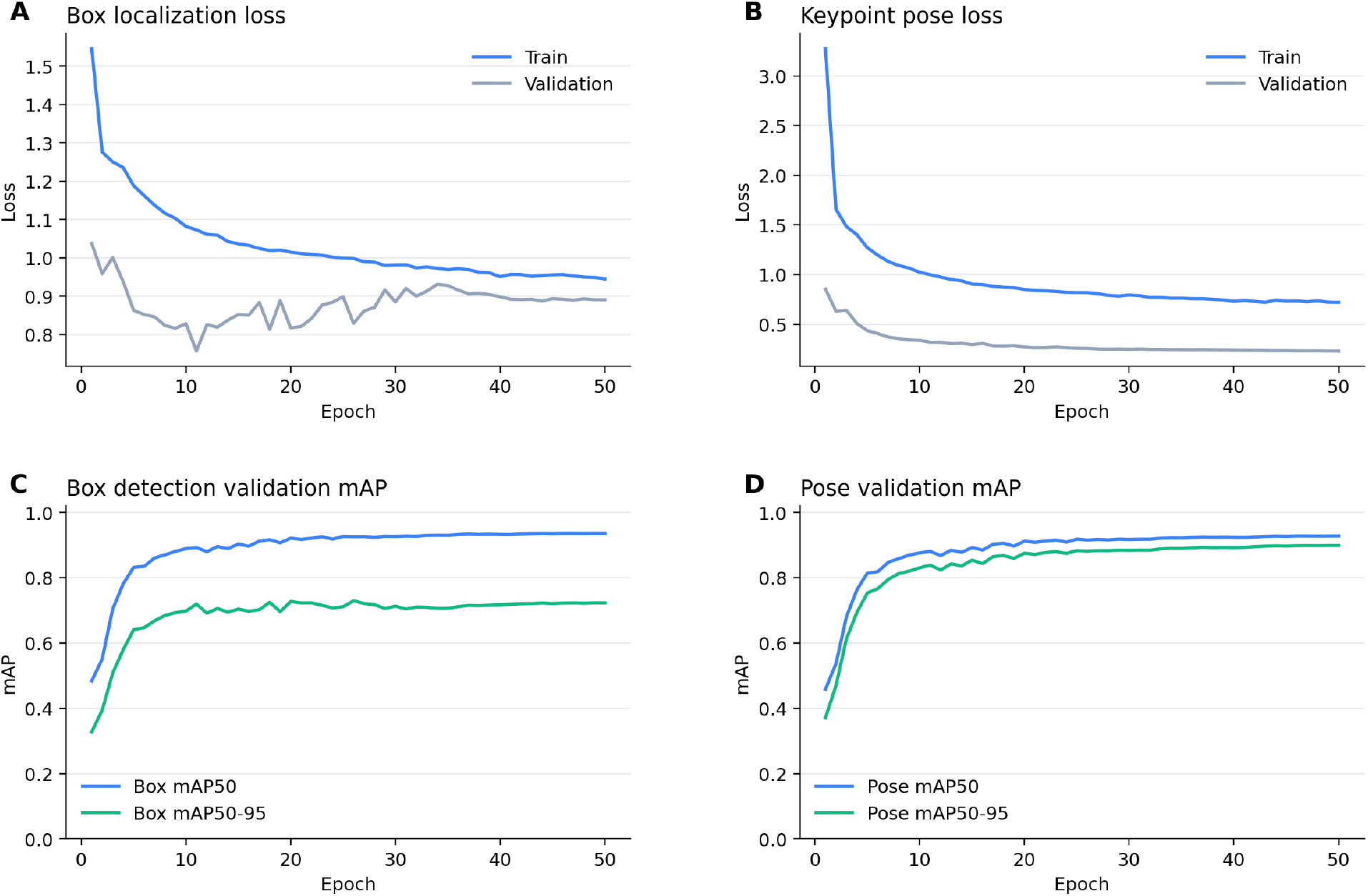
MARS pose-checkpoint convergence over 50 training epochs. (A) Box localization loss and (B) keypoint pose loss both decreased monotonically for training and validation sets. (C) Box detection mAP50 reached 0.936 and mAP50–95 reached 0.723. (D) Pose validation mAP50 reached 0.928 and mAP50–95 reached 0.899. The training curves confirm that the checkpoint had converged before its frozen features were used for downstream behavior classification.

**Figure 3:**
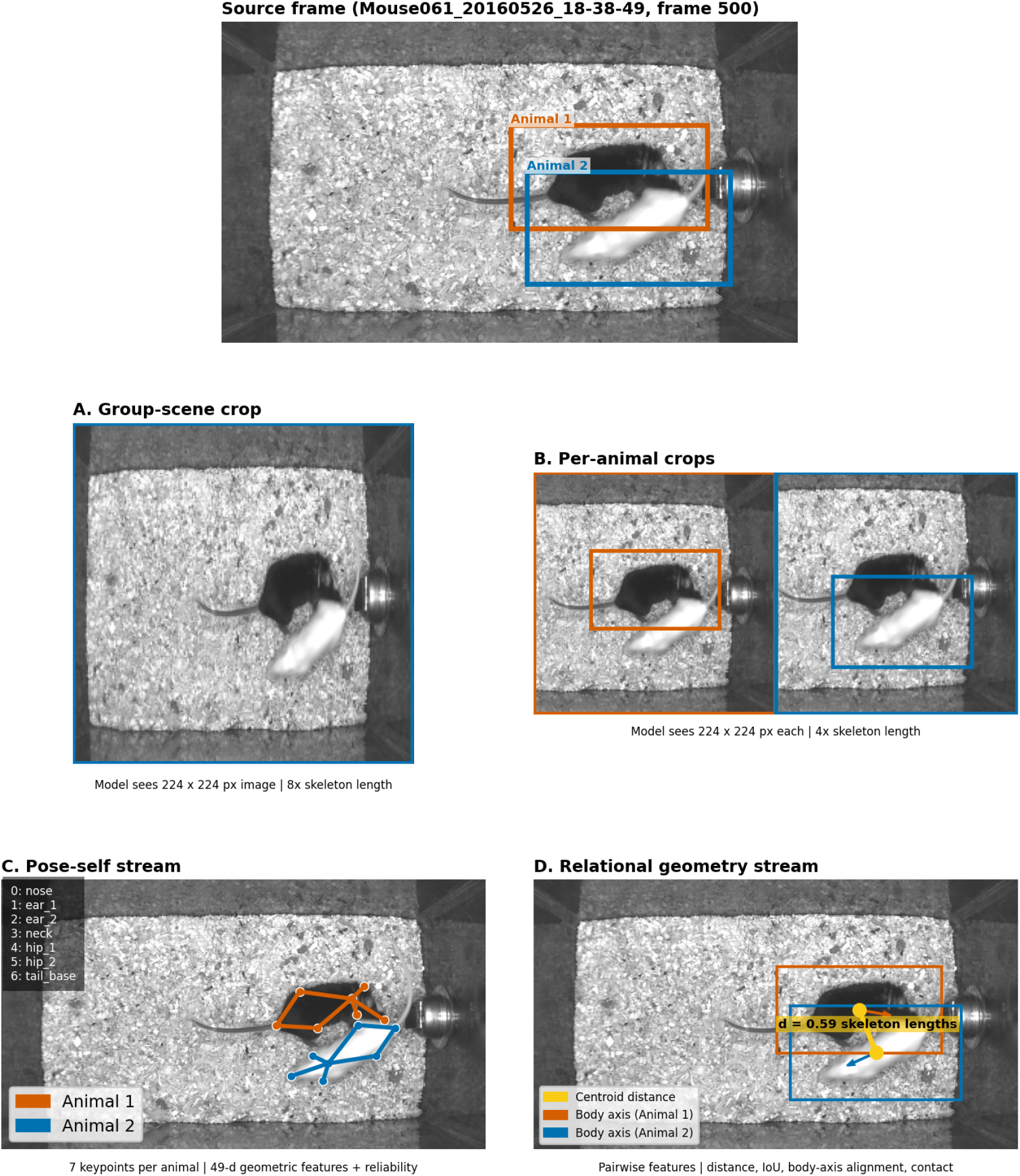
Input-stream decomposition for a representative MARS frame (Mouse061, frame 500). (A) The group-scene crop captures the full interaction context at 224×224 pixels (8× skeleton length). (B) Per-animal crops isolate each detected mouse at 224×224 pixels (4× skeleton length). (C) The pose-self stream encodes seven keypoints per animal into 49-dimensional geometric features plus reliability scalars. (D) The relational geometry stream computes pairwise features (centroid distance, bounding-box IoU, body-axis alignment, approach velocity, and contact indicators) between the two animals. All four streams are derived from the same frozen YOLO-pose checkpoint.

The visual feature tap was chosen to provide a reusable pose-trained backbone representation rather than a detector- or pose-head-specific representation. We took visual descriptors from the final SPPF block of the YOLO backbone, before the detector neck, pose head, and subsequent context-refinement block. SPPF pools spatial context while remaining part of the backbone representation, making it a natural point for extracting reusable appearance descriptors. By contrast, C2PSA applies a later attention-based context-refinement operation closer to the task-specific detection pathway. The retained visual pathway therefore used YOLO layers 0–9, ending at the SPPF output and excluding the C2PSA block, detector neck, and pose head. The extracted 224×224 crop features had shape [*B*, 256, 7, 7], where *B* is the feature-extraction batch size. For one crop, let **F** ∈ ℝ^256×*H*×*W*^ denote the SPPF feature map, with spatial indices *i* and *j*. We obtained a visual descriptor by global average pooling over the spatial dimensions:

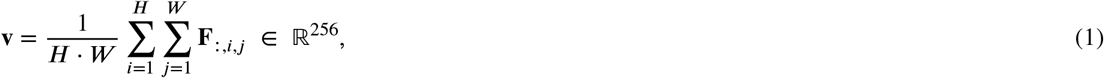

where *H*=*W* =7. Global average pooling was applied identically to the group crop (scale factor 8× skeleton length) and to each per-animal crop (scale factor 4× skeleton length), yielding one 256-dimensional group descriptor 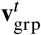 and *N* per-animal descriptors 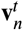. The SPPF-tap procedure fixed the source of visual features across all MARS stream and capacity conditions. Full architectural details of the YOLO26n-pose backbone and SPPF tap are given in the supplementary YOLO/SPPF architecture section, and a multi-scale feature extraction experiment that compared tapping multiple backbone stages against the SPPF-only default is reported in the supplementary multi-scale feature analysis. The key point for the main study is that the visual stream comes from a specified pre-head backbone tap, making feature reuse auditable rather than an implicit byproduct of the detector.

The extraction steps produced a continuous, frame-synchronous feature record for each video: a 768-dimensional visual vector (one 256-d group descriptor plus two 256-d per-animal descriptors) alongside the pose keypoints, all extracted from the same frozen YOLO-pose backbone. The per-frame outputs were cached as compressed float32 NumPy arrays, decoupling behavior-model training from video decoding and backbone inference entirely. During temporal-model training and evaluation, these cached arrays were sliced into overlapping 32-frame sliding windows at stride 16, forming the multi-stream input described in the following section. The downstream experiments vary stream content and temporal classifiers while holding upstream video decoding and pose inference fixed.

Freezing the visual representation was a deliberate constraint that forced any behavior gains to come from information already present in the pose-trained representation and from the temporal classifier’s use of it, rather than from new visual learning. Fine-tuning the backbone for each behavior condition would instead entangle representation learning with the controlled downstream ablations, so we left it as a future comparison.

### 2.4. BehaviorScope-X input streams

The stream design was intended to separate sources of behavior evidence that are often entangled in a single end-to-end video classifier, including scene context, individual appearance, body configuration, and inter-animal geometry. Each MARS temporal window could contain four conceptual streams:

- *Group visual context*: a 256-dimensional descriptor from the group crop, intended to capture the full interaction scene.
- *Per-animal visual descriptors*: a 256-dimensional descriptor for each detected animal crop, intended to capture appearance and local contact evidence around each animal.
- *Pose-self geometry*: a 53-dimensional representation of each animal’s seven-keypoint body configuration, consisting of a 49-dimensional pose vector plus four reliability scalars. The pose vector included centered keypoint coordinates, keypoint confidences, pairwise within-animal distances, and frame-to-frame keypoint velocities.
- *Pairwise relational geometry*: 11 raw inter-animal features per frame, including bounding-box geometry, overlap, relative motion, pose-conditioned distance, orientation, approach velocity, and contact-related features.

Each keypoint’s pose-confidence score from the detector was appended as a reliability scalar, allowing the downstream model to weight well-localized keypoints more heavily than uncertain ones. In the following, *K* denotes the number of keypoints per animal (*K*=7), 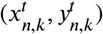 the normalized image coordinates for keypoint *k* of animal *n* at frame *t*, and 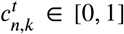 the associated detector confidence. Bounding boxes are denoted 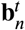 with centroid 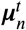 and area 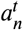; 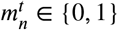 indicates whether animal *n* was detected at frame *t*, and 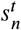 denotes its detector confidence.

#### Pose-self features (49 dimensions per animal)

For each animal *n* at frame *t*, the raw pose-self vector 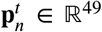 concatenated three sub-vectors. Within-frame coordinate and distance terms omit the superscript *t* when all values refer to the same frame. The first sub-vector contained centered keypoint coordinates and confidences (3*K* = 21 dimensions), computed as follows.

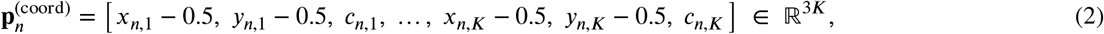

The second sub-vector contained pairwise intra-animal keypoint distances 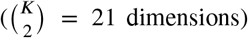, computed as follows.

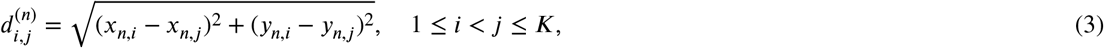

The third sub-vector contained frame-to-frame keypoint velocities (*K* = 7 dimensions), computed as follows.

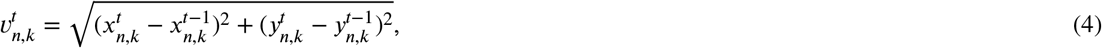

with 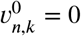 for the first frame of each window. Total: 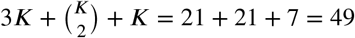 for *K*=7 keypoints.

#### Reliability scalars

Four scalar features were appended to each animal’s pose-self vector, extending the input to 53 dimensions. Here, 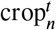 is the per-animal crop window for animal *n*, frame is the video frame extent, and 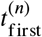 is the first frame in the current track for animal *n*:

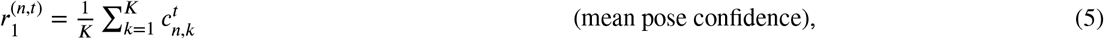

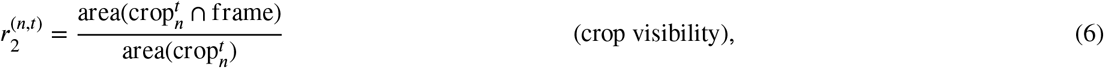

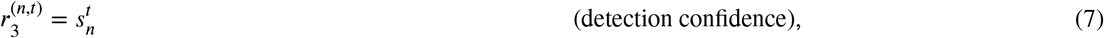

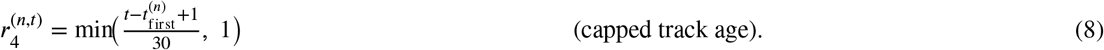

A binary pose mask 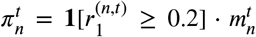 gated the pose-conditional relational features and the fusion bypass below.

#### Pairwise relational features (11 dimensions)

For each ordered animal pair (*i, j*) at frame *t*, an 11-dimensional vector 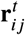 was computed. Features *f*_0_–*f*_5_ derived from bounding-box geometry were always available. In these equations, *ℓ*_*i*_ denotes body length as the bounding-box diagonal, IoU denotes intersection over union, 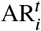 denotes bounding-box aspect ratio, and 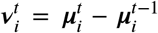 denotes finite-difference centroid velocity. Undotted 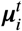 therefore always denotes centroid position:

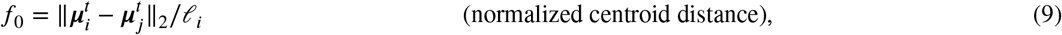

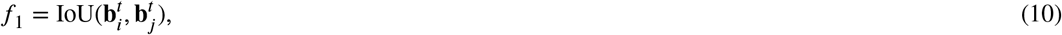

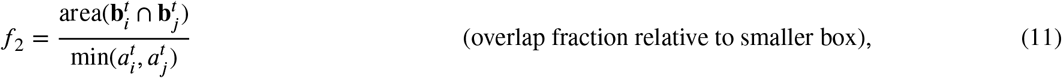

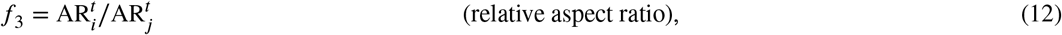

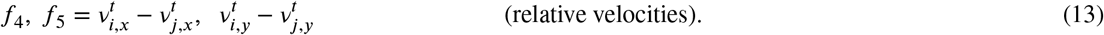

Features *f*_6_–*f*_10_, conditioned on both animals having valid pose 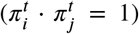, captured minimum inter-keypoint distance, undirected body-axis alignment, axis angular difference, signed approach velocity along the body axis, and a binary contact indicator (**1**[*f*_6_ < 0.35]); these were set to zero when either animal lacked valid pose. The relational encoder processed 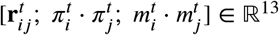 through a two-layer MLP with ReLU activations and dropout.

#### Gated-attention pose-visual fusion

When both pose and per-animal visual features were active, the model fused them via a learned gate. During approach, overlap, and contact, some keypoints become less reliable while animal-centered visual evidence becomes more informative, so a sample-dependent gate lets the model shift weight between pose embeddings and visual descriptors instead of assuming fixed stream weights (Arevalo, Solorio, Montes-y Gómez and González, 2017; Baltrušaitis, Ahuja and Morency, 2019). The pose-self vector (with reliability scalars) was projected to a *d*_*f*_ =128-dimensional embedding:

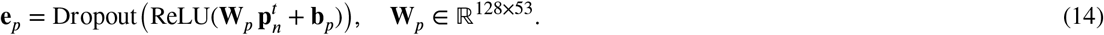

A sigmoid gating vector was computed from the concatenation of the visual descriptor and pose embedding:

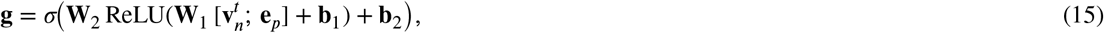

and the fused per-animal output was:

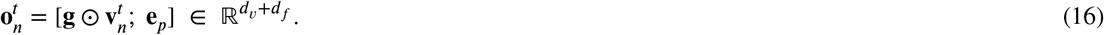

Here, **W**_*p*_, **W**_1_, **W**_2_, **b**_*p*_, **b**_1_, and **b**_2_ are learned parameters, *σ*(⋅) is an element-wise sigmoid, ⊙ is element-wise multiplication, and *d*_*v*_ is the visual descriptor dimension. When the pose mask indicated missing pose 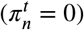, gates were forced to identity so the visual descriptor passed through unmodified. Cross-animal aggregation via masked mean pooling and the temporal-head architecture are detailed in the supplementary temporal sequence classifier section.

Pose and visual information were available together rather than as competing alternatives. During behavior training and held-out inference, each window contained pose-derived geometry and YOLO-pose-derived visual descriptors computed from the same source frames. When pose and per-animal visual features were both active, the model used gated-attention pose fusion with a 128-dimensional pose-fusion projection. We projected relation features through a learned relation pathway whose width matched the corresponding model-capacity setting. Fig 3 illustrates the four conceptual streams for a representative MARS frame.

### 2.5. MobileNetV3 control for reuse across backbones

The YOLO/SPPF feature definition above established the primary BehaviorScope-X stream specification. MobileNetV3-large (Howard, Sandler, Chu, Chen, Chen, Tan, Wang, Zhu, Pang, Vasudevan, Le and Adam, 2019) provided a structurally distinct native pose backbone for testing backbone-dependent reuse. The analysis evaluated whether the same downstream behavior decoders could use native visual descriptors from a second pose-trained backbone when the pose model, feature dimension, and checkpoint provenance were explicitly specified.

We trained the MobileNetV3 pose checkpoint for pose localization on MARS using a pose-only MobileNetV3-large architecture initialized from ImageNet weights, input size 640, ImageNet normalization, seven MARS mouse keypoints, and a decoupled anchor-free pose head. We sampled behavior-context frames from the MARS train and validation source splits, held out `test_1` and `test_2` from checkpoint selection, and selected the checkpoint by pose mAP50–95. The selected checkpoint had 2,971,952 parameters and reached box mAP50 of 0.970, box mAP50– 95 of 0.801, pose mAP50 of 0.907, pose mAP50–95 of 0.670, and PCK@0.05 of 0.950 on a validation set of 3,000 images.

For behavior reuse, we kept MobileNetV3 in its native form rather than modifying it to mimic the YOLO/SPPF architecture. We extracted visual descriptors from the final MobileNetV3 feature block and then passed them through adaptive average pooling. The MobileNetV3 tap yielded a 960-dimensional descriptor for each crop. With one group crop and two per-animal crops, the raw MobileNetV3 visual descriptor was 2,880 dimensions per frame. MobileNetV3 temporal sequence classifiers consumed the native 960-dimensional per-crop descriptors. Classical RF/XGBoost models used the same MobileNetV3 raw descriptors only after the controlled PCA compression described below. This kept the cross-backbone control native to MobileNetV3 while preserving the same downstream cache-and-classify contract.

### 2.6. DeepLabCut SuperAnimal HRNet-W32 top-down route

The DeepLabCut route used a pose checkpoint trained in a widely used DeepLabCut workflow as both a coordinate estimator and a behavior-relevant visual representation source. It complements MobileNetV3, which changes the pose backbone within a native BehaviorScope-X route, by changing the pose framework, inference style, backbone geometry, and feature-tap interface while preserving the downstream cache-and-classify contract. We evaluated DLC-HRNet as a native top-down route in which we generated the full-video pose cache, animal crops, pose-derived relation streams, and HRNet visual feature cache from the fine-tuned DeepLabCut detector and pose models before downstream behavior classification. We fine-tuned a DeepLabCut SuperAnimal TopViewMouse HRNet-W32 pose model on converted MARS pose labels using the YOLO-compatible pose-frame selection and source-video availability specified in the DLC frame manifest. The final DLC project contained 21,614 source frames sampled from attack, investigation, and mount contexts. MARS resident-intruder frames contain two animals, so this corresponded to 43,228 animal-level DLC pose examples. We used 41,066 training images and 2,162 testing images during DLC pose fine-tuning. The validation-selected pose checkpoint was the epoch-27 model, which reached DLC validation mAP 0.918. The detector was selected independently at epoch 4 by detector mAP50–95, reaching mAP50–95 0.710, mAP50 0.960, and mAP75 0.867. Detector fine-tuning was run as a separate stage so that the selected pose checkpoint remained unchanged.

For the native DLC top-down cache, the fine-tuned detector first generated animal boxes from each full-video frame and the fine-tuned HRNet pose model then estimated keypoints within those detections. The cache builder assigned two animal slots through box overlap with previous-frame detections, constructed one group crop and two animal-centered crops per frame, extracted the same pose-derived relation features used by the other BehaviorScope-X routes, and wrote the standard full-video window schema consumed by the downstream temporal classifier. The train/validation DLC top-down cache contained 30,493 windows. The held-out DLC manifest contained 28 MARS test videos with accepted human annotation files, with the original MARS split retained in the manifest. We used these videos for route-level metric summaries; the broader MARS held-out manifest contained 30 videos before this annotation-availability filter.

Visual features were extracted from the fine-tuned DLC HRNet backbone rather than from a YOLO SPPF layer. The adapter registered a forward hook on `PoseModel.backbone.model`, capturing the raw multi-branch TIMM HRNet output before DLC collapsed the branches for keypoint prediction. The default `semantic_concat` tap globally averaged and concatenated the four HRNet-W32 branches, yielding 32 + 64 + 128 + 256 = 480 visual features per crop. Strict tap validation required the hook to fire, the four branch widths to match the expected HRNet-W32 widths, and the concatenated feature dimension to equal 480. We trained the Attention-256 behavior classifier from DLC-derived pose and relation features plus these cached HRNet visual descriptors, then evaluated it on the held-out DLC top-down cache without rerunning classifier training.

### 2.7. Visual descriptor provenance analysis across pose routes

The cross-backbone and DLC-HRNet routes made descriptor provenance directly measurable before temporal classification. These analyses link the machine-learning question of representation accessibility to the ethological question of which visual evidence is preserved for behavior measurement. A visual-only linear probe estimates how much behavior-label information is linearly accessible from frozen descriptors before pose geometry, relation features, and temporal sequence modeling are added (Alain and Bengio, 2016). In biological terms, higher visual-probe performance indicates that the pose-trained visual space already separates behavioral contexts such as attack, investigation, mount, and other. Fisher ratios summarize class separation in the descriptor space, and linear centered-kernel alignment compares the geometry of descriptor spaces across pose routes (Kornblith, Norouzi, Lee and Hinton, 2019a). Video-identity probes use the same descriptor vectors to estimate recording-specific structure, which is useful for distinguishing behavior-accessible evidence from video-specific visual context.

The analysis used held-out MARS windows shared by the YOLO/SPPF, MobileNetV3, and DLC-HRNet caches. Each sample was represented by visual descriptors alone. For each 32-frame window, we concatenated the group descriptor and the two per-animal descriptors at each frame, then averaged across frames to obtain a window-level visual vector. We also analyzed group descriptors and paired-animal descriptors separately to determine whether behavior-accessible signal came primarily from scene context or from animal-centered crops. Balanced matched subsets prevented class imbalance from dominating the geometry estimates. Route-level Fisher ratios, effective dimensionality, PCA spectra, linear centered-kernel alignment, and visual-only behavior probes used 2,000 matched windows, with 500 windows per behavior class. Component and video-identity probes used 1,200 matched windows, with 300 windows per class. Behavior probes used grouped video-level folds so that windows from the same held-out video did not appear in both training and test folds. Feature channels were interpreted within route only because YOLO/SPPF, MobileNetV3, and HRNet channels are not homologous.

### 2.8. Full-video window construction and label assignment

Full-video training required labels that preserved the temporal structure of the recordings rather than only the centers of pre-trimmed behavior clips. We therefore designed the MARS full-video training set around the original human `.annot` files. For each source video, we first converted the human annotations into a per-frame label vector for attack, investigation, mount, and other. We then sampled fixed-length sliding windows across the entire video so that training examples included behavior interiors, behavior boundaries, and non-behavior context. Windows began at frame 0 and advanced in 16-frame steps; we retained a window only when its final frame remained inside the source video. The window construction preserved the continuous-video context needed for later ethogram evaluation.

For MARS full-video training, each 32-frame window was assigned a label from the per-frame human annotation vector. Within a window, we counted the number of frames assigned to attack, investigation, mount, and other. A window was kept only if a single class had the highest count and that class occupied at least 50% of the 32 frames. The retained window then received that class label. If two or more classes tied for the highest count, including exact boundary cases such as 16 frames from one class and 16 frames from another, the window was excluded from training rather than being forced into either label. Full videos contain many more other frames than annotated social-behavior frames; we therefore retained 30% of other-class windows after this label-assignment step when constructing the training set. Other-class subsampling affected the training distribution only; held-out evaluation still used full videos and frame-level ground truth. For held-out MARS evaluation, we retained all valid windows from `test_1` and `test_2`, including boundary and other-dominated windows, so the evaluation distribution reflected continuous video deployment rather than a balanced training sample.

For Fly-v-Fly, we used a separate short-bout-safe labeling rule. If any non-other Fly behavior occurred in a window, we used the highest-priority non-other behavior present in that window. The short-bout labeling rule preserved biologically meaningful positive windows for Fly aggression bouts that were too short for dominance-based labeling. We use the term priority-expanded frame labels for the corresponding per-frame Fly labels generated from the bout tables after resolving overlapping behavior annotations by the same fixed priority order (lunge, tussle, hold, charge, wing threat, then other). The priority rule kept Fly training and evaluation labels sensitive to brief aggressive events that would be diluted by majority-window labeling.

### 2.9. Temporal sequence classifiers and experimental matrix

We designed the MARS behavior classifier as a lightweight sequence-modeling platform for controlled comparisons rather than as an architecture search. The experimental matrix separated four downstream questions: which reused streams carried behavior signal, whether the same reused representation supported multiple temporal decoder families, whether decoder capacity changed held-out recovery, and whether the reuse strategy transferred to a different pose-native visual backbone. All comparable temporal sequence classifier runs used the same window manifests, class labels, optimizer settings, decoding policy, postprocessing rules, and held-out videos.

Each temporal sequence classifier received a sequence of per-frame multi-stream representations and produced a four-class prediction over attack, investigation, mount, and other. We evaluated two temporal heads. The LSTM head used a single-layer unidirectional LSTM and the final temporal output. The attention head used query-based four-head temporal attention pooling over the frame-level sequence, without an additional attention-projection bottleneck. Both heads used learned positional encoding where applicable, dropout 0.5, and a final linear classifier.

We evaluated three capacity settings, with hidden dimensions 256, 512, and 896. The hidden dimension set the temporal-head width and also widened the pose-self and pairwise-relation encoders. The visual descriptor dimension was fixed by the upstream pose backbone: 256 dimensions per crop for YOLO/SPPF and 960 dimensions per crop for MobileNetV3-native. The pose-visual fusion bottleneck was fixed at 128 dimensions. For the MARS and MobileNetV3 matrices, each temporal sequence classifier configuration was run with seeds 42, 43, and 44 to estimate stochastic training variability while holding data splits, feature caches, window definitions, and evaluation metrics fixed.

All MARS training and evaluation parameters were locked through the central configuration file and are summarized in Table 3. The pose backbones remained frozen during behavior-classifier training, so stream, capacity, and temporal-head effects reflect downstream feature processing and sequence decoding rather than changes to the pose-trained visual representation. We performed these experiments on Windows 10 with Python 3.11, 16 logical CPU cores, 31.9 GB RAM, and an NVIDIA GeForce RTX 2080 GPU with 8 GB VRAM. For each stage, we measured CPU and RAM use, GPU utilization, GPU memory, GPU power, elapsed time, and command status. We extracted classifier parameter counts from the run-specific configuration files written during training (Table 2).

**Table 2:** Temporal classifier parameter counts for anchor configurations. Counts were extracted from the trained classifier configuration files for the configurations that anchor the main experimental comparisons. Pose backbones and feature extractors were frozen for behavior-classifier training, so trainable counts correspond to the downstream classifier path. Additional stream and capacity variants are preserved in the configuration-level provenance table.

| Route | Configuration | Visual dim / crop | Total params | Trainable params |
| --- | --- | --- | --- | --- |
| MARS YOLO/SPPF | Full stream, attention-256 | 256 | 4.98M | 3.86M |
| MARS YOLO/SPPF | Full stream, LSTM-256 | 256 | 2.94M | 1.82M |
| MARS YOLO/SPPF | Pose and relations only, attention-256 | 0 | 1.40M | 1.40M |
| MARS YOLO/SPPF | Visual only, attention-256 | 256 | 3.27M | 2.15M |
| MARS YOLO/SPPF | Animal visual only, attention-256 | 256 | 2.09M | 0.97M |
| MARS MobileNetV3 | Full stream, attention-256 | 960 | 23.12M | 23.12M |
| MARS MobileNetV3 | Full stream, LSTM-256 | 960 | 4.48M | 4.48M |
| DLC-HRNet top-down | Full DLC route, attention-256 | 480 | 8.21M | 8.21M |
| Fly-v-Fly YOLO/SPPF | Full stream, attention-256, 8/16 frames | 256 | 4.97M | 3.86M |

**Table 3.**
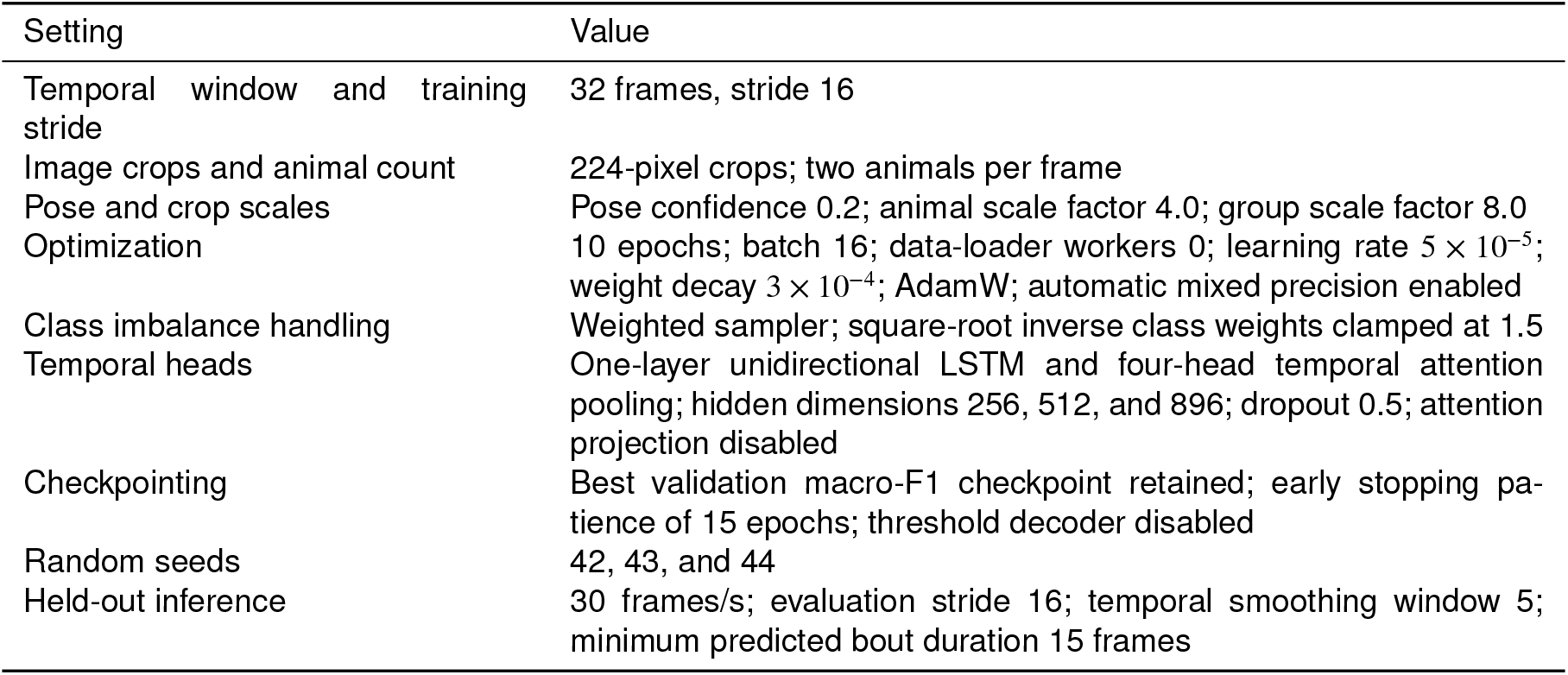
Fixed temporal sequence classifier training settings and MARS held-out evaluation settings. The central configuration defined these values before model training and held-out evaluation. Fly-v-Fly reused the optimizer, feature-caching, and decoder structure but used the species-specific temporal windows, class-balance setting, and postprocessing values reported below.

### 2.10. Training objective and validation selection

We trained temporal sequence classifiers with cross-entropy, with class-weighting specified by experiment. For a batch of *B* windows with logits ***z***_*b*_, label *y*_*b*_, and class weight 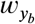, the loss for model parameters *θ* was

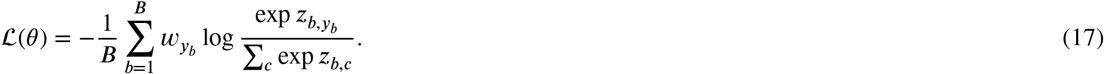

Here, *z*_*b,c*_ is the logit assigned to class *c* for window *b*. We computed class weights from training-window labels only using square-root inverse-frequency weighting. Let *n*_*c*_ denote the number of training windows assigned to class *c*. We rescaled weights relative to the minimum class weight, with an experiment-specific maximum clamp *w*_max_:

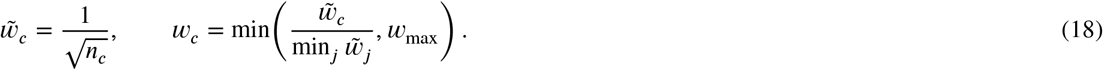

For MARS and MobileNetV3 behavior classifiers, *w*_max_ = 1.5 and the training loader used a weighted sampler based on the same class weights. For the focused Fly-v-Fly adaptation, *w*_max_ = 1.2 and the training loader used random sampling. The square-root weighting retained mild loss-level compensation for the short-bout Fly class imbalance while avoiding simultaneous sampler-level and loss-level upweighting of rare behaviors. MARS and MobileNetV3 temporal sequence classifiers used the fixed optimizer and data-loader settings in Table 3; Fly-v-Fly used the same optimizer, learning rate, weight decay, dropout, and learned positional encoding, with the batch size and early-stopping settings reported in Section 2.13.

Validation used the same argmax decoding rule as training-time inference:

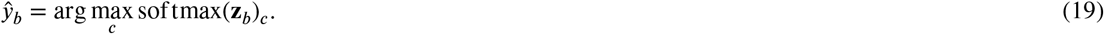

For each class *c*, we computed precision, recall, and F1 from the validation-window confusion counts:

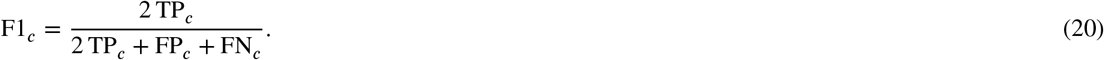

where TP_*c*_, FP_*c*_, and FN_*c*_ are class-specific true positives, false positives, and false negatives. We selected the checkpoint with the highest validation macro-F1 for each temporal sequence classifier run. Threshold decoders were disabled for the controlled model-family comparisons so that held-out performance did not depend on post hoc threshold tuning. Thus, model selection and held-out decoding used the same unthresholded decision rule across temporal-model comparisons.

### 2.11. MARS temporal sequence classifier stream, head-family, and capacity matrix

We designed the MARS stream-ablation matrix to test which information sources carried behavior signal under a common training and evaluation protocol. For the primary YOLO/SPPF route, we trained six stream conditions for each temporal head, capacity, and seed. One condition used only per-animal visual descriptors. A second used only pose-self and pairwise relational geometry. A third used the two visual streams without pose-derived geometry. Two additional ablations removed either the group visual descriptor or the pairwise relation features while retaining the remaining streams. The full-stream condition used group visual descriptors, per-animal visual descriptors, pose-self features, and pairwise relations. The stream matrix yielded 6 ×2 ×3 ×3 = 108 YOLO/SPPF MARS temporal sequence classifiers. Fig 4 summarizes the stream conditions and the factors held fixed across this matrix.

**Figure 4:**
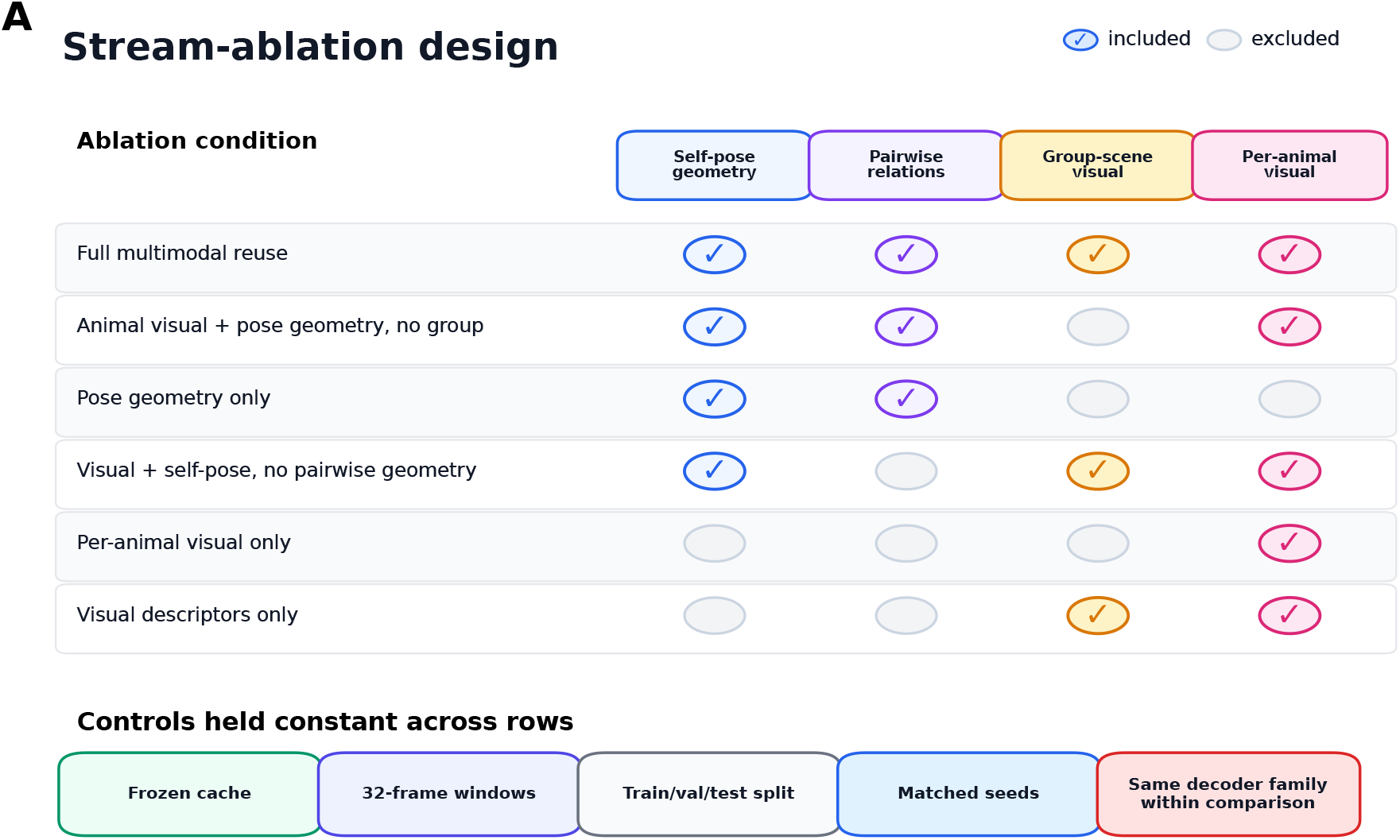
Stream-ablation design for the primary YOLO/SPPF temporal sequence-classifier matrix. Each row defines one stream condition by including or excluding self-pose geometry, pairwise relations, group-scene visual descriptors, and per-animal visual descriptors. Across rows, the frozen feature cache, 32-frame temporal windows, train/validation/test split, random seeds, and decoder family within a comparison were held fixed.

The MobileNetV3 cross-backbone analysis used the same full-stream input layout, the same two temporal heads, hidden dimension 256, and the same three seeds. We did not repeat the full stream-ablation or capacity matrix for MobileNetV3 because this experiment asked whether a second native pose backbone could support the full amortized pose vision workflow, not whether each MobileNetV3 stream had the same ablation profile as YOLO/SPPF.

The factorial design supports three within-condition comparisons. Holding the other axes fixed, the stream ablations isolate which reused streams matter, the 256/512/896 settings isolate the effect of decoder capacity, and the LSTM and attention heads isolate dependence on the sequence-modeling family.

### 2.12. Controlled Random Forest and XGBoost baselines

Pose-derived features are often paired with practical non-sequential classifiers in animal-behavior pipelines. We evaluated Random Forest (Breiman, 2001) and XGBoost (Chen and Guestrin, 2016) baselines to construct a negative control for static, non-sequential reuse of the same cached representation. The classical models used the same MARS train/validation window manifest and the same 30-video held-out evaluation manifest as the temporal sequence classifiers. However, they differed in how temporal context and visual information were represented. Instead of learning a temporal decoder over the 32-frame window, the classical models received explicit tabular feature vectors. In addition, frozen visual descriptors were compressed with PCA before tabular concatenation; otherwise, each sample would contain tens of thousands of visual values and the comparison would be dominated by high-dimensional tree fitting rather than controlled access to visual evidence.

The controlled classical matrix used four feature sets for each visual route. The *pose-frame* condition used only the 132-dimensional center-frame pose-and-relation vector. The *pose-window* condition flattened the full 32-frame pose- and-relation window, yielding 32 × 132 = 4,224 input features. The *pose-visual-frame-PCA* condition concatenated the center-frame 132-dimensional pose-and-relation vector with 64 PCA-compressed visual features, yielding 196 features per sample. The *pose-visual-window-PCA* condition flattened the full 32-frame pose-plus-visual sequence, yielding 32 × 196 = 6,272 features. Thus, the windowed classical baselines were provided the same 32-frame span used by the MARS temporal sequence classifiers, but as an explicit tabular concatenation rather than as a learned sequence.

For visual feature conditions, raw frozen visual descriptors were first standardized and compressed with incremental PCA to 64 dimensions per frame using batches of 8,192 training-frame samples. The scaler and PCA were fit on training frames only and then applied without refitting to validation and held-out data. The training-only transform prevented validation or held-out videos from influencing the visual basis while avoiding a high-dimensional tree-model comparison in which raw visual windows would contain 24,576 YOLO/SPPF visual values (32 × 768) or 92,160 MobileNetV3 visual values (32 × 2,880) per sample before pose features. PCA was necessary to keep the RF/XGBoost comparison focused on whether non-sequential tabular models could use frozen visual evidence, rather than on memory demand and split-search behavior in extremely high-dimensional raw visual windows. The same 64-dimensional PCA target was used for YOLO/SPPF and MobileNetV3 so that the classical visual-feature representation was controlled across backbones.

We implemented the Random Forest baseline in scikit-learn (Pedregosa, Varoquaux, Gramfort, Michel, Thirion, Grisel, Blondel, Prettenhofer, Weiss, Dubourg, Vanderplas, Passos, Cournapeau, Brucher, Perrot and Duchesnay, 2011) with 500 trees, 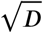 candidate features per split, balanced inverse-frequency class weighting, no maximum-depth limit, and a minimum of five samples per leaf, using all available CPU workers. The XGBoost baseline used gradient-boosted trees with a multiclass soft-probability objective, 1,000 estimators, maximum depth 8, learning rate 0.1, row and column subsampling rates of 0.8, validation multi-class log-loss as the evaluation metric, and early stopping after 50 rounds without validation improvement, also using all available CPU workers. We trained each classical configuration with seeds 42, 43, and 44.

Held-out evaluation used the same 30-video MARS test set, overlapping-window probability averaging, median-filter smoothing (*k*=5), minimum predicted bout duration of 15 frames, and the same frame-level and bout-level metrics as the temporal sequence classifiers. Because static tabular concatenation does not preserve the temporal order and stream structure available to the LSTM and attention heads, these baselines function as a negative control for non-sequential reuse.

### 2.13. Fly-v-Fly short-bout stress test

The Fly-v-Fly experiment evaluated whether the same pose-to-behavior design could be adapted to a different species, keypoint geometry, behavior repertoire, and bout-duration scale. The primary question was whether short-bout Fly aggression could be decoded after reusing a Fly YOLO-pose checkpoint as the frozen representation source. The Fly-v-Fly classifier capacity was chosen from the MARS temporal sequence classifier design while adapting the framework to shorter behavior bouts: we used a focused attention-256 decoder rather than repeating the full MARS head and capacity matrix. A secondary question was how model interpretation changed when the available second annotation layer was used as a human-reference comparison. We did not extend the RF/XGBoost matrix to Fly-v-Fly because it would conflate this short-bout adaptation test with an underpowered tabular baseline comparison. The held-out Fly-v-Fly set contains five test videos, many target events are shorter than or comparable to the classifier windows, and secondary annotations are available locally for only one video. The Fly-v-Fly analysis focused on temporal sequence classifier adaptation, event recovery, and annotation-aware interpretation.

Fly-v-Fly aggression (Eyjolfsdottir et al., 2014) provided an adaptation setting with short, imbalanced aggression behaviors performed by much smaller animals than the MARS resident-intruder mice. For Fly-v-Fly, we used a separate Fly YOLO-pose checkpoint, the full Fly feature set, two animals per frame, crop size 224, pose-confidence threshold 0.2, and a single downstream hidden dimension of 256 for the behavior decoder. Similar to the MARS pose-training dataset, we trained the Fly pose checkpoint as a YOLO26n-pose nano model using the Ultralytics framework (Ultralytics, 2023) on a Fly-v-Fly pose dataset. The Fly pose-training set contained 23,066 sampled frames across movies 1–10, split into 18,452 training frames and 4,614 validation frames. The behavior-context distribution of these pose-training frames was 6,000 lunge, 6,000 wing-threat, 3,003 tussle, 1,894 hold, 169 charge, and 6,000 other frames.

Movies 6–10 were later used as the behavior-label held-out set, so the Fly pose checkpoint had already seen frames from those movies during keypoint-localization training. For Fly behavior results, the held-out unit was the behavior label. The frozen backbone supplied localization and feature extraction for these videos, while behavior labels from movies 6–10 were excluded from behavior-model training and checkpoint selection. Since the public Fly-v-Fly subset contains a limited number of videos, this analysis is interpreted as a behavior-label and temporal-scale stress test using a frozen assay-trained pose checkpoint, rather than as a strict unseen-video pose-generalization benchmark. The Fly pose-checkpoint training and validation curves are shown in Fig 5.

**Figure 5:**
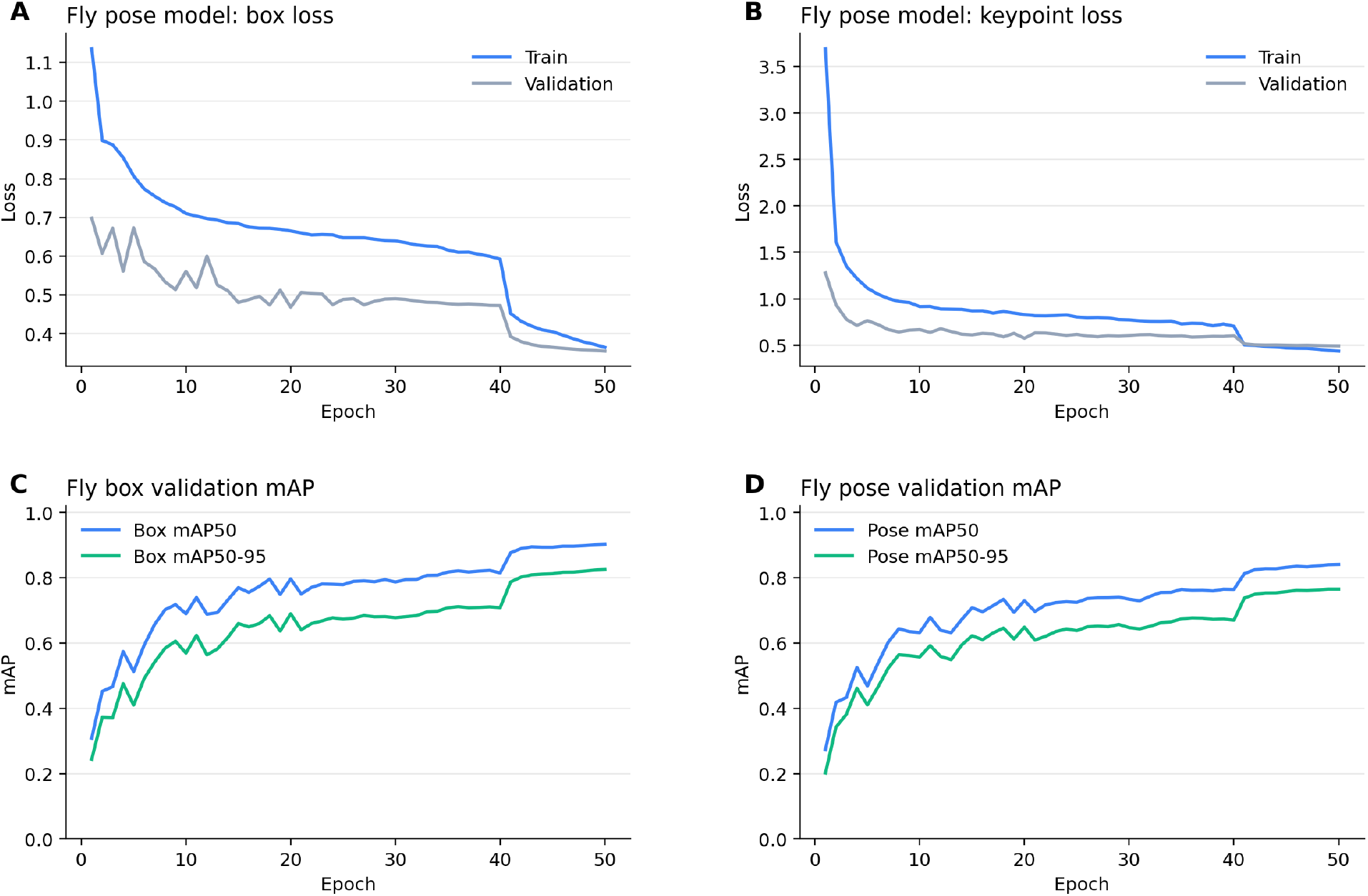
Fly-v-Fly pose-checkpoint convergence over 50 training epochs. (A) Box localization loss and (B) keypoint pose loss both decreased for training and validation sets. (C) Box detection mAP50 reached 0.903 and mAP50–95 reached 0.826. (D) Pose validation mAP50 reached 0.841 and mAP50–95 reached 0.765. Although final Fly pose accuracy was lower than the MARS checkpoint (pose mAP50–95 of 0.765 versus 0.899), the curves confirm that the checkpoint had converged before downstream behavior classification.

Temporal scale was treated as the main adaptation variable. We evaluated two short-window configurations, w16/s8 (16-frame windows, stride 8) and w8/s4 (8-frame windows, stride 4). The short-window settings were chosen because several Fly-v-Fly behaviors are much shorter than the MARS mouse social behaviors. In the held-out movies, charge had a median ground-truth duration of one frame and lunge had a median duration of four frames (Table 4). Both Fly configurations used the same optimizer, learning rate, weight decay, dropout, pose-fusion bottleneck, and learned positional encoding as the MARS temporal sequence classifier runs (Table 3). The remaining training settings were adapted to the Fly-v-Fly regime. We trained the attention-256 decoder with batch size 64, a maximum duration of 40 epochs, and early-stopping patience 20. To account for the smaller and more imbalanced Fly training set, Fly-v-Fly used square-root inverse class weights clamped at 1.2 in the loss and random sampling in the training loader. The behavior decoder used validation movies 3 and 5 for checkpoint selection and a single prespecified model seed, 42. Thus, the focused Fly adaptation contained two temporal sequence classifier runs: attention-256 with w16/s8 and attention-256 with w8/s4. We used this narrowed design because the Fly-v-Fly experiment tested adaptation to a different species and bout-duration regime, whereas the MARS experiments above provided the downstream head, capacity, and seed-variability analysis.

**Table 4.** Fly-v-Fly ground-truth behavior-bout distribution after priority-label expansion. The Fly-v-Fly aggression dataset contained 10 movies. Held-out evaluation used movies 6–10. Median durations are reported in frames at 30 frames/s for the same priority-expanded frame labels used by the Fly training and evaluation code.

| Behavior | All movies | Median | Held-out | Median |
| --- | --- | --- | --- | --- |
| Lunge | 3061 | 4 | 1590 | 4 |
| Wing threat | 291 | 13 | 154 | 13 |
| Charge | 114 | 1 | 57 | 1 |
| Hold | 131 | 7 | 50 | 12 |
| Tussle | 261 | 7 | 155 | 8 |

To avoid erasing brief positive events through majority labeling over a window, Fly training windows used a short-bout-safe priority-OR rule. If any non-other aggressive behavior occurred within the window, the window was assigned the highest-priority non-other class present.

Fly held-out evaluation followed the same amortized pose vision principle as the MARS held-out analysis. We built held-out window manifests separately from movies 6–10 and retained all valid evaluation windows. Since Fly bouts are short, the held-out evaluation stride was 4 frames for both temporal-window lengths, yielding dense w16/s4 and w8/s4 held-out window sets. Frozen YOLO/SPPF features were precomputed once for each held-out window length and then reused by the corresponding trained Fly temporal classifier. Thus, the Fly comparison varied temporal-window length while holding the Fly pose checkpoint, feature tap, held-out windows within each length, and visual-feature cache fixed. We used temporal smoothing window 3 and minimum bout duration 3 frames, reflecting the shorter duration of Fly-v-Fly aggression bouts.

### 2.14. Fly-v-Fly annotation-aware evaluation

We evaluated Fly-v-Fly aggression with primary-label metrics and, where available, secondary-annotation sensitivity analyses. Before defining the final Fly-v-Fly evaluation modes, we audited the available primary and secondary annotation files for movie 6 (54,000 video frames). Overall frame agreement was 94.1%, but this value was dominated by the other class; among frames assigned a non-other behavior by the primary annotator, same-label agreement was 55.0%. If non-other frames were defined as frames where either annotator marked behavior, same-label agreement was 49.1%, whereas frames marked as behavior by both annotators had 82.0% same-label agreement. The disagreement was also behavior-specific. Conditioning on the primary annotation, wing threat showed 95.5% inter-annotator agreement, whereas lunge showed 18.5% agreement, with 69.1% of primary-labeled lunge frames labeled as other by the secondary annotator. Hold showed 16.7% agreement as hold and 81.2% confusion with tussle. The single-movie audit defines the available human-human comparison and motivates reporting model performance under multiple ground-truth assumptions.

For Fly-v-Fly videos with secondary annotations, the evaluator constructs five ground-truth modes. The *primary* and *secondary* modes evaluate against each annotator separately. The *union* mode treats a behavior frame as present if it appears in either annotation file, which captures a more permissive estimate of behavior occurrence before bout metrics are computed from the resulting frame sequence. The *intersection* mode retains only frames where both annotators agree on the same non-other behavior and assigns disagreement frames to other, thereby testing performance on the most conservative consensus subset. The *padded-primary* mode expands each primary bout by an empirical boundary margin estimated from primary-versus-secondary start and end differences. Let *s*_*A*_, *e*_*A*_ and *s*_*B*_, *e*_*B*_ be the start and end boundaries of a matched same-class primary and secondary bout pair. The standard deviation was computed across all such start- and end-boundary differences, and each primary bout [*s, e*] was then padded as follows:

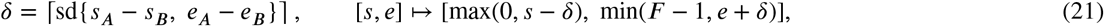

where *F* is the number of video frames. In the available movie 6 secondary-annotation audit, this procedure matched 209 same-class bout pairs and yielded a 21-frame padding margin. Human-human agreement is computed by treating one annotator as the prediction and the other as the reference under the same frame-level and bout-level metrics used for model evaluation. Secondary, union, and intersection modes are reported only for videos where secondary annotations are available; padded-primary evaluation uses the empirical boundary margin estimated from those available secondary annotations.

### 2.15. Prediction postprocessing

Postprocessing was required to convert overlapping window predictions into a single continuous ethogram while limiting isolated one-window fluctuations that would be implausible as biological bouts. For MARS held-out inference (32-frame windows, stride 16, 30 frames/s), we merged overlapping windows into frame-level predictions, applied a five-frame temporal smoothing window, and removed predicted behavior bouts shorter than 15 frames (0.5 s at 30 frames/s). We selected these values *a priori* based on biological plausibility for MARS mouse social behaviors and were not tuned on held-out performance. The 15-frame minimum reflects the approximate lower bound on biologically meaningful social-behavior episodes at 30 frames/s, and the five-frame smoothing window was chosen to suppress isolated single-window fluctuations without erasing genuine short bouts. We kept raw and smoothed metrics separately so that the contribution of postprocessing could be assessed. For the native YOLO/SPPF and MobileNetV3 routes this contribution was negligible (per-video frame macro-F1 changes below 0.002), so their reported held-out metrics are effectively unchanged by smoothing; for the DLC-HRNet route, smoothing reduced held-out performance, and raw-window outputs are therefore reported as the primary readout (Section 3.6). Formally, overlapping-window predictions were merged by averaging per-frame softmax vectors:

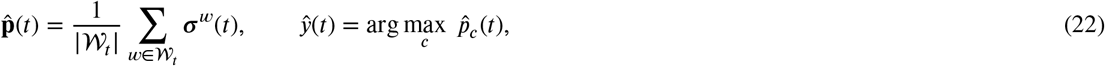

where *W*_*t*_ is the set of windows covering frame *t* and ***σ***^*w*^(*t*) is the softmax vector from window *w* at frame *t*. A one-dimensional median filter of width *W*_med_ (*W*_med_=5 for MARS, 3 for Fly) was then applied to the integer label sequence. Finally, contiguous same-class runs shorter than a minimum duration *D*_min_ were iteratively reassigned to the longer adjacent run, with the preceding run used to break ties, until no run remained below the minimum. For MARS, *D*_min_ = 15 frames (0.5 s at 30 fps); for Fly, *D*_min_ = 3 frames. Raw and postprocessed outputs were retained so that postprocessing effects could be separated from decoder performance during held-out analysis.

Fly evaluation used 30 frames/s, evaluation stride 4, temporal smoothing window 3, minimum bout duration 3 frames, and temporal IoU thresholds 0.10, 0.25, and 0.50. We used shorter smoothing and minimum-duration settings for Fly-v-Fly to reflect the shorter time scale of fly aggression bouts. Since some Fly-v-Fly annotations, particularly charge and lunge, occur at the one-to-four-frame scale and show substantial annotator uncertainty, raw and postprocessed Fly predictions were both retained. Raw predictions quantify the decoder’s sensitivity to short bouts before temporal cleanup, whereas the output after smoothing and minimum-duration filtering provides a conservative continuous ethogram that suppresses isolated single-frame fluctuations. The rationale for these species-specific choices is further discussed in the Supplementary Material.

### 2.16. Evaluation metrics and statistical units

We evaluated the system at four levels: pose localization, frame-wise behavior classification, bout recovery, and ethogram-level summary structure. The evaluation levels were kept separate, as they answer different biological and computational questions. Formal metric definitions, statistical testing procedures, and ethology-level summary statistics are detailed in Supplementary Sections S5–S7.

For pose localization, we used object-keypoint similarity (OKS), keypoint root-mean-square error in pixels, and percentage of correct keypoints (PCK). PCK@0.05, PCK@0.10, and PCK@0.20 report the fraction of matched keypoints whose localization error falls within 5%, 10%, or 20% of the corresponding animal body-length scale. The pose metrics were used to establish whether the pose checkpoint trained for the assay provided a credible geometric basis for behavior classification; they were not used to select the temporal behavior classifier.

Frame-level behavior metrics treat each video frame as the observational unit. For each behavior class, precision is the fraction of frames predicted as that class that were correct, recall is the fraction of ground-truth frames from that class that were recovered, and F1 is the harmonic mean of precision and recall. Frame macro-F1 is the unweighted mean of the per-class frame F1 scores. This matters for full-video recordings because most frames are the background other class. Overall accuracy is dominated by that majority class and can look high even when the social behaviors are missed, whereas macro-F1 weights attack, investigation, mount, and other equally and therefore reflects whether the rare social behaviors are actually recovered. Class-specific frame F1 values are reported separately when the class identity of the error matters.

Bout-level metrics treat each contiguous behavior episode as the observational unit. We defined a bout as a maximal contiguous run of frames assigned to the same non-other behavior. A predicted bout was eligible to match a manual-annotation bout of the same class when their temporal intersection-over-union (TIoU) exceeded a specified threshold. Temporal IoU is the number of overlapping frames divided by the number of frames in the union of the predicted and ground-truth bout intervals. Bout precision, recall, and F1 were then computed from matched bouts, unmatched predicted bouts, and unmatched ground-truth bouts. Following the segmental F1 evaluation protocol established in temporal action segmentation (Lea, Flynn, Vidal, Reiter and Hager, 2017; Abu Farha and Gall, 2019; Ding, Sener and Yao, 2023), MARS summary tables report bout F1 at TIoU 0.10, 0.25, and 0.50. The lower threshold is reported as a sensitivity measure for short bouts, whereas 0.25 and 0.50 retain stricter temporal-boundary criteria. We report frame macro-F1 and bout macro-F1 separately, since a model can classify many frames correctly while fragmenting a single biological episode into multiple predicted bouts. These separable readouts also provide the basis for the diagnostic taxonomy used in the Discussion, where identity errors, boundary displacement, bout fragmentation, and bout merging are interpreted as distinct evaluation outcomes.

For ethogram-level summaries, we compared behavior budgets, bout counts, bout-duration sensitivity, bout-sequence structure, and transition probabilities. Behavior budget is the total time assigned to each behavior within a video. Bout count is the number of non-other bouts per behavior. Transition probabilities were computed from the sequence of non-other bouts after collapsing contiguous frames of the same class, so they summarize the order of behavioral episodes rather than individual frame-to-frame persistence.

Training, validation, and held-out evaluation were treated as distinct statistical contexts. Validation metrics were used for model selection and to describe training behavior, whereas held-out metrics were reserved for final performance comparisons. For paired model comparisons, the held-out video was the statistical unit. Paired deltas were computed per video, and bootstrap confidence intervals (Efron and Tibshirani, 1993) were sampled over videos so that repeated frames or bouts from the same recording were not treated as independent experiments. We report seed summaries as mean and standard deviation across seeds, with individual seed values retained for rank-stability assessment.

### 2.17. Ethogram-structure and bout-resolution analyses

The biological output of the system is a continuous ethogram rather than only a frame-level label sequence. Additional analyses separated three related quantities: whether behavior bouts were detected, whether their temporal boundaries were recovered, and whether the order of behavioral episodes was preserved. Unless otherwise specified, these analyses were computed per held-out video and then summarized across videos.

First, we audited bout under-segmentation and postprocessing effects. For each behavior class *c* and model *m*, we computed the predicted-to-human bout-count ratio,

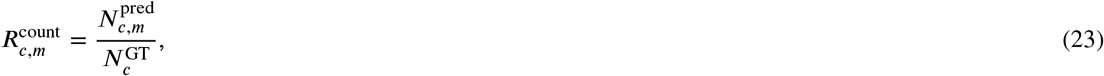

and the median-duration ratio,

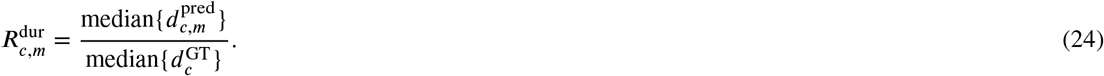

Here, 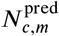 and 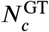 are the predicted and ground-truth bout counts for class *c*, and 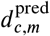 and 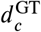 are the corresponding individual bout durations. We also quantified whether individual predicted bouts merged multiple ground-truth bouts of the same class by counting the number of same-class manual-annotation bouts overlapped by each predicted bout. The bout measurements were computed for both raw overlapping-window predictions and for ethograms after smoothing and minimum-duration filtering. The raw-versus-smoothed comparison was used to distinguish temporal compression introduced by the classifier from temporal compression introduced by postprocessing.

Second, we estimated the operating envelope of held-out bout recovery as a function of ground-truth bout duration. A manual-annotation bout was considered recovered when a same-class predicted bout matched it at the specified TIoU threshold. Recall was then summarized within duration bins < 0.25, 0.25–0.5, 0.5–1, 1–2, 2–4, 4–8, and ≥ 8 seconds at 30 frames/s. Duration bins containing fewer than 10 ground-truth bouts were not used for summary interpretation. The operating-envelope plots report both raw and postprocessed outputs and compare matched-capacity temporal heads so that duration sensitivity is not confounded with model capacity.

Third, we evaluated whether the predicted ethograms preserved behavioral sequence structure beyond behavior base rates. For each full video, the ethogram was converted to a sequence of non-other bouts after collapsing contiguous same-class episodes. For an *n*-gram length *n*, let *G*_*n*_(*g*) and *P*_*n*_(*g*) denote the ground-truth and predicted counts of motif *g*. Multiset *n*-gram precision and recall were computed as

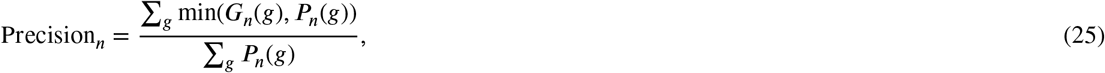

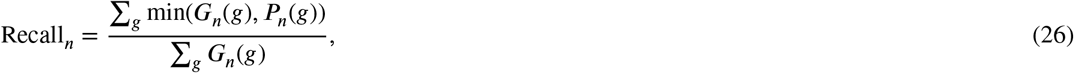

with *F* 1_*n*_ defined as their harmonic mean. We report *n* = 1–4 to span single-bout composition through short behavioral motifs. Since weighted *n*-gram scores can be dominated by common behaviors, we also stratified ground-truth 3-gram motifs by corpus frequency and reported recall for rare, intermediate, and common motif groups.

Two null sequence models were included to anchor the *n*-gram analysis. The composition-shuffle null permuted the ground-truth bout labels within each video, preserving the video-specific behavior composition and sequence length while destroying temporal order. The Markov null generated sequences from corpus-level first-order transition probabilities while preserving each video’s non-other sequence length. The sequence nulls tested whether the model recovered ethological sequence organization beyond unigram composition or a simple first-order transition process. Transition-probability matrices were retained as diagnostics, but they were interpreted together with the under-segmentation audit because row-normalized transition matrices can be distorted when a model merges many short bouts into fewer long bouts.

### 2.18. End-to-end deployment throughput benchmark

The final analysis measured whether the same decomposition used for controlled training and held-out evaluation remained practical when run as a complete video-to-ethogram deployment path. The benchmark used the selected full-stream LSTM-256 and attention-256 temporal sequence classifiers and three held-out MARS MP4 recordings. Hardware was treated as the measured variable rather than an ancillary factor, because the goal was to estimate deployment throughput under practical laboratory conditions.

Each benchmark run included the stages required during standard inference: video loading, YOLO-pose tracking, crop construction, frozen YOLO/SPPF feature extraction, temporal decoding, postprocessing, CSV output, and runtime measurement. We computed aggregate throughput as total processed frames divided by total wall-clock time. We also measured interval throughput after startup so that steady-state performance could be separated from launch and initialization overhead. Stage-timed profiling runs decomposed interval wall time into video handling, pose and crop preparation, frozen feature encoding, window assembly, and temporal classification.

The primary deployment benchmark used the standard OpenCV/Ultralytics inference route. A secondary RTX4080 profiling run used FFmpeg predecode/cache to test whether video preparation was the dominant discrete-GPU bottleneck. These measurements were not used for model selection. They were included to determine whether amortized pose vision could support near-real-time full-video behavior classification under practical laboratory conditions, and to identify which parts of the deployed path should be optimized without changing the scientific comparison between frozen pose-trained descriptors and downstream temporal decoding.

### 2.19. Open-source annotation and analysis application

For use beyond controlled benchmarks, we built BehaviorScope-X as an open-source Qt (PySide6) desktop application for laboratories whose primary expertise is behavioral biology rather than software engineering. The application organizes annotation, dataset preparation, feature caching, training, evaluation, and output inspection into a shared graphical workflow (supplementary GUI workflow section). Its central deployment design is flexibility across pose-model families: users first annotate videos and assign train, validation, and held-out splits, then choose the model-family tab corresponding to the assay’s pose model. The supported tabs are YOLO-pose, MobileNetV3, and DeepLabCut-HRNet. Each route has route-specific pose inference and feature-tap logic, but all routes emit compatible full-video sequence manifests, NPZ windows, pose-derived geometry streams, visual feature caches, classifier inputs, validation summaries, held-out evaluation tables, confusion matrices, bout summaries, and ethogram exports. The shared artifact contract makes the GUI routes comparable despite route-specific pose inference and feature taps.

The annotation interface exports only approved spans and supports both class-folder clips for legacy clip-based workflows and full-video BENTO annotation files with a source manifest recording video paths, class names, and split assignments. The YOLO-pose workflow builds full-video windows directly from an Ultralytics-compatible pose checkpoint, extracts frozen late-backbone visual descriptors, trains the temporal classifier, and supports single-video or folder-level prediction. The MobileNetV3 workflow uses a native MobileNetV3 pose checkpoint to build the same window-cache schema before extracting MobileNetV3 visual descriptors and training the same downstream temporal classifier. The DeepLabCut-HRNet workflow wraps DLC SuperAnimal pose/detector fine-tuning, top-down cache construction, HRNet feature extraction, classifier training, and held-out evaluation. Together, the tabs provide a common workflow surface for the three pose-model families evaluated here, and we provide the underlying scripts for automated or high-throughput execution.

Each tab constructs the same CLI command accepted by the underlying script, so every GUI-initiated run is reproducible from the command line. After YOLO-pose training, the application can export a bundled deployment checkpoint that embeds both the classifier checkpoint and the YOLO-pose weights, allowing inference from one deployable model package. For MobileNetV3 and DeepLabCut-HRNet, the application preserves the corresponding pose-checkpoint provenance, project-level metadata, and feature-cache manifests so evaluation remains reproducible even when inference is run through staged scripts rather than a bundled YOLO-style model. Held-out evaluation and bout-level metrics can also be run through companion command-line scripts included in the repository. The shared backend keeps the graphical workflow aligned with the scripted reproduction path used for the manuscript analyses.

## 3. Results

The Results first establish that cached pose-trained representations provide behavior-discriminative evidence in held-out MARS videos. We validate the pose checkpoint, measure full-video decoding, and use stream ablations and classical controls to identify the evidence carried by visual and geometric streams. MobileNetV3 and DeepLabCut-HRNet then define how reuse changes across pose backbones and pose-training ecosystems, and descriptor-provenance probes identify how the cached visual spaces differ before temporal decoding. Bout-compression and Fly-v-Fly analyses define temporal limits, and the final deployment section treats throughput and GUI/scripted reproducibility as operational consequences of the validated cache-and-classify decomposition. Table 5 summarizes the scientific role of each route and makes the evidentiary hierarchy explicit.

**Table 5.**
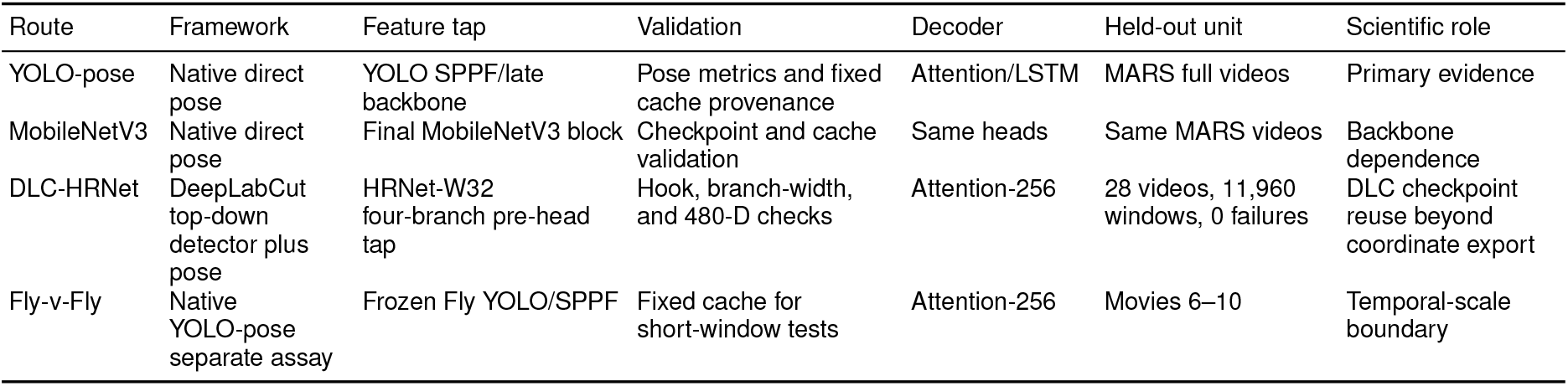
Pose-framework routes used to test amortized pose vision. The DLC-HRNet route uses a widely used DeepLabCut top-down workflow to produce both coordinate outputs and reusable pre-head visual descriptors for behavior decoding.

### 3.1. A YOLO-pose checkpoint trained on MARS provides the reusable representation substrate

BehaviorScope-X first requires that the upstream pose model provide a reliable assay-adapted representation. The analysis began by evaluating the primary YOLO-pose checkpoint before using its intermediate features for behavior decoding. The checkpoint was the YOLO-pose nano model used for the MARS resident-intruder experiments. Its frozen visual descriptors were extracted through the SPPF-containing feature tap `layers[0:10]` and adaptive average pooling, yielding 256 dimensions per crop. For each frame, the downstream behavior model received a 768-dimensional raw visual representation before any stream ablation: one group-scene descriptor and two animal-centered descriptors. The visual representation was paired with a 132-dimensional pose-derived geometry vector. Thus, the first result establishes the fixed pose-trained measurement substrate on which the downstream behavior analyses depend.

The pose checkpoint converged smoothly across box and keypoint objectives, and held-out keypoint recovery was high enough to support the downstream reuse experiments (Fig. 6). Across the seven MARS keypoints, PCK@0.10 ranged from approximately 0.79 to 0.90 and PCK@0.20 from approximately 0.91 to 0.97, with a mean matched-frame OKS of 0.642. The tail base and posterior hip landmarks were the least accurate under the strictest thresholds, whereas the neck and anterior landmarks were more stable. The validation pattern is consistent with the visual difficulty of rear-body landmarks during contact and occlusion, but it also provides a necessary control for the reuse experiments: the behavior classifiers below are evaluated on a validated pose source. They test whether a pose-trained checkpoint that already solves keypoint localization in this assay also contains reusable visual and geometric evidence for behavior classification.

**Figure 6:**
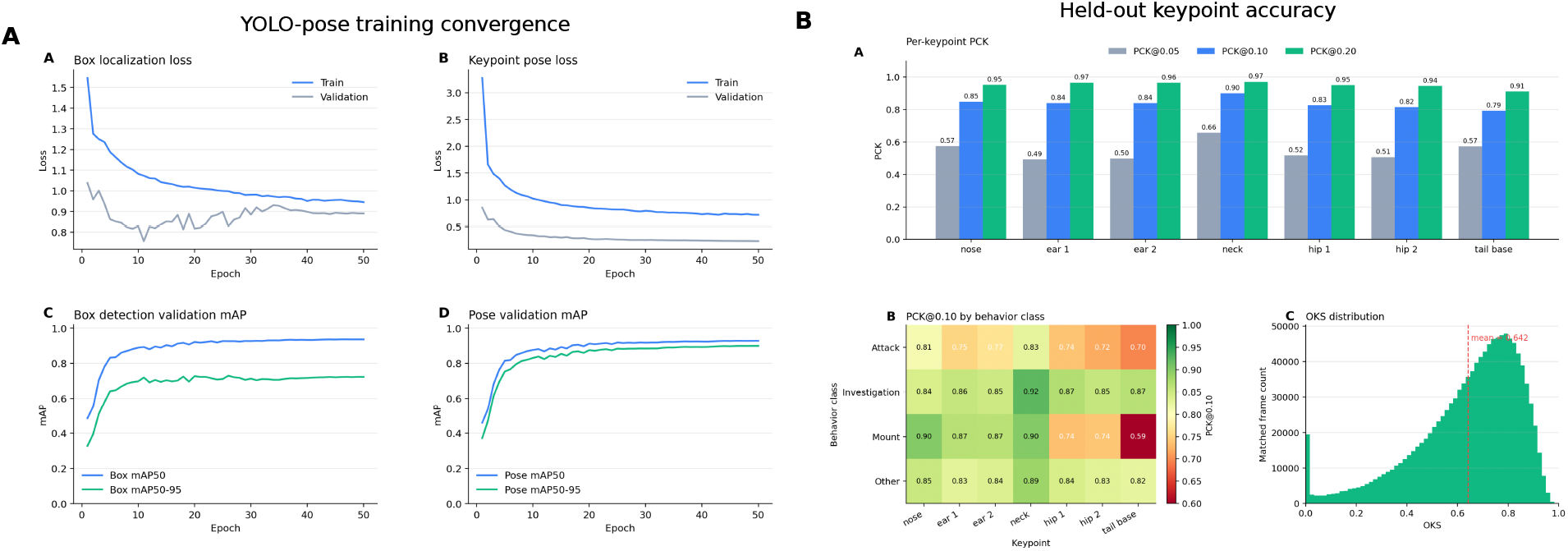
YOLO-pose quality and held-out keypoint accuracy. (A) Training and validation curves for box localization, keypoint pose loss, box mAP, and pose mAP show stable convergence of the pose checkpoint used as the primary BehaviorScope-X representation source. (B) Held-out keypoint accuracy summarizes per-keypoint PCK, behavior class-conditioned PCK@0.10, and the matched-frame OKS distribution. The figure documents the keypoint geometry and SPPF visual feature source before downstream behavior classifiers are evaluated.

### 3.2. Reusable YOLO/SPPF features support behavior decoding in full videos with recurrent and attention-based temporal heads

The frozen YOLO/SPPF representation supported full-video behavior decoding across temporal head families and capacity settings. To test this under controlled conditions, we evaluated a temporal sequence classifier matrix in held-out MARS resident-intruder recordings (Segalin et al., 2021c), with six stream conditions, two temporal heads (LSTM and temporal attention pooling), three hidden dimensions (256, 512, and 896), and three random seeds. The held-out manifest contained 30 videos; frame and bout metrics were computed on the 28 videos with accepted human annotation files. Model selection used validation macro-F1, but all primary results below are reported on the held-out videos. Training dynamics across the controlled matrix are summarized in Fig 7.

**Figure 7:**
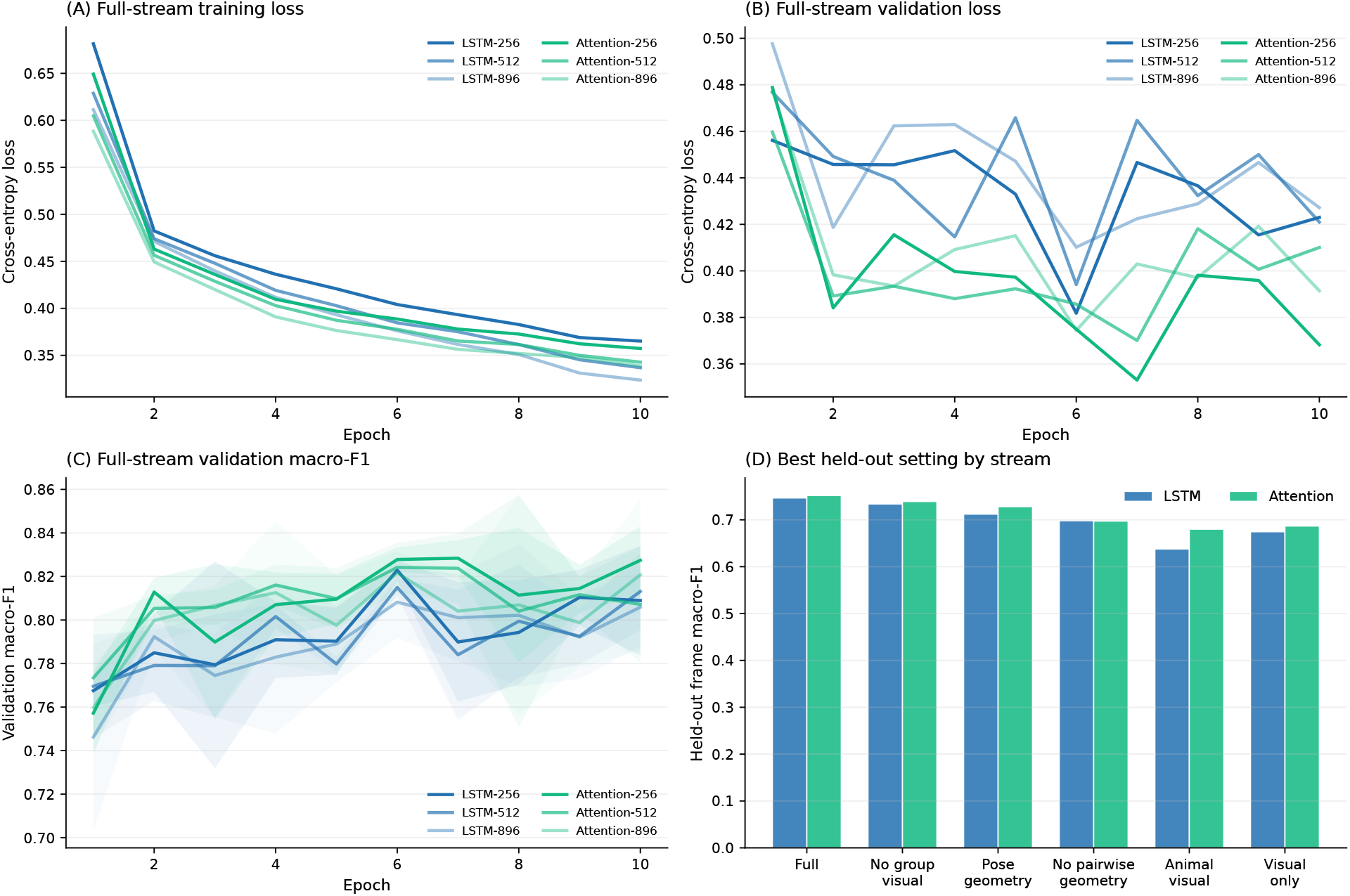
Training stability of YOLO/SPPF temporal sequence classifiers. Training loss, validation loss, validation recall, and validation macro-F1 are shown across stream conditions, head families, capacities, and seeds. The validation curves provide the validation-selection context for the held-out comparisons reported below.

The cached representation also made the controlled matrix computationally tractable after the frozen descriptors had been built. Across the 108 completed YOLO/SPPF temporal sequence classifier runs in the MARS matrix, each run trained for 10 epochs on the same 20,414 training windows and 10,079 validation windows. Median classifier-training wall time was 32.2 minutes per run (range, 25.3–58.5 minutes). One visual-only LSTM-256 run produced the 58.5-minute upper bound, during which epochs 6 and 7 had unusually low GPU utilization. The overall range reflects run-level system or I/O variability as well as model configuration. The full-stream runs had a similar median of 32.4 minutes (range, 31.5–39.0 minutes; Supplementary training-throughput figure). Thus, the stream ablations and temporal-head comparisons were not separate end-to-end video-training jobs; they reused a fixed representation cache and primarily retrained compact temporal decoders.

The best held-out configuration was full multimodal reuse with the attention-256 head, which reached a frame macro-F1 of 0.752 ± 0.006 and a bout macro-F1 of 0.680 ± 0.011 at TIoU 0.25 across seeds. The best LSTM configuration was the matched full-stream LSTM-256 model, with frame macro-F1 0.747 ± 0.006 and bout macro-F1@0.25 0.672 ± 0.002. Larger hidden dimensions did not improve held-out performance for the full-stream models (Fig. 8). Both decoder families reused the frozen pose-trained YOLO/SPPF representation under matched training conditions, indicating that the central result did not depend on a single temporal head.

**Figure 8:**
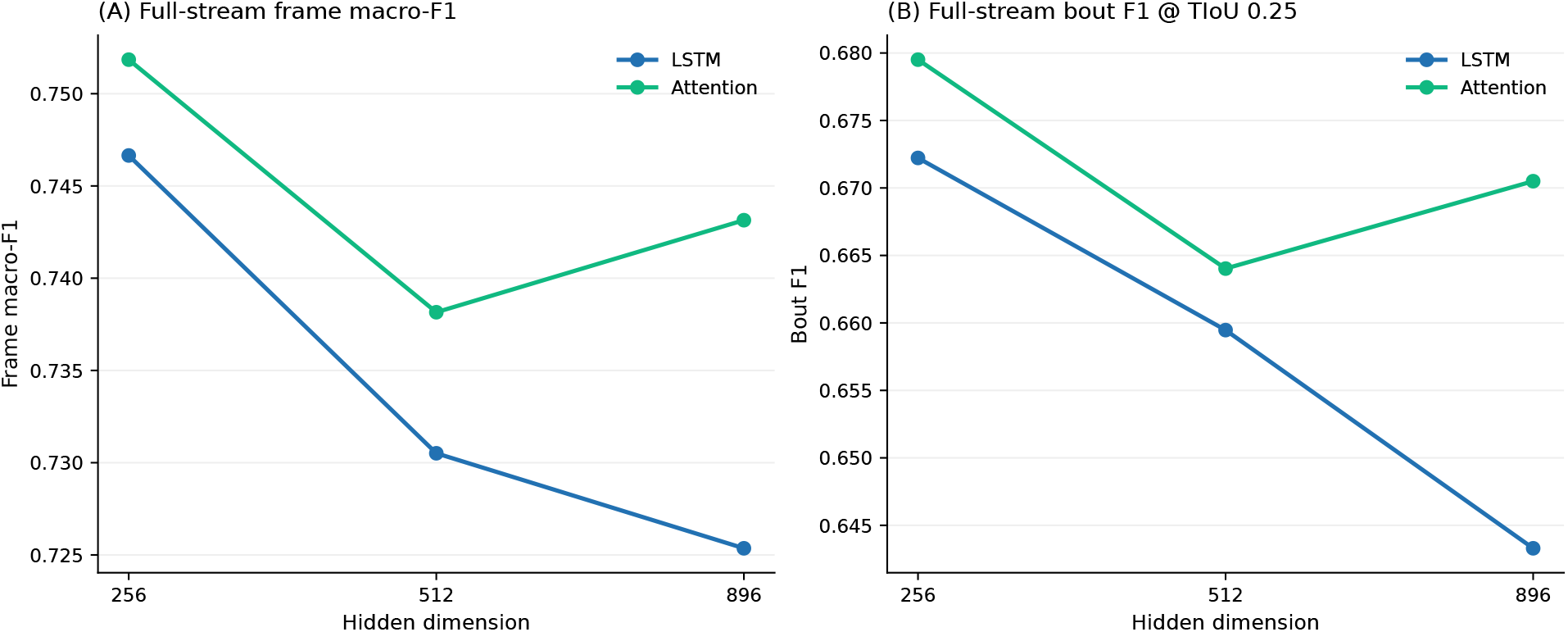
Full-stream performance across temporal head family and capacity. Full multimodal reuse was evaluated with LSTM and attention heads at 256, 512, and 896 hidden dimensions across three seeds. The 256-dimensional attention head gave the highest held-out frame macro-F1, while the matched LSTM head remained close, indicating that BehaviorScope-X is not dependent on one temporal decoder family.

### 3.3. Pose-derived geometry and animal-centered visual descriptors are complementary

The stream-ablation matrix asked which parts of the reused representation carry behavior signal. Full multimodal reuse combined self-pose geometry, pairwise relational geometry, group-scene visual descriptors, and per-animal visual descriptors. The remaining rows removed one or more streams while preserving the same held-out videos, model families, capacities, and seeds.

Full multimodal reuse was the strongest stream condition for both temporal heads (Fig. 9). Removing group-scene visual context caused only a modest reduction relative to the best full model: frame macro-F1 and bout macro-F1@0.25 were each lower by 0.013 for attention and by 0.013 for LSTM. In contrast, visual-only and per-animal-visual-only models lost substantially more performance. For attention, per-animal visual-only decoding was 0.072 lower in frame macro-F1 and 0.099 lower in bout macro-F1@0.25 than full reuse. For LSTM, the corresponding drops were 0.109 and 0.115. The ablations show that pose-derived geometry and pose-trained visual descriptors are complementary: animal-centered visual evidence is useful, but it is most informative when fused with explicit social geometry.

**Figure 9:**
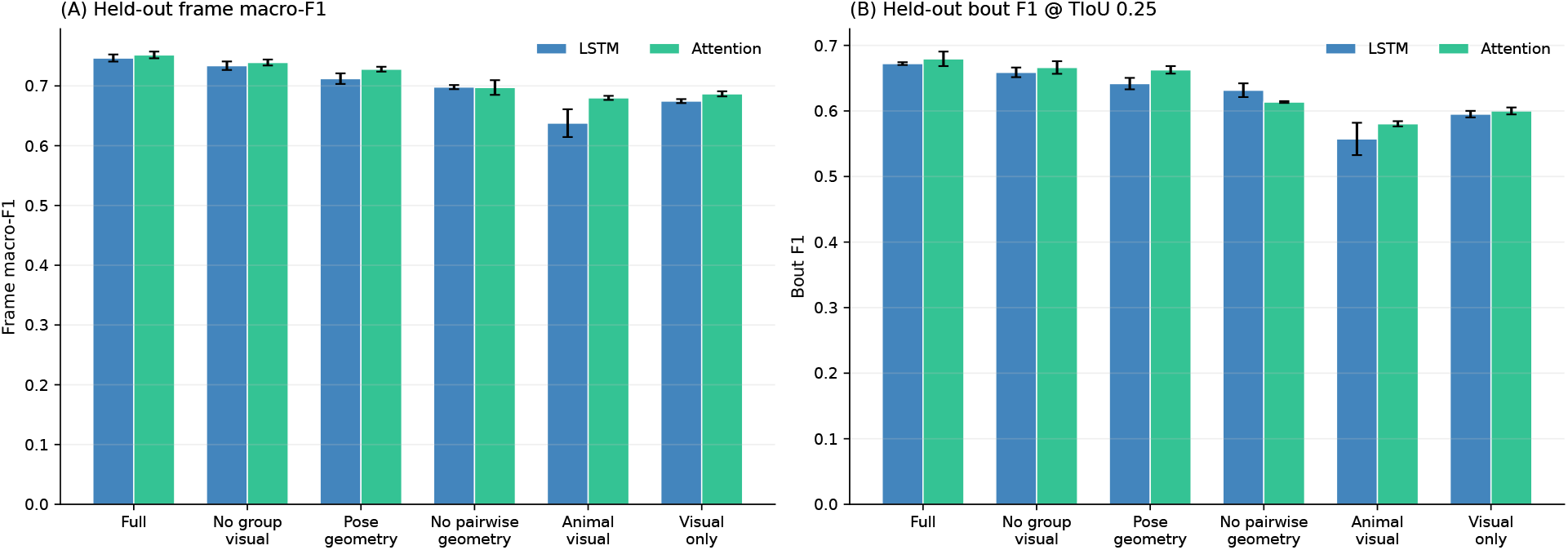
YOLO/SPPF stream-ablation matrix. Held-out frame macro-F1, bout macro-F1@0.25, and validation macro-F1 are summarized for LSTM and attention heads across stream conditions, capacities, and seeds. Values are seed means with seed standard deviations.

The paired-video analysis gave the same conclusion at the level of individual held-out recordings (Fig. 10). Full attention-256 exceeded the matched pose+relations-only attention-256 model by 0.031 frame macro-F1 (95% bootstrap CI, 0.018 to 0.047), 0.032 bout macro-F1@0.25 (95% CI, 0.016 to 0.049), and 0.042 bout macro-F1@0.50 (95% CI, 0.021 to 0.062). The contrast against visual-only attention was larger: 0.068 frame macro-F1, 0.083 bout macro-F1@0.25, and 0.123 bout macro-F1@0.50. The direct comparison between full attention-256 and full LSTM-256 was much smaller, with overlapping confidence intervals for frame macro-F1 and bout macro-F1@0.50. Thus, the dominant effect in this experiment was the reused input stream composition, not a large separation between the two temporal-head families.

**Figure 10:**
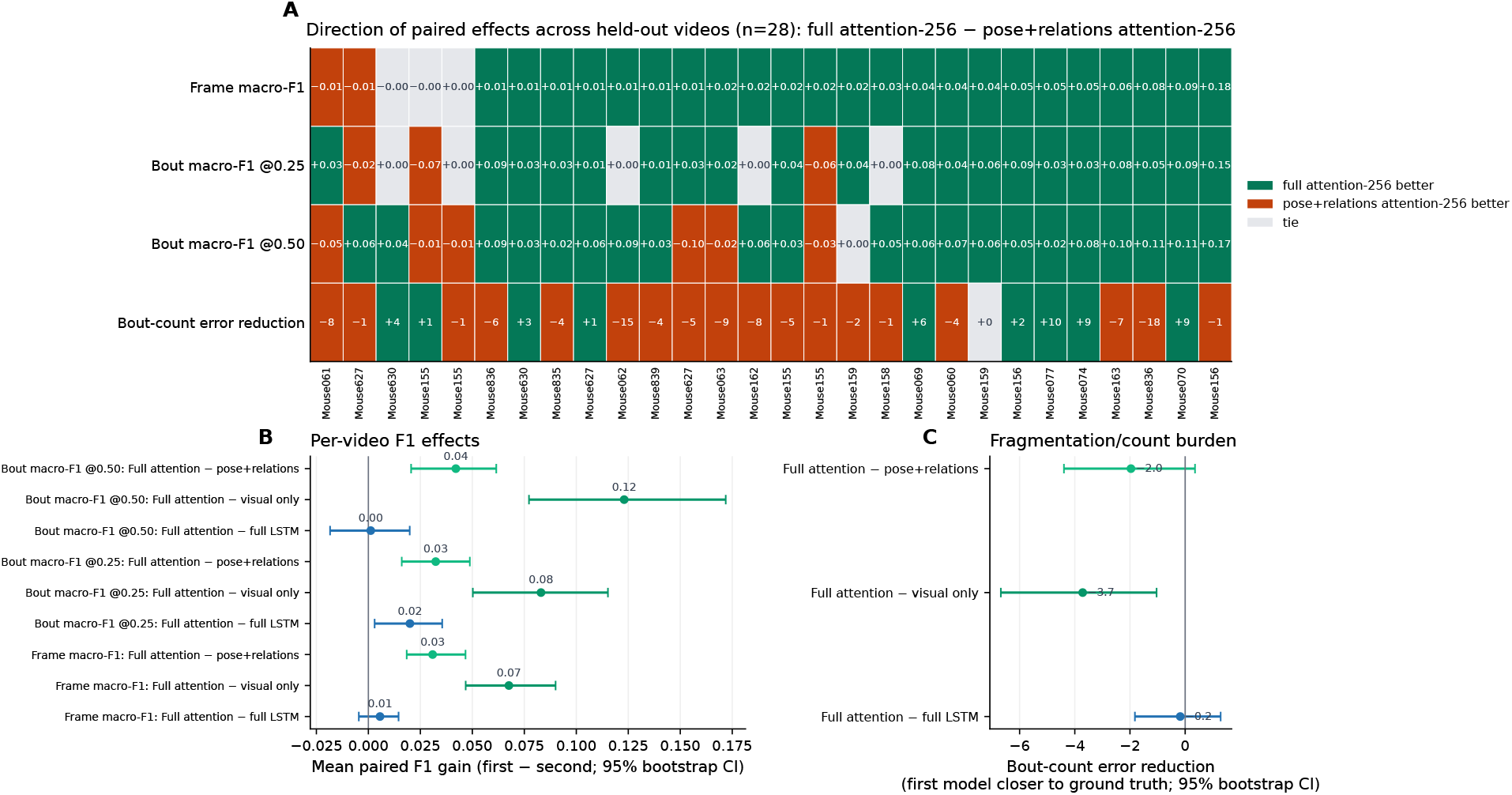
Per-video paired effects of multimodal reuse. Each held-out video is treated as the paired statistical unit. Full attention-256 is compared with the matched pose+relations-only attention-256 model, visual-only attention-256 model, and full LSTM-256 model. Positive values indicate that the first model in the comparison is closer to the human annotation.

### 3.4. Static/PCA baselines define a negative control for non-sequential reuse

The LSTM and attention stream-ablation results show that reused visual descriptors are useful when fused with pose-derived geometry by temporal sequence classifiers. The classical-baseline analysis served as a negative control for the flattened, non-sequential use of the same cached representation. To avoid a confounded comparison, the Random Forest and XGBoost models included window-level variants instead of only single-frame baselines. Pose-window models received the same 32-frame local context used by the LSTM and attention decoders, and visual descriptors were compressed with PCA fit only on the training split before window concatenation.

Under this controlled setup, temporal pose windows improved the classical models. Adding PCA-compressed visual descriptors to the pose-window inputs did not improve frame macro-F1 or bout F1@0.25 for either Random Forest or XGBoost; the only positive difference after adding visual PCA to pose-window inputs was a small Random Forest increase at TIoU 0.50 (Fig. 11). The LSTM and attention ablation matrices show that visual descriptors became useful when fused with pose geometry by sequence models. The classical baseline result therefore defines the cost of static, dimension-reduced reuse in this setting. After training-split PCA compression and window concatenation, the tabular models did not exploit the visual stream as effectively as the LSTM and attention heads. The comparison supports the broader interpretation that amortized pose vision depends on both cached content and a temporal classifier that can preserve and weight visual evidence across the behavior window.

**Figure 11:**
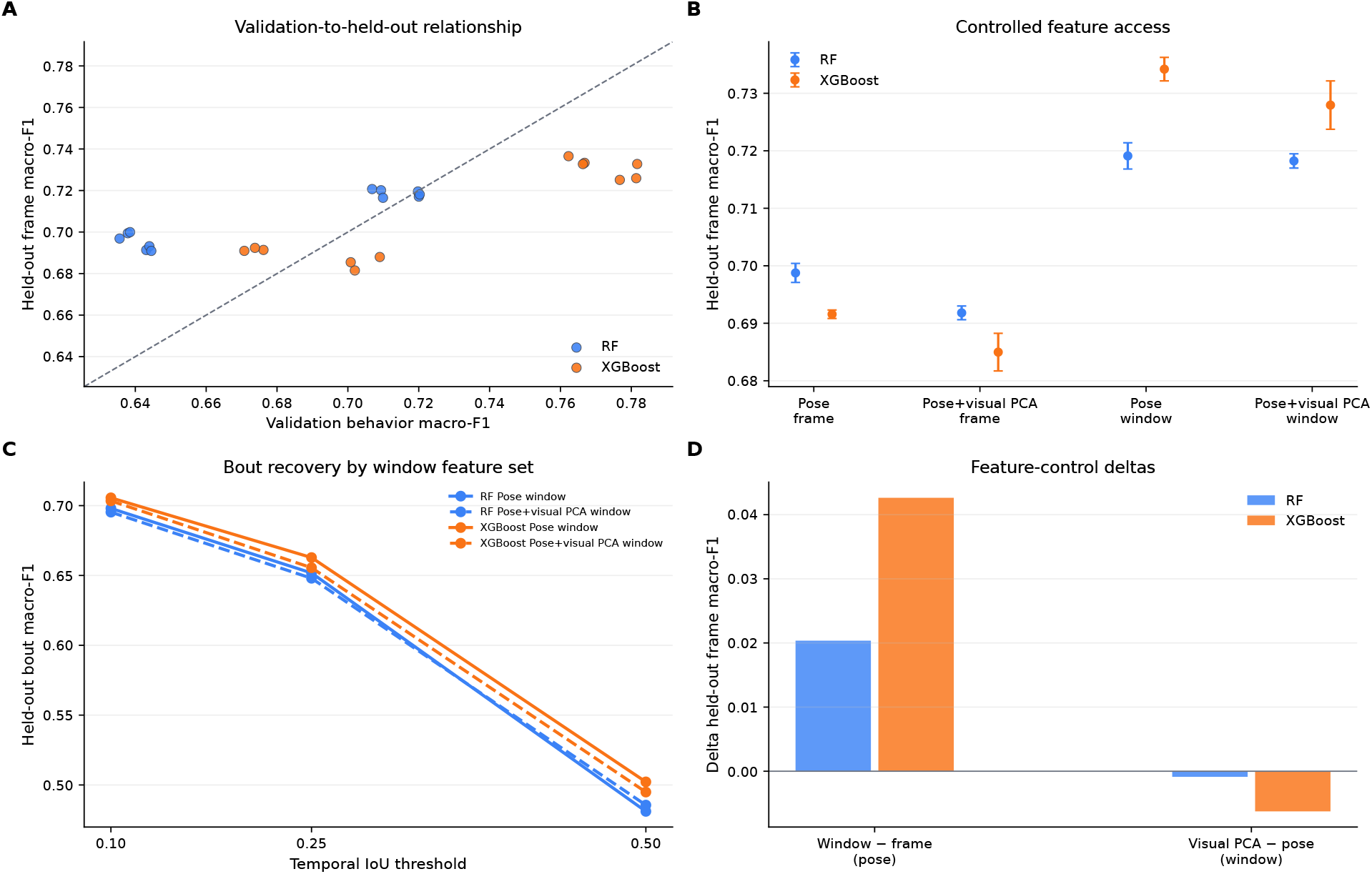
Controlled RF/XGBoost baselines with window-level inputs and PCA-compressed visual descriptors. Classical baselines were evaluated with pose-frame, pose-window, pose+visual PCA frame, and pose+visual PCA window inputs. The comparison tests whether tabular classifiers can exploit controlled access to the same frozen representation after static flattening and PCA compression, without learned temporal decoding. Frame macro-F1 is shown across feature sets and as window-versus-frame and visual-PCA-versus-pose-window deltas; bout macro-F1 is shown across TIoU thresholds for pose-window and pose+visual PCA-window inputs. The comparison is a negative control for non-sequential reuse and should not be interpreted as a direct test of the raw visual descriptors.

### 3.5. A native MobileNetV3 pose backbone supports pose-trained reuse across backbones

The YOLO/SPPF results establish the primary controlled comparison, and MobileNetV3 provides a structurally distinct cross-backbone control. We reused frozen descriptors from a MobileNetV3-large pose checkpoint without modifying the backbone to mimic YOLO/SPPF. The experiment then held the downstream ethogram workflow fixed: full-stream inputs, matched LSTM and attention heads, hidden dimension 256, and the same three seeds.

The MobileNetV3 pose checkpoint supplied a substantially larger native visual descriptor than the YOLO/SPPF tap: 960 dimensions per crop and 2,880 visual dimensions per frame, compared with 256 and 768 for YOLO/SPPF (Fig. 12A). In its pose-validation summary, the MobileNetV3 checkpoint reached pose mAP50 of 0.907, pose mAP50– 95 of 0.670, and PCK@0.05 of 0.950 across 3,000 validation images. These pose metrics supported its use as a second pose-trained representation source and defined a distinct provenance condition for the reuse experiment: a different model family, feature layer, and descriptor-size regime from the YOLO/SPPF route.

**Figure 12:**
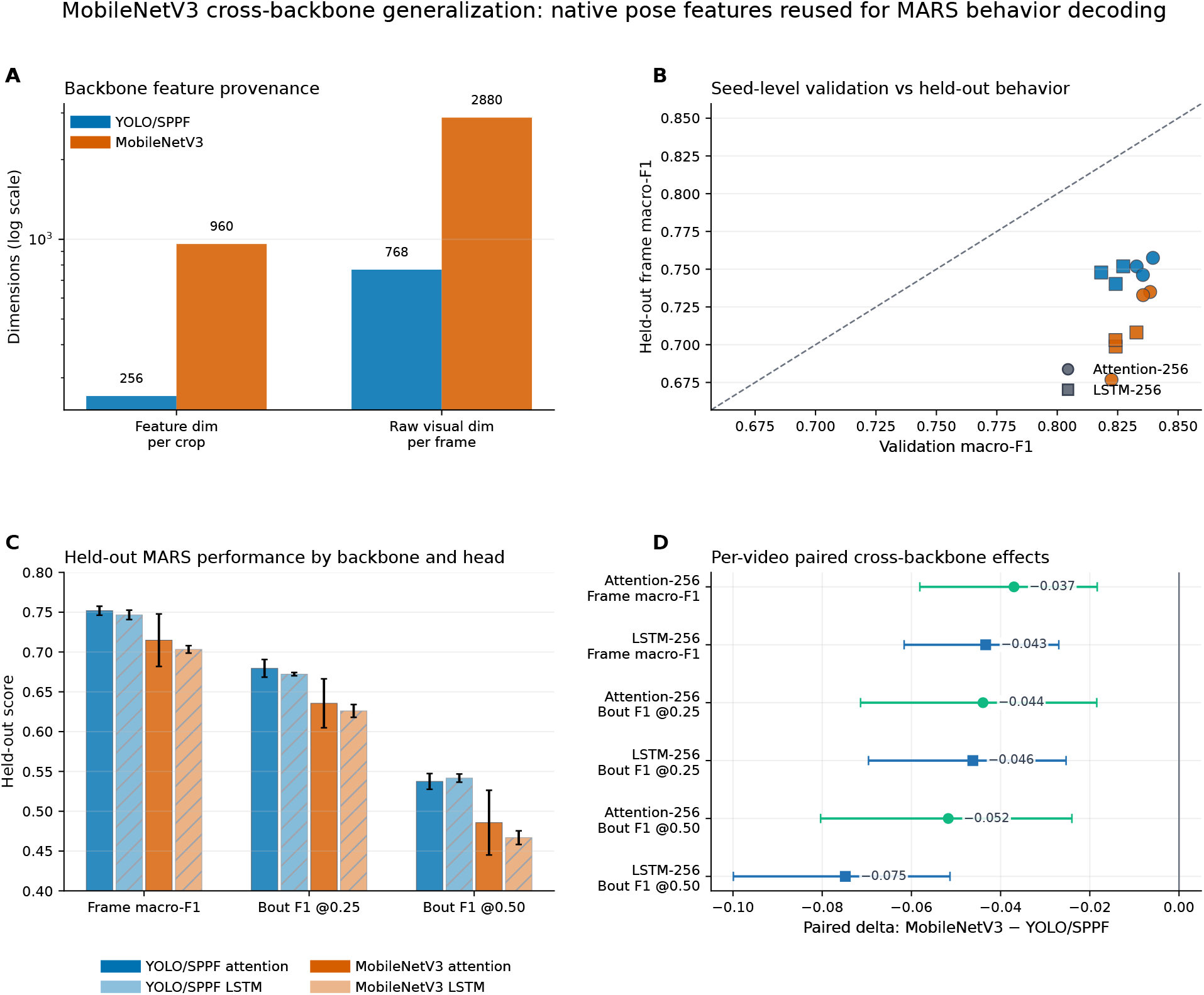
Native MobileNetV3 control for backbone-dependent reuse. (A) The MobileNetV3 feature source produced 960 dimensions per crop and 2,880 raw visual dimensions per frame, compared with 256 dimensions per crop and 768 raw visual dimensions per frame for the primary YOLO/SPPF feature tap. (B) Validation and held-out frame macro-F1 are shown for seed-level full-stream LSTM-256 and attention-256 models. (C) Held-out frame and bout metrics quantify backbone dependence under matched full-stream temporal heads. (D) Paired video-seed deltas report MobileNetV3 relative to the primary YOLO/SPPF route for frame macro-F1 and bout macro-F1 at TIoU 0.25 and 0.50.

When reused for behavior decoding, MobileNetV3 supported full-video MARS classification with both temporal heads, with lower scores than the primary YOLO/SPPF route (Fig. 12B–D). The MobileNetV3 attention-256 model reached frame macro-F1 0.715 ± 0.033, bout macro-F1@0.25 0.636 ± 0.031, and bout macro-F1@0.50 0.486 ± 0.041. The MobileNetV3 LSTM-256 model reached frame macro-F1 0.703 ±0.005, bout macro-F1@0.25 0.626 ±0.008, and bout macro-F1@0.50 0.467 ± 0.009. Paired video-seed comparisons showed that MobileNetV3 trailed YOLO/SPPF by 0.037 frame macro-F1 for attention-256 (signed 95% bootstrap CI for MobileNetV3 minus YOLO/SPPF, −0.058 to −0.018) and by 0.043 for LSTM-256 (signed 95% CI, −0.062 to −0.027). Bout-level differences showed the same direction: MobileNetV3 was 0.044 lower at TIoU 0.25 for attention-256 and 0.046 lower for LSTM-256, with larger separation at TIoU 0.50.

The ethogram-level analysis clarifies what was preserved and what degraded under the MobileNetV3 backbone. In a representative held-out recording, MobileNetV3 predictions followed the broad investigation and mount structure of the human ethogram, similar to the matched YOLO/SPPF models (Fig. 13A). Across held-out videos, however, the same bout-compression pattern seen in YOLO/SPPF persisted and was not improved by the larger MobileNetV3 descriptor. Investigation bout counts were approximately 0.62–0.66 predicted bouts per annotated bout across the four matched models, and predicted investigation bouts had median durations approximately 2.3–2.5 times the human median (Fig. 13B,C). MobileNetV3 also showed a slightly weaker bout-boundary profile, especially for attack and mount. At TIoU 0.25, median absolute boundary offsets were 16.7 frames for MobileNetV3 attention-256 attack bouts compared with 15.4 frames for YOLO/SPPF attention-256, and 10.5 frames for MobileNetV3 attention-256 mount bouts compared with 9.0 frames for YOLO/SPPF attention-256 (Fig. 13E).

**Figure 13:**
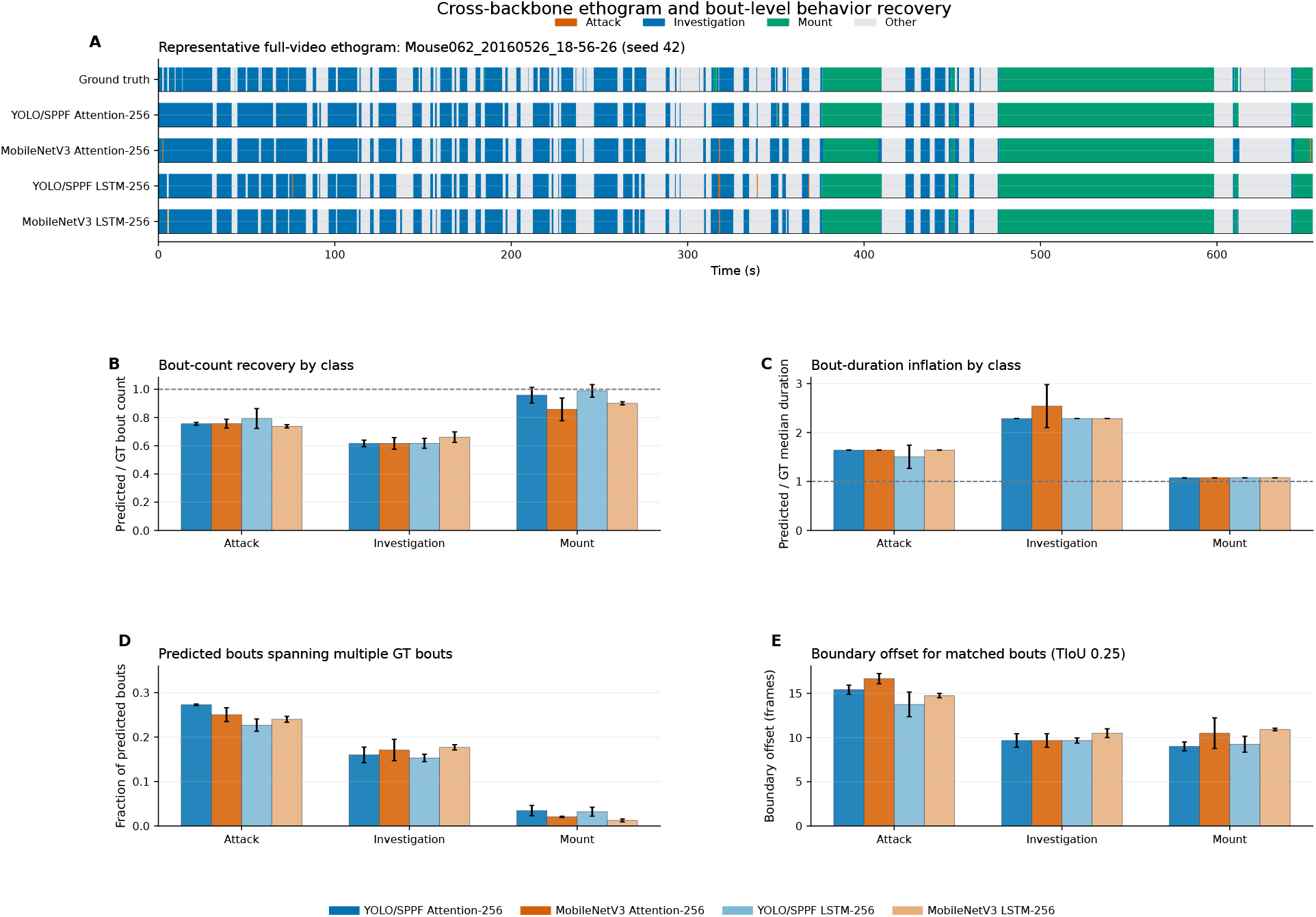
MobileNetV3 ethogram and bout-level behavior recovery. (A) Representative full-video ethograms compare ground truth with matched full-stream YOLO/SPPF and MobileNetV3 attention-256 and LSTM-256 predictions. (B) Bout-count recovery, (C) bout-duration inflation, (D) fraction of predicted bouts spanning multiple ground-truth bouts, and (E) matched-bout boundary offsets summarize bout-level structure across the held-out videos and three seeds.

The MobileNetV3 analysis shows that a structurally distinct non-YOLO pose backbone can be cached, reused, fused with pose-derived geometry, and decoded into full-video ethograms. This held even though the MobileNetV3 checkpoint was a substantially weaker pose estimator than the primary YOLO route (pose mAP50–95 0.670 versus 0.899): bout recovery degraded only modestly (bout macro-F1@0.25 0.636 versus 0.680), indicating that the cache-and-classify behavior readout is comparatively robust to pose-backbone quality. Its lower held-out scores and bout-boundary profile also establish that pose-trained reuse is backbone-dependent, motivating a descriptor-level analysis of what each route preserves in the frozen visual space once all three routes have been established.

### 3.6. DeepLabCut-HRNet reuses a top-down pose checkpoint beyond coordinate export

The YOLO and MobileNetV3 analyses tested representation reuse within native BehaviorScope-X pose routes. The DLC-HRNet experiment evaluated a complete DeepLabCut top-down route rather than an isolated feature layer. Detector outputs, keypoints, crops, relation features, and HRNet visual descriptors were all produced through the DLC path before downstream behavior classification. Within this route, a fine-tuned DeepLabCut pose checkpoint became both a coordinate estimator and a descriptor source. The same training investment provided body landmarks for coordinate analysis and frozen visual descriptors for behavior decoding, extending amortized pose vision into a widely used DeepLabCut workflow. DeepLabCut-style workflows are often used primarily to export coordinates; the DLC-HRNet route makes the same checkpoint a source of both coordinates and reusable visual descriptors. DeepLabCut SuperAnimal HRNet-W32 uses a top-down detector-plus-pose workflow and an HRNet backbone rather than a YOLO-style neck. Since HRNet-W32 has no SPPF module, we defined the reusable visual descriptor by functional position rather than layer name. The adapter tapped the raw four-branch HRNet backbone output before DLC branch collapse and before keypoint heads, globally pooled each branch, and concatenated the resulting 32 + 64 + 128 + 256 = 480-dimensional descriptor for each group and animal crop.

Using this validated tap, we built train/validation feature caches for 30,493 windows and trained the same Attention-256 behavior classifier used in the primary analyses, reaching validation macro-F1 0.788. We then evaluated all 28 manifest videos with accepted human annotations and 11,960 windows with no blocking failures. Raw-window predictions reached mean per-video accuracy 0.895, frame macro-F1 0.749, bout macro-F1@0.25 0.670, and bout macro-F1@0.50 0.555 (Fig. 14A). Applying the MARS smoothing recipe reduced DLC-HRNet performance to frame macro-F1 0.638, bout macro-F1@0.25 0.537, and bout macro-F1@0.50 0.443, primarily because smoothing expanded attack predictions and reduced other-class recall (Fig. 14B,C). Temporal smoothing is a postprocessing step that can interact differently with each pose route. In the DLC-HRNet route, raw outputs provide the primary held-out readout, while smoothed outputs serve as a postprocessing sensitivity analysis. We designed the DLC experiment as route-level validation of coordinate-and-descriptor reuse in an established DeepLabCut workflow. A DLC SuperAnimal HRNet-W32 detector-plus-pose workflow supported held-out behavior decoding after strict feature-tap validation. We required cache creation to stop unless the hook fired, four HRNet branches were returned, branch widths matched [32, 64, 128, 256], and the concatenated descriptor was 480-dimensional. Together, these results show that amortized pose vision can be instantiated in a native DLC top-down route. The route-level result validates coordinate-and-descriptor reuse in a native DLC workflow. It does not isolate the incremental contribution of HRNet descriptors relative to DLC-derived pose and relation streams alone.

**Figure 14:**
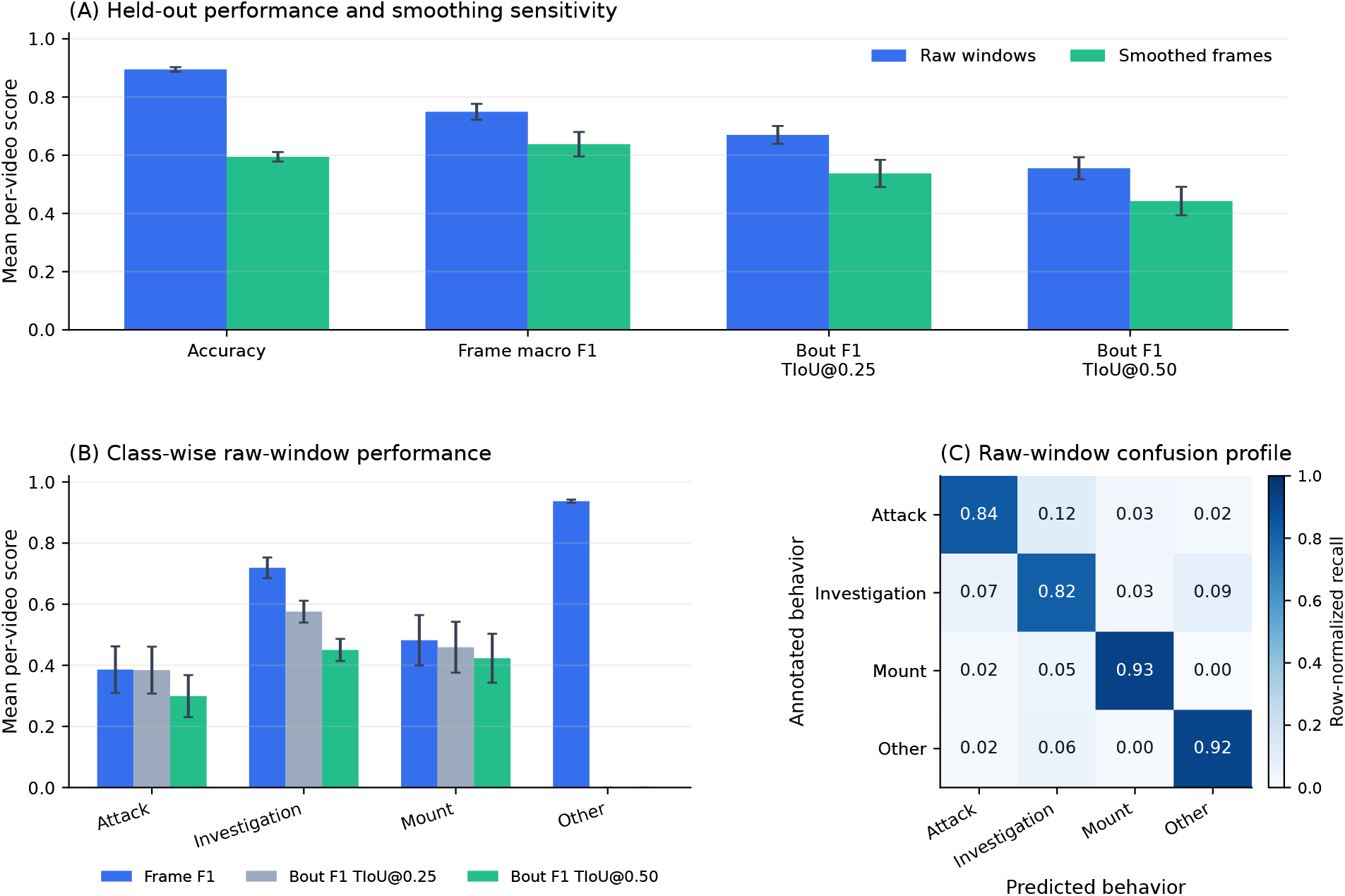
DLC-HRNet held-out behavior decoding in a native DeepLabCut top-down route. (A) Raw-window and smoothed-frame outputs are summarized across the 28 held-out MARS videos with accepted human annotations. Raw DLC-HRNet outputs provide the primary held-out readout because the MARS smoothing recipe reduced performance for this route. Error bars show SEM across held-out videos. (B) Per-class metrics identify which behavior classes drive the aggregate scores and show the class-specific effect of postprocessing. (C) Row-normalized held-out confusion for raw-window outputs shows how the route distributes errors across attack, investigation, mount, and other behavior.

### 3.7. Visual descriptor provenance links backbone-dependent reuse to behavior-accessible evidence

The YOLO/SPPF, MobileNetV3, and DLC-HRNet results establish three routes for amortized pose vision. We next asked how the frozen visual descriptors differed before temporal classification. This probe analysis treats the descriptors as measurement objects in their own right: if a pose checkpoint preserves behavior-relevant visual evidence, that evidence should be partly accessible from the cached visual space before pose-derived geometry and sequence modeling are added.

In 2,000 matched held-out windows balanced across behavior classes, YOLO/SPPF descriptors reached a behavior Fisher ratio of 54.0 and visual-only grouped linear-probe macro-F1 0.849 ± 0.015. MobileNetV3 descriptors were larger but less organized by behavior label, with Fisher ratio 26.1 and visual-only macro-F1 0.757 ± 0.028 (Fig. 15A). DLC-HRNet descriptors showed a high Fisher ratio of 63.3 together with lower and more variable visual-only macro-F1 0.704 ± 0.059.

**Figure 15:**
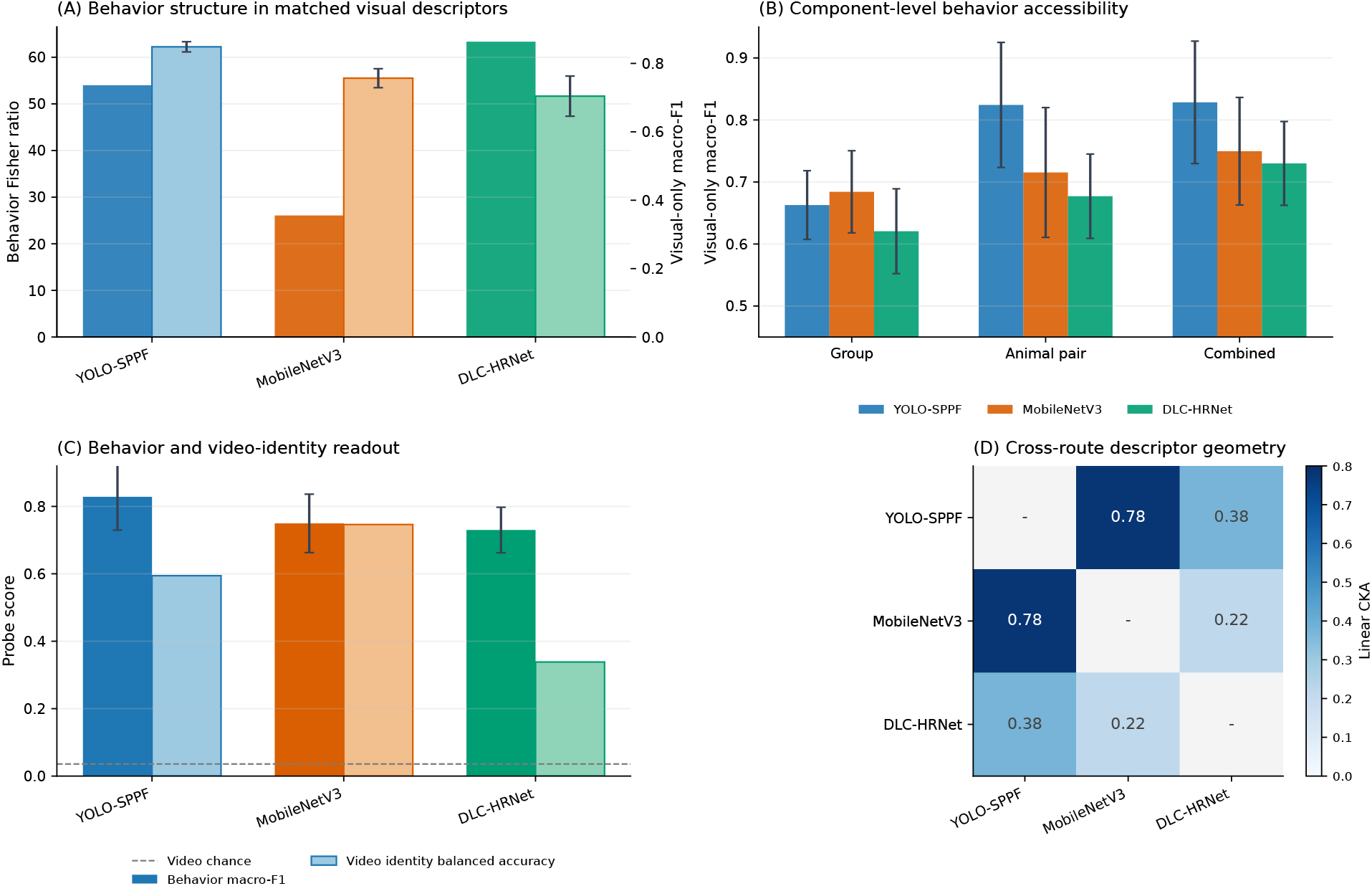
Visual descriptor provenance across pose routes. Matched held-out MARS windows were used to compare frozen visual descriptors from YOLO/SPPF, MobileNetV3, and DLC-HRNet before temporal behavior classification. (A) Behavior-label separability and visual-only grouped linear-probe macro-F1 measure how accessible behavior information is in each descriptor space. (B) Group-only, paired-animal, and combined probes show how scene context and animal-centered visual evidence contribute to behavior readout. Error bars in panels A–C show SD across five grouped behavior-probe folds where fold variability was estimated. (C) Video-identity probes quantify recording-specific structure in the same descriptors. MobileNetV3 descriptors preserved strong video-specific structure, with video-identity balanced accuracy close to behavior-probe accuracy for combined descriptors. The dashed line marks video-identity chance level. (D) Linear centered-kernel alignment shows route-specific descriptor geometry after removing trivial self-comparisons from the visual scale. Probe analyses used visual descriptors only.

Component probes identified where behavior-accessible visual evidence was concentrated. In 1,200 matched held-out windows balanced across behavior classes, YOLO/SPPF paired-animal descriptors reached visual-only macro-F1 0.824, close to the combined-descriptor value of 0.828 and higher than the group-scene value of 0.663 (Fig. 15B). This pattern is biologically consistent with a social-behavior readout in which animal-centered crops preserve contact, overlap, posture, and local appearance cues that are partly lost in sparse landmark coordinates. MobileNetV3 paired-animal descriptors reached macro-F1 0.715, while the combined descriptor reached 0.750. The same MobileNetV3 combined descriptors were also strongly predictive of video identity, with video-identity balanced accuracy 0.746 compared with behavior-probe accuracy 0.753 (Fig. 15C). For YOLO/SPPF, the corresponding behavior-probe accuracy was 0.828 and video-identity balanced accuracy was 0.594.

Linear centered-kernel alignment showed route-specific descriptor geometry. YOLO/SPPF–MobileNetV3 similarity was 0.78, YOLO/SPPF–DLC-HRNet similarity was 0.38, and MobileNetV3–DLC-HRNet similarity was 0.22 (Fig. 15D). The descriptor probes therefore provide a mechanistic bridge between the route-level results above and the feature-provenance argument developed in the Discussion. Pose-trained visual descriptors carried behavior signal across routes, but the form of that signal depended on the feature tap, crop definition, and pose framework that produced the cache. Together, these route-specific patterns indicate that behavior-label separability, video-identity structure, descriptor geometry, and classifier accessibility are distinct properties of pose-trained visual evidence.

### 3.8. Full-video ethograms recover sustained episodes but compress dense short-bout structure

Frame-level and bout-level scores summarize performance, but neuroethology experiments ultimately depend on ethograms: the timing, duration, and sequence of behavior bouts across complete recordings. We examined the held-out predictions as continuous ethograms and compared them against human annotation at multiple temporal scales.

The representative ethograms show that full multimodal reuse reconstructs sustained investigation and mount episodes and can correct behavior identity errors made by pose+relations-only decoding (Fig. 16). However, when ethograms were analyzed as ordered behavior bouts, the models also showed a consistent bout-compression pattern (Figs. 17 and 18). For the representative full-video example, the human annotation contained 79 behavior bouts, whereas full attention-256 produced 54 and full LSTM-256 produced 63. Across all held-out videos, investigation was the clearest example of this under-segmentation: full attention-256 produced approximately 0.62 investigation bouts per annotated investigation bout, and the median predicted investigation duration was approximately 2.29 times the human median duration. Bout compression occurred despite the MARS annotation itself containing a high frequency of naturally short biological episodes: median ground-truth bout durations were 39 frames for attack, 21 frames for investigation, and 74.5 frames for mount, corresponding to 1.30, 0.70, and 2.48 seconds at 30 frames/s. Investigation was especially sensitive to temporal compression, with 37.7% of annotated investigation bouts lasting 15 frames or less. Overall, 64.9% of investigation bouts fit within a single 32-frame model window. This set included the 37.7% of investigation bouts that lasted 15 frames or fewer. The same compression pattern was present in raw window predictions and in outputs after smoothing and minimum-duration filtering, indicating that it was not introduced primarily by postprocessing.

**Figure 16:**
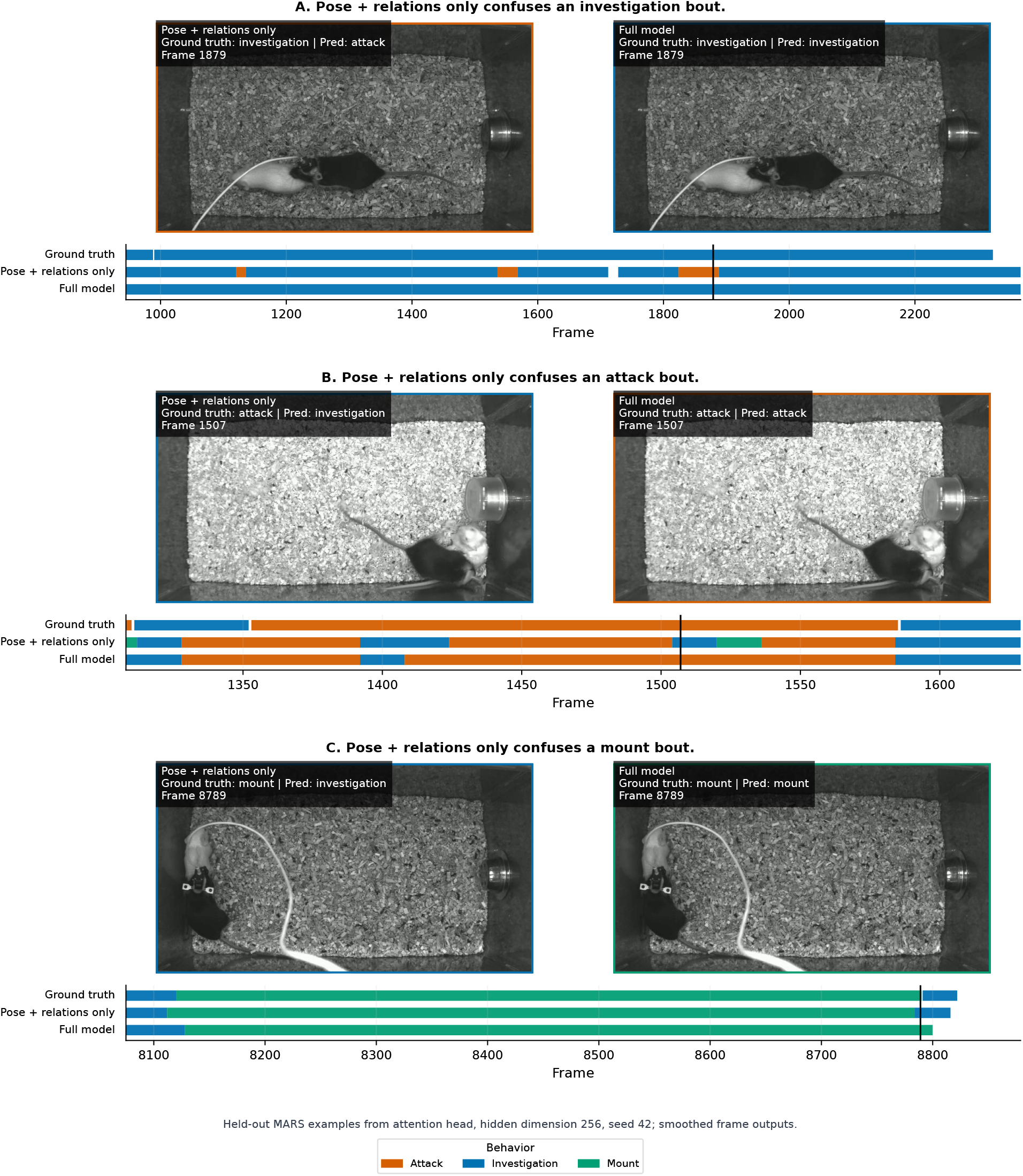
Representative bout-level examples from held-out MARS recordings. For attack, investigation, and mount examples, the pose+relations-only model is compared with the full multimodal model. The examples illustrate cases in which visual descriptors help resolve behavior identity and bout placement.

**Figure 17:**
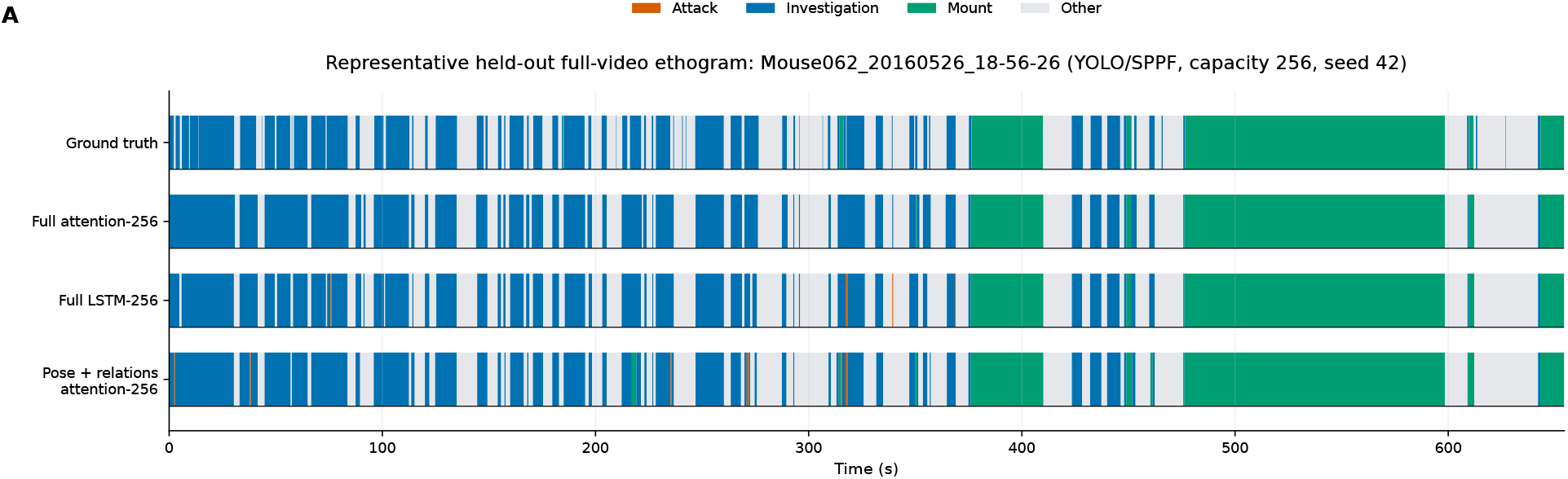
Full-video ethogram reconstruction and bout-to-bout transition structure. A representative held-out video is shown with ground truth, full attention-256, full LSTM-256, and the matched pose+relations-only attention-256 ablation. Transition matrices summarize behavior-bout to behavior-bout transitions across all 28 held-out videos with accepted human annotations; model panels show model-minus-ground-truth transition probabilities.

**Figure 18:**
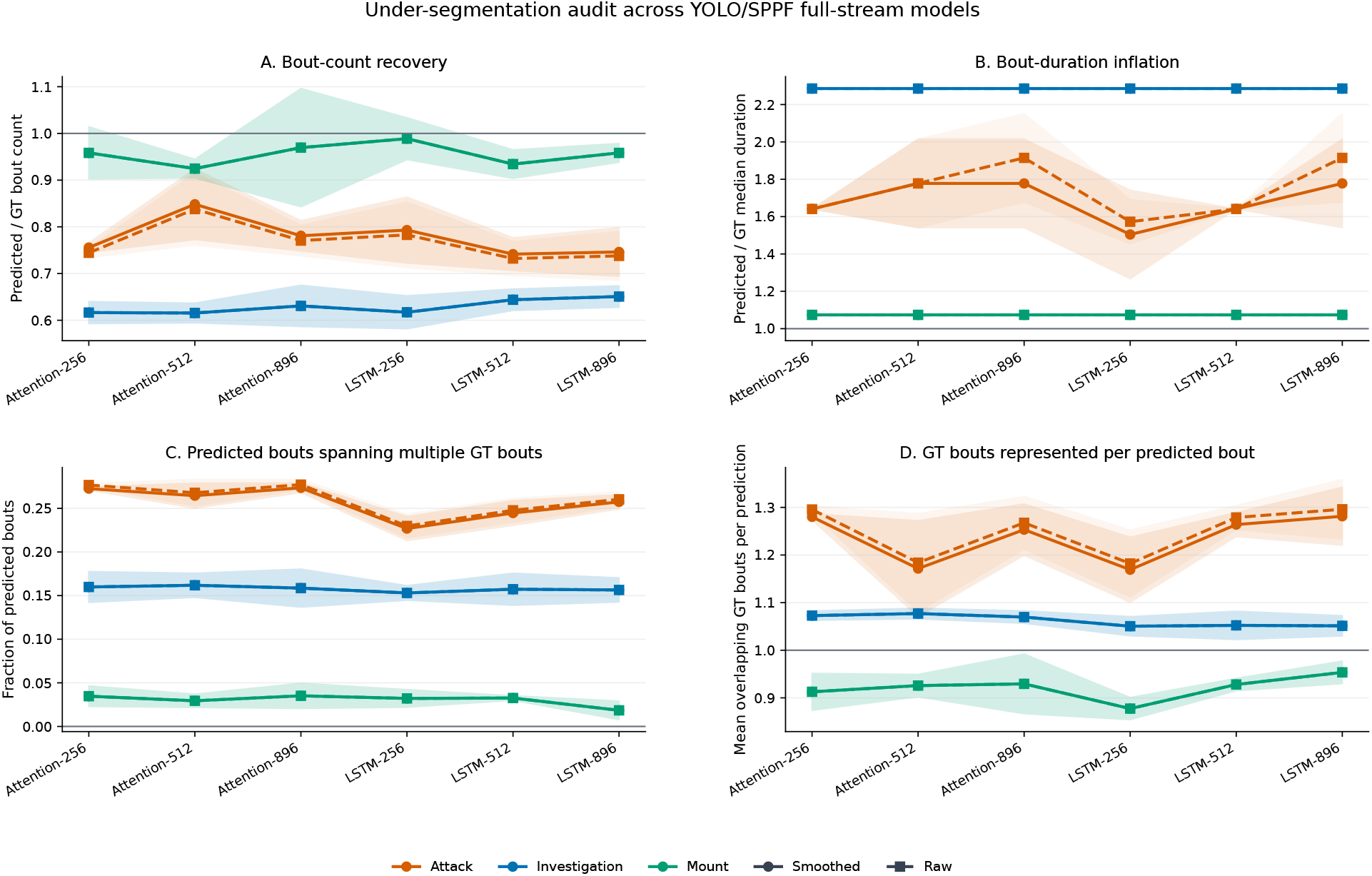
Bout under-segmentation audit across raw and postprocessed outputs. The audit compares predicted and ground-truth bout counts, same-class consecutive bout rates, same-class gap distributions, and matched-bout fragmentation. Raw window predictions and outputs after smoothing and minimum-duration filtering are shown together to test whether postprocessing is responsible for bout compression.

The bout-compression results are consistent with a boundary-specific limit of the current MARS implementation. Cached visual descriptors are extracted from a frozen spatial pose backbone before temporal decoding, so adjacent frames are not jointly encoded at the visual-feature stage. Downstream LSTM and attention heads can integrate sequences of cached descriptors, but they may not recover boundary evidence that would require earlier spatiotemporal fusion, behavior-specific visual updates, or shorter-window temporal recipes. This late-fusion structure may be well suited to sustained behavioral states, where posture, contact, relative geometry, and appearance remain stable across multiple windows, and less well matched to rapid event boundaries, where the discriminative signal may be a short transition rather than a persistent state. The observation that bout compression appears in raw-window outputs, before smoothing is applied, indicates that the pattern is not solely a postprocessing artifact. It marks a temporal operating limit of reusing frozen spatial pose descriptors with late temporal fusion for frame-scale event segmentation. The MARS held-out evaluation used one accepted human annotation file per evaluated video (Segalin et al., 2021a); as a result, this analysis supports a segmentation-compression interpretation but does not estimate whether MARS bout-boundary differences fall within inter-annotator temporal variability. Fly-v-Fly provides the explicit inter-annotator uncertainty analysis because secondary annotation files were available there.

### 3.9. Behavioral sequence structure is preserved beyond simple base-rate matching

Since bout under-segmentation can distort strict transition matrices, we tested whether predicted ethograms still preserve behavior sequence structure beyond class base rates. We analyzed behavior n-grams from the ordered bout sequence and compared model n-gram recovery against two nulls: a composition-shuffled ground-truth sequence and a Markov sequence generated from first-order transition statistics. The sequence analysis evaluates whether the model recovers ethologically meaningful local behavior motifs, rather than isolated frames or individual bouts alone.

Full attention-256 recovered behavior-sequence structure across increasing n-gram order. Across seeds, its n-gram F1 was 0.792 ± 0.014 for unigrams, 0.743 ± 0.013 for bigrams, and 0.682 ± 0.014 for trigrams. The n-gram scores are not interpreted in isolation, because MARS ethograms contain strong base-rate and transition regularities. Instead, the null comparisons in Fig. 19 provide the relevant control: the model preserved local behavior motifs beyond what would be expected from composition alone, while the under-segmentation audit explains residual differences in rapid self-transitions and micro-alternations. The ethogram result defines an operating characteristic of the framework. The YOLO/SPPF reuse model preserves much of the biological sequence grammar of the recording, while compressing some dense short-bout structure.

**Figure 19:**
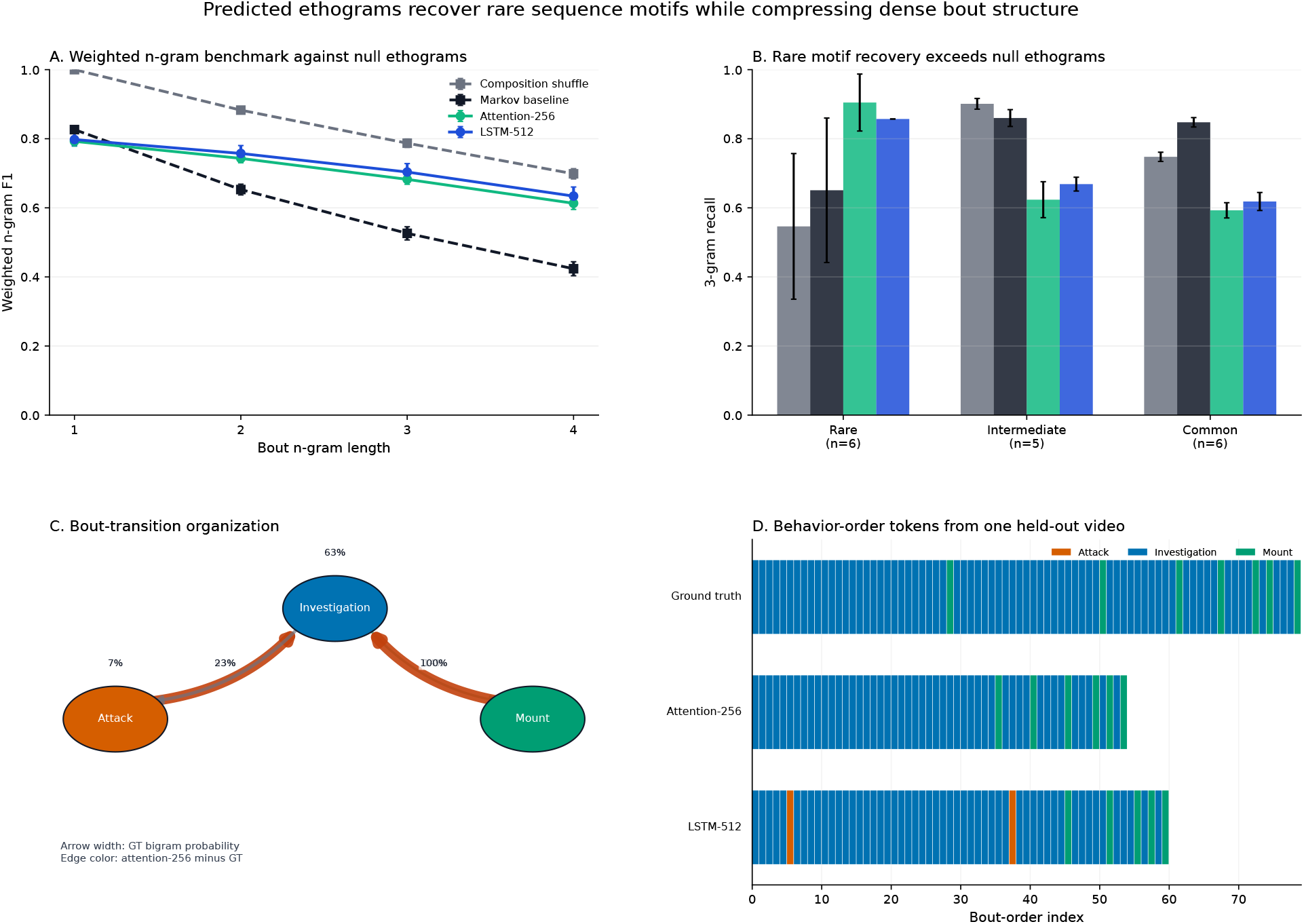
Ethogram sequence grammar relative to null baselines. N-gram recovery is compared against composition-shuffled and Markov null sequences, rare behavior motifs are summarized separately, and the node-edge graph visualizes ground-truth bout grammar with model-minus-ground-truth differences.

### 3.10. Bout recall defines the temporal operating envelope of the YOLO/SPPF reuse model

Finally, we quantified the temporal scale at which strict bout recovery breaks down. Ground-truth bouts were binned by duration, and recall was computed at TIoU 0.25 and TIoU 0.50 for full attention-256 and full LSTM-256. Both raw and smoothed outputs were retained so that duration-dependent recall could be separated from postprocessing effects.

The operating-envelope analysis confirms that strict bout recovery is duration dependent (Fig. 20). Short attack and investigation bouts are the most sensitive to boundary placement and merging, whereas longer sustained bouts are recovered more consistently. Neither attention-256 nor LSTM-256 reached 80% recall for every behavior class at TIoU 0.25 or TIoU 0.50, emphasizing that the framework should not be described as perfectly reconstructing all manual-annotation bout boundaries. Instead, the appropriate interpretation is more specific and more informative: amortized pose vision with a pose-trained YOLO/SPPF representation supports accurate frame-level ethograms and substantial recovery of behavior episodes, while strict sub-bout segmentation remains the main temporal limitation of the current frozen-descriptor, late-fusion architecture on MARS. The remaining sub-bout segmentation error identifies a specific target for future boundary-aware or earlier spatiotemporal variants, without implying that it is intrinsic to representation reuse.

**Figure 20:**
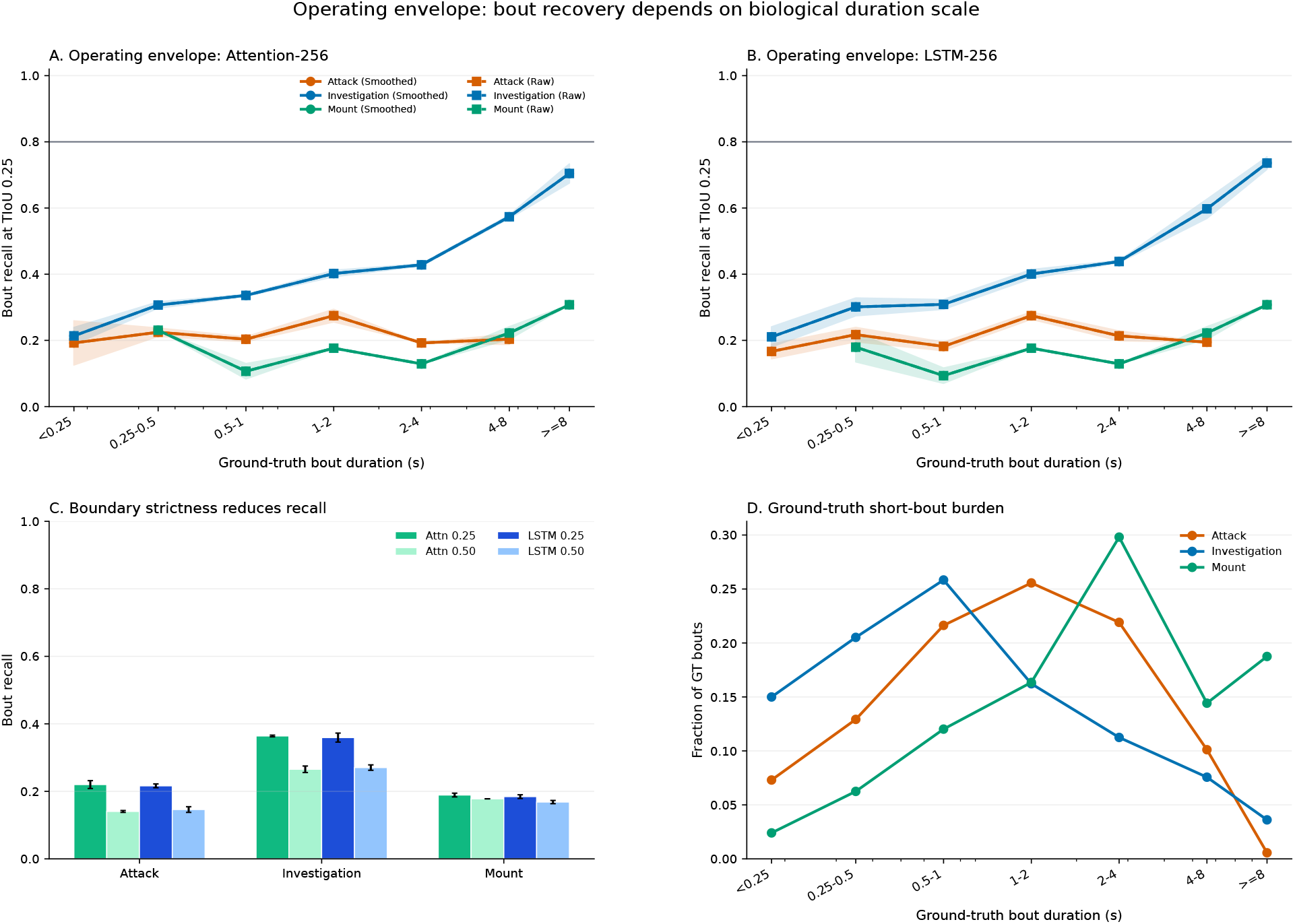
Duration-dependent bout recall defines the temporal operating envelope. Bout recall is shown as a function of ground-truth bout duration for full attention-256 and full LSTM-256, using raw window outputs and outputs after smoothing and minimum-duration filtering. The curves quantify the bout durations at which strict temporal overlap becomes reliable for each behavior class.

### 3.11. Annotation uncertainty constrains sub-second Fly-v-Fly evaluation

Fly-v-Fly aggression was used as a stress test of amortized pose vision at a shorter behavioral time scale than MARS resident-intruder behavior. Unlike the MARS analyses, this experiment does not isolate temporal scale alone: it also changes species, morphology, behavior definition, and annotation reliability. We first quantify annotation uncertainty to define the evaluation context for downstream model comparisons. Movies 6–10 were assigned to the held-out test set, each recording contained 54,000 frames, and primary annotations were converted to the same priority-expanded frame labels used by training and evaluation (per-frame labels after fixed-priority resolution of overlapping Fly behaviors). Secondary annotation was available only for movie 6, so we report the interannotator comparison in text rather than as a sparse table.

The held-out Fly-v-Fly ethogram made the shorter time scale explicit. Across movies 6–10, the priority-expanded ground truth was dominated by brief events, placing this evaluation in a temporal regime distinct from the MARS analyses. Lunge was the most common behavior, with 1,590 held-out bouts and a median duration of four frames. Charge was rarer, with 57 held-out bouts and a median duration of one frame. Tussle, hold, and wing threat had held-out median durations of 8, 12, and 13 frames, respectively (Table 4). At 30 frames/s, these medians place the shortest Fly-v-Fly behaviors at approximately 0.03–0.13 s. The focused w16/s8 and w8/s4 window recipes, as well as the shorter Fly-specific smoothing and minimum-duration settings, were chosen to match the temporal scale of the annotations.

The same short-bout structure also made annotation uncertainty part of the result. Small boundary differences can change both frame labels and bout matches, so model scores must be read against the available human-reference evidence. For reference, Eyjolfsdottir et al. reported that the original Fly-v-Fly work included a second annotation layer for human-performance reference, in which trained novice annotators re-annotated part of the test set, including one tenth of the Aggression movies (Eyjolfsdottir et al., 2014; Eyjolfsdottir, 2014). For Aggression, reported human recall was generally above 80%, whereas precision was in the 60–80% range (Eyjolfsdottir, 2014). The available Aggression subset showed the same limited second-layer coverage: among the held-out Aggression movies, only movie 6 contained a secondary annotation file. Within that video, overall frame agreement was high because most frames were other, but same-label agreement on primary non-other frames was 55.0%. The disagreement was also class-specific. Wing threat showed 95.5% same-label agreement, whereas lunge showed 18.5% agreement and 69.1% of primary-labeled lunge frames were labeled other by the secondary annotator. Hold showed 16.7% agreement as hold and was primarily confused with tussle. The human-reference evidence motivated the Fly-v-Fly reporting strategy below: precision and recall are reported alongside frame-level and bout-level macro-F1, and boundary sensitivity is evaluated against the 21-frame padded-primary margin estimated from the movie 6 secondary annotation.

### 3.12. Fly-v-Fly pose provenance supports but bounds short-bout decoding

The Fly adaptation also required pose-provenance checks, because it used a species-specific pose checkpoint rather than the MARS YOLO-pose model. The Fly YOLO-pose checkpoint reached pose mAP50 of 0.841 and pose mAP50– 95 of 0.765 on its validation ground truth (Fig. 5). We also evaluated the checkpoint against held-out Fly-v-Fly track geometry as used by the behavior pipeline. Across 132,497 matched animal-frames, the mean OKS-style score was 0.941 with a mean body length of 19.0 pixels (Fig. 21). Centroid localization was nearly exact (PCK@0.10, 1.000), body-axis endpoints were stable (PCK@0.10, 0.961 for each endpoint), and wing keypoints were less precise but still high overall (PCK@0.10, 0.966–0.971). The weakest pose-localization conditions were rare interactive behaviors, particularly charge, hold, and tussle. Thus, the Fly checkpoint provided a usable upstream representation source, while wing localization and short-bout class rarity remained important provenance for behavior decoding.

**Figure 21:**
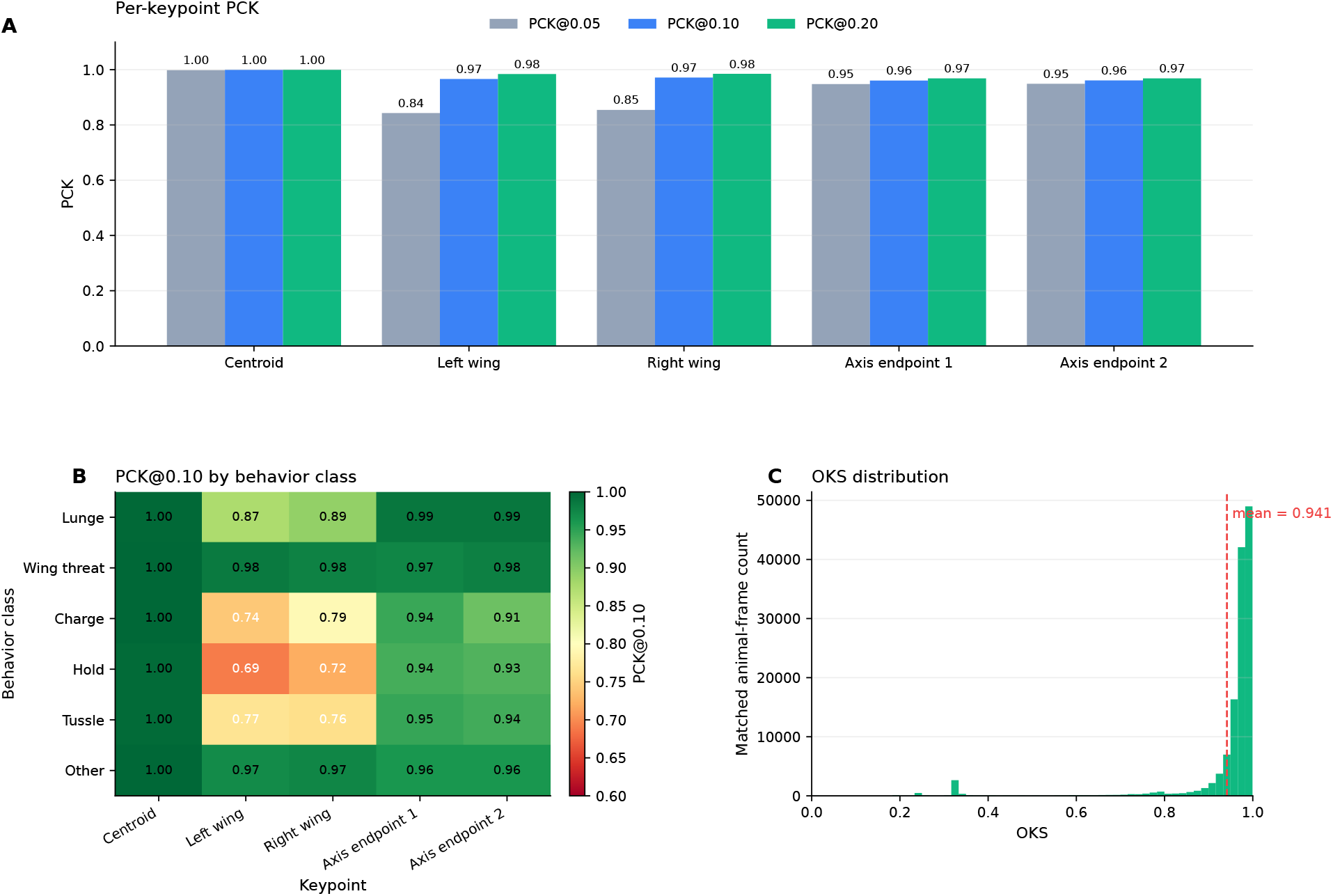
Fly-v-Fly pose-checkpoint error profile on held-out track geometry. Cached YOLO-pose keypoints from the held-out Fly-v-Fly evaluation cache were compared with keypoint geometry reconstructed from the public Fly-v-Fly tracking files. (A) Per-keypoint PCK at three body-length-normalized thresholds. (B) PCK@0.10 by behavior class. (C) OKS-style matched-frame distribution.

### 3.13. Fly-v-Fly short-bout decoding tests retuning under sub-second behavior

Within this annotation- and pose-provenance context, we compared temporal decoding windows as a bounded test of short-bout recovery. The Fly-v-Fly analysis measured how far cached pose-trained descriptors and compact temporal heads can be pushed when event boundaries approach the reliability limit of manual labels. The focused classifier comparison used the short-window adaptation most closely connected to the MARS results: an attention temporal sequence classifier with 256 hidden units and a single prespecified seed, using either 16-frame or 8-frame windows. The shorter window improved strict primary-label behavior decoding. Mean frame macro-F1 increased from 0.376 to 0.441, with the gain driven primarily by higher recall (0.555 to 0.627) and a smaller increase in precision (0.386 to 0.412; Fig. 22C). Bout recovery moved in the same direction. At TIoU 0.25, macro-F1 increased from 0.237 to 0.374, and at TIoU 0.50 from 0.163 to 0.214 (Fig. 22D).

**Figure 22:**
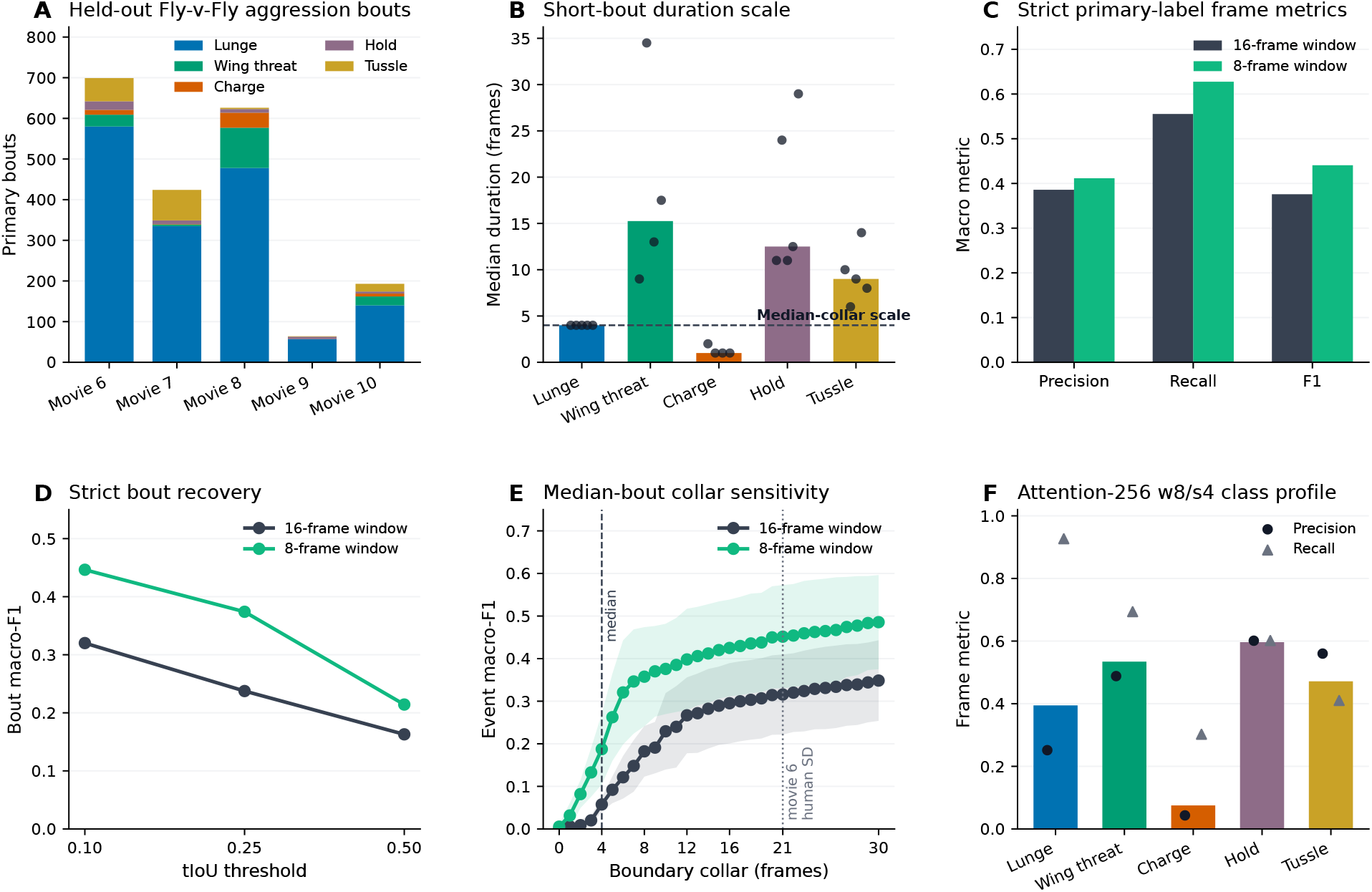
Focused Fly-v-Fly short-bout stress test. (A) Held-out Fly-v-Fly primary bout counts by movie and behavior class. (B) Behavior-specific median bout durations at 30 frames/s. (C) Strict primary-label frame precision, recall, and macro-F1 for the focused attention-256 adaptation runs. (D) Strict bout macro-F1 across TIoU thresholds. (E) Boundary-collar event macro-F1 sweep, with the median bout scale and movie 6 human-boundary reference marked. (F) Class-specific frame F1 for the 8-frame attention model, with precision and recall shown separately.

The class profile explains how this aggregate gain was achieved. For the 8-frame attention model, hold and wing threat had the strongest frame-level F1 values (0.596 and 0.534), tussle was intermediate (0.471), and charge remained weak (0.075) because it was both rare and often only one frame long. Lunge showed the opposite profile: recall was high (0.927), but precision was low (0.252), consistent with broadening of the most frequent short-bout class. Thus, the same properties that motivated the focused Fly-v-Fly adaptation also defined its main failure modes: rare one-frame behaviors, boundary-sensitive lunge bouts, and classes with larger manual-annotation ambiguity.

Boundary-collar analysis made this temporal limitation explicit. The 8-frame model reached half of its sweep maximum by a 5-frame collar and 90% of its maximum by a 19-frame collar, close to the 21-frame manual-annotation boundary-uncertainty reference estimated from movie 6 (Fig. 22E). At a four-frame collar, matching the median held-out bout duration, the 8-frame window reached event macro-F1 of 0.187 compared with 0.058 for the 16-frame window. As the collar widened, both models improved, but the shorter window retained the advantage: the 8-frame model reached a maximum mean event macro-F1 of 0.486 across the collar sweep, compared with 0.348 for the 16-frame model. Thus, boundary placement, in addition to class identity, is a major determinant of Fly-v-Fly bout scores.

Taken together, the Fly-v-Fly results extend the amortized pose vision analysis to a different species and a substantially shorter behavioral time scale. The cache-and-classify design transferred without changing the upstream reuse principle, and the short-window adaptation improved both frame-level and bout-level recovery. The absolute scores remain bounded by the available annotation regime, one-frame and four-frame events, strong class imbalance, and wing-dependent pose uncertainty. Fly-v-Fly supports the central reuse principle while defining the conditions needed for sub-second ethogram recovery. Reusable pose-trained features can support cross-assay short-bout decoding when the temporal window and class-balance settings are adapted to the dataset, while dense annotation and boundary-aware training are likely needed for MARS-level ethogram recovery in this sub-second regime.

### 3.14. End-to-end inference supports laboratory-scale deployment

Having established that cached pose-trained representations contain reusable behavior-discriminative evidence, we next evaluated the same cache-and-classify decomposition as a full-video deployment path. The laboratory-scale benchmark used the selected full-stream 256-capacity LSTM and attention temporal sequence classifiers on three held-out MARS videos. Unlike the training and held-out accuracy comparisons, which were tied to the controlled analysis environment, this deployment benchmark treated hardware as the measured variable. Each run included video loading, YOLO-pose tracking, crop construction, frozen YOLO/SPPF feature extraction, temporal decoding, postprocessing, CSV output, and measurement of runtime resource use.

The standard OpenCV/Ultralytics inference path processed 125,152 frames across six model-video runs at an aggregate 25.4 frames/s on an RTX2080 workstation and 32.6 frames/s on an RTX4080 workstation. On Apple Silicon MPS, the same benchmark averaged 60.7 frames/s. Thus, the tested implementation was near-real-time on an older discrete GPU and real-time or faster on the RTX4080 and Apple Silicon systems. The temporal decoder was not the dominant runtime cost. Classifier time remained a minor component of interval wall time, averaging less than 1% in the stage-timed RTX2080 OpenCV and RTX4080 profiling runs.

A secondary profiling run tested whether video preparation accounted for the remaining discrete-GPU overhead. FFmpeg predecode/cache modestly increased RTX4080 median interval throughput to approximately 36 frames/s, but did not approach the MPS throughput. Stage-timed profiling instead localized most of the remaining cost upstream of classification. In the completed profiling rows, the combined pose-and-crop path accounted for approximately 62– 74% of interval wall time, frozen YOLO/SPPF feature encoding for approximately 19–30%, window assembly for approximately 6%, and temporal classification for less than 1%. Thus, the tested deployment path is already near-real-time for full-video analysis, while further speed gains are most likely to come from optimizing frame handling, pose tracking, crop construction, and frozen feature extraction rather than from changing the temporal sequence classifier, as discussed in the future-directions section.

## 4. Discussion

The main implication of this study is a shift in how pose-estimation outputs should be interpreted in computational ethology. Coordinate export remains a powerful measurement strategy, but pose training can also produce a second scientific product in the form of behavior-discriminative visual evidence adapted to the assay. BehaviorScope-X shows that this evidence can be recovered for continuous ethology and evaluated as part of the behavioral measurement itself.

The evidence for that interpretation comes from both ablation and route-level tests, which anchor the conceptual point in held-out behavior measurements. In MARS, adding visual descriptors to pose-plus-relations inputs improved held-out frame macro-F1 by 0.031 and bout macro-F1@0.25 by 0.032 (Fig. 10). Descriptor probes provided an intermediate check before temporal decoding: YOLO/SPPF descriptors reached visual-only grouped linear-probe macro-F1 0.849 ± 0.015, whereas MobileNetV3 descriptors reached 0.757 ± 0.028 and preserved stronger video-identity structure (Fig. 15A,C). In the DLC-HRNet route, a coordinate-and-descriptor cache built with a validated pre-head HRNet tap reached raw-window held-out frame macro-F1 0.749 across 28 videos with accepted human annotations (Fig. 14A). The ablation, descriptor-probe, and route-level results also bound the interpretation. Pose-trained descriptors are not automatically behavior models; their value is interpretable when visual evidence, pose-derived geometry, and temporal classification are evaluated together on held-out full videos.

### 4.1. Pose checkpoints as task-specialized frozen representations

Taken together, the results change how the pose checkpoint itself is interpreted. Pose-trained descriptors can be behavior-discriminative, but their value depends on how the evidence is exposed and read. The MARS stream ablations showed that visual descriptors and pose-derived geometry were complementary, the RF/XGBoost controls further showed that flattened PCA-compressed summaries lost part of the benefit, and the MobileNetV3 and DLC-HRNet routes showed that feature quality remains empirical across backbones and pose ecosystems. A pose checkpoint therefore supplies an experimentally defined measurement object. The checkpoint, feature tap, crop definition, geometry streams, and downstream classifier together determine which behavioral evidence is recoverable.

Frozen-representation studies provide a useful analogue. Yosinski et al. showed that feature transfer depends on where a network is split and on the distance between source and target tasks. Alain and Bengio used classifier probes to measure which intermediate layers carry linearly accessible task information. Kornblith et al. showed that ImageNet-trained networks can serve as fixed feature extractors, while also showing that transfer depends on training details and target datasets (Yosinski, Clune, Bengio and Lipson, 2014; Alain and Bengio, 2016; Kornblith, Shlens and Le, 2019b). BehaviorScope-X extends this logic from object-recognition backbones to pose-trained ethology. The source task here is assay-adapted keypoint localization, and the target task is continuous behavior decoding from held-out videos drawn from the same assay distribution. The within-assay scope matters because a descriptor extracted from a pose checkpoint is tied to the camera view, arena, animal appearance, and pose-labeling regime that shaped the checkpoint. Reuse across a new assay domain would require its own calibration experiment rather than extrapolation from within-assay held-out performance.

The next representation-learning question is whether pose-adapted descriptors carry behavior-relevant evidence that differs from domain-general visual pretraining. The present experiments make that comparison worth testing because pose-specialized descriptors remained useful after coordinate export, while MobileNetV3 and DLC-HRNet showed that reusable information depended on feature-tap provenance and pose ecosystem. Descriptor content alone, however, is not the behavior model. Recovery improved when descriptors were paired with pose-derived geometry and read by a temporal classifier that preserves sequence structure. The coupling between representation and classifier leads directly to the ethological question of what kind of measurement the workflow creates.

### 4.2. Ethological stakes of reusable pose evidence

At the ethological level, reusing the same pose-and-descriptor cache across full videos turns the representation into a measurement layer. Existing behavior tools reuse different measurement units. Coordinate-centered systems such as MARS, SLEAP-based workflows, and SimBA reuse keypoints and derived geometry. Pixel-centered systems such as DeepEthogram and FERAL learn behavior-specific visual evidence directly from video. BehaviorScope-X reuses the trained pose checkpoint itself, so coordinate inspectability and pose-trained visual descriptors share the same upstream provenance. Classifier accuracy remains necessary, but the upstream evidence preserved by the model, the provenance fixed by the workflow, and the biological comparisons made auditable are also part of the measurement.

A particularly important case is an experiment that repeats the same assay across many animals or conditions. Studies comparing behavioral phenotypes across genotypes, pharmacological manipulations, developmental time points, or social contexts often face a tradeoff between manual scoring and repeated model-building. Manual scoring limits scale; repeated raw-video model training can make the visual measurement vary with the comparison being tested. In that setting, a fixed pose-and-descriptor measurement layer lets the same upstream evidence support time budgets, episode distributions, local sequence motifs, stream ablations, seed repeats, and postprocessing sensitivity analyses across the experimental grid.

Once the upstream evidence is fixed, evaluation still has to preserve the biological unit of interest. Frame accuracy is insufficient because the same number of correctly labeled frames can imply different time budgets, bout counts, episode durations, and transition structure. The MARS analyses therefore treated the ethogram as a biological object rather than as a sequence of independent frame labels. Full-video timelines and bout examples linked predictions to time budgets and episode structure (Figs. 16 and 17), bout diagnostics exposed dense-bout compression, and transition and n-gram analyses tested whether predicted timelines preserved local behavior grammar beyond class base rates (Fig. 19). The result is a more biologically specific evaluation: a classifier can be accurate at the frame level while still distorting bout persistence or sequence organization.

The same frame, bout, and sequence readouts also reveal a boundary between sustained-state measurement and event-boundary measurement. For sustained resident-intruder behaviors in the MARS held-out videos, cached pose-trained evidence can support broad episode recovery and ethogram structure. For dense short-bout structure, especially rapid investigation fragments or sub-second Fly-v-Fly aggression events, exact onset and offset recovery becomes a more delicate measurement problem. The distinction is useful for experimental design because studies centered on stable states can emphasize full-video coverage and held-out ethogram structure, whereas studies centered on brief events can plan denser annotation, boundary-tolerant metrics, shorter-window models, or earlier spatiotemporal fusion from the outset.

### 4.3. Complementary evidence and sequence-aware classifiers

A mechanistic explanation for the state/event boundary comes from how the cached streams partition behavioral evidence. The stream-ablation results give the representation a biological interpretation. Pose-self geometry and pairwise social geometry encode body configuration, distance, orientation, and relative position. Frozen visual descriptors can retain evidence that sparse landmarks do not fully capture, including contact surfaces, body overlap, occlusion, local posture, animal appearance, and scene context around interacting animals. Their complementarity predicts that descriptor gains should be largest for behaviors whose labels depend on contact, overlap, or posture details that are poorly represented by sparse landmarks, and smallest for behaviors defined primarily by distance, orientation, or locomotor trajectory.

The paired held-out ablations support that interpretation directly. Full attention decoding exceeded pose-plus-relations-only decoding by 0.031 frame macro-F1 and 0.032 bout macro-F1@0.25, while its advantage over visual-only attention was larger at 0.068 frame macro-F1 (Fig. 10). The ablation gains are meaningful on the scale of the present experiments. The 0.031 frame macro-F1 improvement over pose-plus-relations-only decoding is approximately five times the across-seed variation of the full attention model reported in the held-out MARS summary (±0.006), and the 0.032 bout macro-F1@0.25 improvement is approximately three times the corresponding full-model variation (±0.011). The larger 0.068 frame macro-F1 advantage over visual-only decoding further indicates that the visual stream does not replace pose geometry. The ablations support a fusion interpretation in which visual and geometric evidence become more useful when interpreted together.

The classifier determines whether this complementarity remains available. Random Forest and XGBoost controls received window-level inputs and dimension-reduced visual evidence, but static tabular concatenation did not preserve temporal order and stream structure in the same way as LSTM and attention heads. The RF/XGBoost comparison is therefore a control on representation use rather than a model leaderboard (Fig. 11). A direct comparison against an external behavior classifier is not straightforward here, because the MARS benchmark reports frame-level agreement with human annotators rather than a held-out classifier score under a shared protocol; the Random Forest and XGBoost controls therefore provide the pose-feature supervised reference most comparable to standard tools such as SimBA (Goodwin et al., 2024), evaluated under matched held-out videos and postprocessing. Amortized pose vision is best understood as a coupled design that joins a cached pose-trained representation, explicit geometric streams, and a downstream classifier that can preserve temporal structure.

Recent pose-sequence transformer work provides a useful point of reference: BehaVERT reported CalMS21 Task 1 class-averaged F1 of 0.848 ± 0.003, or 0.869 ± 0.001 with masked-keypoint pretraining, for attack, investigation, and mount classification from skeletal motion (Shin, Lee, Ko and Kim, 2026). Those scores show the strength of keypoint-only temporal transformers, but they are not directly comparable to the present held-out full-video frame macro-F1 and bout macro-F1@0.25 readouts, which include continuous ethogram reconstruction, the other class, temporal overlap of bouts, and sequence-structure diagnostics.

### 4.4. Failure modes reveal what aggregate scores hide

The evaluation problem becomes clearer once this coupled measurement design is treated as the object of evaluation. The MARS merge audit showed that dense investigation bouts could be compressed even when frame-level ethograms were strong. The Fly-v-Fly analysis showed that one-to four-frame events and annotation-boundary uncertainty could limit strict bout recovery. The DLC-HRNet analysis showed that route-inappropriate smoothing could reduce performance relative to raw-window predictions, and the MobileNetV3 analysis showed that changing the pose backbone could change bout-boundary behavior. Together, the four analyses show the limits of describing behavior classifiers only as accurate or inaccurate. A model can confuse identities, shift boundaries, merge nearby bouts, overcall common short events, degrade under a new backbone, or fail after route-inappropriate smoothing, and each failure points to a different scientific interpretation and next experiment. Table 6 turns the error patterns observed here into a diagnostic vocabulary for pose-trained ethogram inference.

**Table 6.** Failure-mode taxonomy grounded in the present held-out evaluations. Each row links a recurring error type to evidence observed in the BehaviorScope-X experiments and to a diagnostic that can be computed from held-out predictions.

| Failure mode | Evidence in the present study | Diagnostic |
| --- | --- | --- |
| Identity confusion | Representative MARS ethograms showed behavior-identity errors in pose+relations-only decoding that were corrected by full multimodal reuse; DLC-HRNet confusion profiles localized residual class errors across attack, investigation, mount, and other (Figs. 16 and 14C). | Confusion matrix, class precision and recall |
| Boundary displacement | MobileNetV3 showed larger median absolute boundary offsets for attack and mount than matched YOLO/SPPF attention-256, and Fly-v-Fly collar sweeps showed that strict event scores depended strongly on boundary tolerance (Figs. 13 and 22). | Onset-offset error, collar sweep |
| Bout merging | MARS investigation bouts were compressed: full attention-256 produced approximately 0.62 predicted investigation bouts per annotated bout, with predicted median duration approximately 2.29 times the human median (Fig. 18). | Predicted-to-true bout ratio, gap audit |
| Overcalling common short events | In Fly-v-Fly, lunge had high recall (0.927) but low precision (0.252), consistent with broadened prediction of a frequent short-bout class (Fig. 22). | High recall with low precision |
| Backbone-dependent degradation | MobileNetV3 supported reuse but trailed the primary YOLO/SPPF route by 0.037 frame macro-F1 and 0.044 bout macro-F1@0.25 for attention-256 in paired comparisons (Fig. 12). | Paired-video deltas across pose routes |
| Postprocessing mismatch | Applying the MARS smoothing recipe to DLC-HRNet reduced frame macro-F1 from 0.749 to 0.638 and bout macro-F1@0.25 from 0.670 to 0.537 (Fig. 14A). | Raw versus smoothed comparison |

Read against this taxonomy, error type changes biological interpretation. Identity confusion affects behavior labels and time budgets, boundary displacement affects event timing, bout merging affects episode counts and transition statistics, and postprocessing mismatch can create apparent biological differences from an analysis recipe. The same frame macro-F1 can therefore hide biologically opposite errors. A model that merges nearby bouts may inflate apparent episode persistence, whereas a model that overcalls short events may fragment the same behavior into many brief episodes. Reporting diagnostics that identify how a model fails, as well as how often it fails, makes those biological risks visible.

### 4.5. Feature provenance is part of measurement provenance

The diagnostic vocabulary addresses failures after a route has produced predictions, while provenance addresses how those routes create the evidence that classifiers read. Across the MobileNetV3 and DLC-HRNet experiments, feature provenance becomes part of the measurement rather than a software detail. A reusable feature tap defines the kind of information being carried forward, such as late semantic context, multi-resolution spatial detail, compressed bottleneck evidence, or head-proximal localization evidence. For pose-trained ethology, tap location, descriptor dimensionality, pose-checkpoint quality, crop definition, and postprocessing recipe function like recording location or sampling resolution in other biological measurements because they determine what the measurement can mean.

MobileNetV3 and DLC-HRNet illustrate two different provenance tests. MobileNetV3 provided a native backbone-dependence test within the same cache-and-classify design. The descriptor probes explain why that test matters biologically: the MobileNetV3 visual space carried behavior signal, yet its behavior organization was weaker than YOLO/SPPF and its video-identity structure was stronger (Fig. 15). For behavior measurement, this means that a larger descriptor can preserve usable visual evidence while also preserving recording-specific structure that does not map cleanly onto behavior labels. Descriptor size alone is therefore a poor proxy for reusable behavioral evidence. DLC-HRNet addressed a different provenance question by asking whether an established DeepLabCut SuperAnimal HRNet-W32 top-down workflow could be made coordinate-and-descriptor reusable through a validated pre-head multi-branch tap. That route reached raw-window frame macro-F1 0.749 with a validated 480-dimensional descriptor (Figs. 14A and 15). These route-specific results support a provenance view of pose-trained reuse in which the feature tap, descriptor geometry, crop definition, and pose framework define the visual evidence available to the downstream classifier.

Table 7 summarizes the feature-tap provenance fields that should be reported when extending this approach to another pose family. Without tap provenance, two behavior classifiers can appear comparable while drawing evidence from different visual abstractions, descriptor dimensions, crop definitions, or postprocessing recipes. Stating the tap in advance, reporting its descriptor size, and evaluating it under the same held-out ethogram readouts turn layer selection into a reproducible experimental choice rather than hidden software plumbing.

**Table 7.** Feature-tap provenance for pose-backbone reuse. Candidate taps should be specified by architectural position, descriptor dimensionality, and validation requirement before downstream behavior evaluation. The examples identify reportable provenance fields rather than universal layer choices.

| Pose architecture | Candidate reusable tap | Rationale and validation requirement |
| --- | --- | --- |
| YOLO-style pose | Shared neck or SPPF-adjacent features before detection/keypoint heads | Captures late visual context in the shared pose backbone before task-specific heads. YOLO11-style C2PSA or analogous attention blocks are best treated as empirical ablations rather than assumed defaults. |
| HRNet-style pose | Pooled multi-resolution branches before keypoint heads | Preserves simultaneous high- and low-resolution streams. For HRNet-W32, a semantic concatenation of pooled branches yields $32 + 64 + 128 + 256 = 480$ features per crop. |
| ResNet/FPN-style pose | Late trunk or pyramid features, such as C4/C5 or layer3/layer4, before deconvolutional heatmap prediction | Provides high-level pose-trained visual context while avoiding the final coordinate-localization head. Multiple adjacent stages can be compared when spatial precision and semantic abstraction trade off. |
| MobileNet/EfficientNet-style pose | Late inverted-residual or MBConv features, optionally paired with a mid-depth higher-resolution feature map | Gives a compact native descriptor from the pose checkpoint while retaining a candidate path for spatial detail. Descriptor dimensionality is part of the reportable provenance because it changes downstream capacity and cache size. |
| U-Net, hourglass, or SLEAP-style pose | Bottleneck features and late decoder features immediately upstream of confidence maps | Tests compressed semantic context and recovered spatial detail without using the final confidence-map output itself. Evaluating both candidates helps distinguish behaviors that depend on context from behaviors that depend on localization. |

### 4.6. Practical consequence of fixed upstream evidence

With feature provenance reported and held fixed, amortized reuse changes the cost structure of behavior-model development. In the MARS experiments, one upstream pose-trained cache supported stream ablations, LSTM and attention heads, RF/XGBoost controls, seed repeats, and postprocessing audits. A conventional raw-video workflow can entangle each of those comparisons with a new visual representation. A fixed pose-and-descriptor cache separates the downstream question from the upstream visual measurement, so the classifier, stream composition, random seed, label definition, or smoothing rule can change without rebuilding the evidence source.

The deployment benchmarks show that this separation also affects practice. End-to-end processing reached 25.4 frames/s on the RTX2080 workstation, 32.6 frames/s on the RTX4080 workstation, and 60.7 frames/s on Apple Silicon MPS under the tested configurations. The measured rates make the representation result operationally useful for continuous phenotyping runs in which multiple decoder, stream, seed, and postprocessing choices can be compared under a fixed upstream measurement. At this scale, studies that compare behavioral time budgets, episode structure, or local sequence motifs across many animals, treatments, or recording sessions can treat the visual measurement as a controlled component rather than rebuilding it for each downstream analysis.

### 4.7. The unresolved boundary between states, events, and domains

The remaining boundary concerns the point at which an engineering limitation becomes a measurement limitation. In the current MARS implementation, bout compression is consistent with the late-fusion architecture. Cached visual descriptors are extracted from a frozen spatial pose backbone, and temporal heads integrate the cached descriptors after the visual representation has already been formed. Shorter Fly-v-Fly windows improved recovery in a sub-second behavior regime, showing that the cache-and-classify design can be retuned. At the same time, one-to four-frame events and annotator disagreement at similar time scales show that some boundary errors occur near the resolution of the labels themselves.

The Fly-v-Fly stress test grounds the temporal part of the open question. Reducing the temporal window from 16 to 8 frames increased mean frame macro-F1 from 0.376 to 0.441 and recall from 0.555 to 0.627 (Fig. 22), but strict bout recovery remained bounded by events that often lasted only a few frames (Table 4). The window reduction is encouraging because the reuse design adapted to a shorter behavioral time scale. It is also cautionary because short-bout ethology asks both model and annotation to resolve events near the frame scale.

The domain-shift boundary is separate from temporal scale. The present validation tests held-out videos within assay-matched imaging and annotation conditions. A different strain, lighting condition, camera position, arena, or laboratory could change both pose quality and the visual cues captured by a feature tap. For example, an HRNet or YOLO descriptor that captures contact and occlusion in one top-view resident-intruder arena may not encode the same evidence after a change in viewpoint or animal appearance. Controlled domain-shift experiments would determine when a checkpoint can be reused as a measurement instrument and when the pose representation requires recalibration. A matched comparison between pose-adapted descriptors and domain-general visual pretraining is a related unresolved test, because the present data show that pose-trained descriptors can be reused within assay but do not establish whether they outperform generic frozen visual features. Behavioral scope is a further boundary: both validation assays involve two interacting animals recorded from above, so transfer to single-animal, larger-group, or non-social paradigms remains untested.

Temporal scale and domain shift also clarify where BehaviorScope-X sits within the broader decision space for behavior analysis. A behavior may fall in the safe regime if its identity persists across multiple windows and is supported by stable posture, contact, geometry, or appearance. Event-specific modeling becomes more appropriate when identity depends on a transient visual change, fast onset, or short gap between bouts. Existing behavior-analysis stacks mark nearby choices along this boundary. SimBA-style Random Forest classifiers read engineered pose-derived features after pose estimation, whereas joint pose-behavior systems such as IntegraPose predict frame-level behavior during pose processing. Unsupervised or expert-guided discovery stacks such as VAME and A-SOiD occupy another adjacent space by organizing pose dynamics into motifs or discovered behavior classes. BehaviorScope-X makes a different deployment contrast by coupling pose tracking, pose-trained descriptor reuse, and behavior decoding in one full-video route. The practical output is pose plus behavior rather than a serial pose-then-behavior handoff. In each case, bouts are assembled from the resulting label sequence or discovered state sequence.

The same logic gives users a practical decision rule. Coordinate-only workflows remain appropriate when body geometry, motion, and social configuration capture the biology of interest. BehaviorScope-X is most useful when coordinate and motion measurements are important but incomplete, and when a validated pose checkpoint can be reused as a controlled source of visual and geometric evidence. End-to-end video models, behavior-specific fine-tuning, or earlier spatiotemporal fusion become more appropriate when the behavior is defined by transient visual changes, fine contact timing, or event boundaries near the scale of annotation uncertainty.

The next experiments can make these decision rules empirical by linking each model change to a biological measurement question. Temporal experiments, including window-length sweeps and boundary-aware losses, would test how precisely bout onset, offset, and short gaps can be recovered for rapid behaviors. Provenance experiments, including frozen-versus-fine-tuned taps and early spatiotemporal adapters, would test whether contact timing, overlap, or brief posture transitions require behavior-specific visual updates. Annotation experiments with denser multi-annotator labels would estimate the temporal resolution at which a short-bout event is reliably defined by human observers. Domain-shift experiments across strains, lighting conditions, arenas, or laboratories would test whether a fixed pose checkpoint remains a stable measurement layer for phenotype comparisons. Together, these experiments would turn the state/event distinction and the assay-domain boundary into predictive design rules.

Viewed as a research program, BehaviorScope-X is best read as an opening argument for pose checkpoints as reusable scientific resources. The paper establishes conditions under which the resource is recoverable: an explicit feature tap, a frozen pose checkpoint, a downstream classifier that preserves temporal structure, and held-out full-video evaluation. It also gives diagnostics for when recovery fails and identifies the temporal and domain boundaries for future model families to target.

## 5. Limitations & Future Directions

### 5.1. Frozen reuse should be compared with targeted fine-tuning

BehaviorScope-X intentionally froze the pose-trained visual backbone so that the amortized pose vision principle could be tested cleanly. Freezing the backbone isolates the information already present in an assay-trained pose representation, but it does not establish that freezing is optimal for every assay or behavior class. A targeted future comparison should train matched fine-tuned variants under the same train, validation, and held-out splits, while preserving the same frame-level, bout-level, and ethogram-level readouts used here. Such a comparison would separate the additional representational benefit of behavior-specific visual updates from their computational cost, overfitting risk, and effect on model provenance.

### 5.2. Reusable feature taps require architecture-specific validation

The YOLO/SPPF, MobileNetV3, and DLC-HRNet results support pose-trained reuse across multiple implementations while showing that backbone and framework provenance matter. The present study validates a YOLO/SPPF tap, tests one native MobileNetV3 pose backbone, and validates one DLC-HRNet `semantic_concat` tap for the reported top-down MARS workflow. The DLC-HRNet result establishes that a native top-down DLC route can support the full cache-and-classify path with a validated pre-head descriptor, but it does not by itself isolate the incremental value of HRNet descriptors beyond DLC-derived keypoints and relation features. A direct DLC-internal ablation of pose/relation-only, HRNet-descriptor-only, and fused inputs remains an important next calibration step. More broadly, the study does not establish universal layer choices for other DeepLabCut configurations, SLEAP, ResNet, EfficientNet, hourglass, or future YOLO-style pose models. The rubric in Table 7 provides reportable provenance fields rather than a definitive layer map. Candidate shared-trunk or late pre-head features should be frozen, cached, fused with the same pose-derived geometry, and evaluated on held-out frame, bout, and ethogram metrics. Backbone extension should be treated as an empirical calibration step rather than an assumption transferred from YOLO/SPPF.

### 5.3. Short-bout boundaries require denser annotation and task-specific temporal models

The strongest MARS models recovered frame-level ethograms and much of the behavior-sequence structure, but dense short-bout regions were still compressed. Fly-v-Fly made this temporal limitation more explicit because some aggression bouts lasted only a few frames, and the available secondary annotation was limited to one held-out movie in the local public subset. The same public-data constraint limits the scope of the Fly-v-Fly generalization: the analysis tests behavior-label transfer and temporal-scale adaptation with a frozen assay-trained pose checkpoint, not strict unseen-video generalization of the upstream pose model.

Future work should collect denser multi-annotator labels for short-bout assays and report boundary-aware metrics alongside strict TIoU matching. Collar sweeps are useful because they connect model performance to the time scale of both behavior and annotation uncertainty. Model comparisons should then test whether short-bout recovery improves by changing the temporal head, the objective, or the representation path. One direction is a hybrid temporal head that combines TCN-style local boundary sensitivity with attention-based longer-context integration, allowing the same cached representation to read short, medium, and sustained bouts at different temporal scales (Lea et al., 2017; Abu Farha and Gall, 2019; Ding et al., 2023). A second direction is to treat cached pose-trained descriptors as variable-length temporal sequences of embeddings or multivariate time series, following encoder-agnostic temporal matching approaches such as DejaVid, which preserve temporal order and allow feature importance to vary across time without retraining the visual encoder (Ho and Madden, 2025). The connection to the present short-bout problem is that the limiting errors involve variable-duration bouts read from fixed cached descriptors, rather than only a failure to classify isolated frames. A third direction is temporal-aware supervision: onset/offset losses, duration-weighted losses, boundary collars, and logit-smoothing penalties selected by boundary-aware validation rather than frame accuracy alone. Earlier spatiotemporal variants could test short frame-stack crops, optical-flow or frame-difference channels, lightweight spatiotemporal adapters before caching, or behavior-specific fine-tuning of the late pose-backbone tap. The proposed short-bout strategies are hypotheses for the next model family, not conclusions from the present runs. They should be compared against direct frame-wise behavior-detection strategies, including SimBA-style coordinate-feature Random Forest classifiers and IntegraPose-style frame-wise behavior outputs, as well as unsupervised or expert-guided pose-dynamics stacks such as VAME and A-SOiD, under the same held-out videos and boundary-aware metrics.

### 5.4. Deployment speed is tractable but not fully optimized

The end-to-end benchmark showed that the current inference path is near-real-time or faster on the tested consumer hardware, and that the temporal sequence classifier is a minor component of runtime. Most cost remained upstream in video handling, pose tracking, crop construction, and frozen feature extraction. Future optimization should focus on batched crop construction, lower-overhead frame transfer, device-native video decoding where available, and tighter integration between pose inference and feature extraction. Throughput-focused engineering can improve speed without changing the scientific comparison between frozen feature reuse and downstream temporal decoding.

## 6. Conclusion

This work demonstrates that an assay-trained pose checkpoint can serve as more than a coordinate generator for animal behavior analysis. By freezing a validated pose backbone, caching its visual descriptors and pose-derived social geometry, and training compact temporal sequence classifiers on full-video windows, the framework operationalizes amortized pose vision as a computational design principle for full-video ethology, demonstrated here in two-animal social-interaction assays. Its pose-model-flexible implementation further turns that principle into a reproducible analysis workflow where supported pose-model families can differ in detector, backbone, feature tap, and runtime requirements while still producing compatible caches and evaluation artifacts. Across MARS YOLO/SPPF, MobileNetV3, DLC-HRNet, and Fly-v-Fly analyses, the results support a bounded conclusion. Pose-trained representations contain reusable visual and geometric evidence for continuous behavior analysis, but their utility depends on temporal decoding, annotation quality, bout duration, and backbone validation. For behavioral neuroscience, this creates a route to compare behavioral phenotypes across genotypes, disease models, treatments, and developmental stages against a stable pose-trained evidence source. In this form, pose training becomes a reusable investment that can support new behavior models, diagnostic ablations, and full-video ethogram measurements while keeping the upstream visual evidence inspectable and experimentally bounded.

## Ethics Statement

The study used only publicly available datasets and previously collected video and annotation resources. No new animal experiments were performed for this work.

## Acknowledgments

We thank Drs. Weihong Lin and Tatsuya Ogura, graduate student Sean O’Sullivan, and undergraduate researcher Shaun Salvador Andrade for valuable discussions during the conceptual development of BehaviorScope-X. We are grateful to the MARS team (Segalin et al., 2021c) and the Fly-v-Fly team (Eyjolfsdottir et al., 2014) for publicly releasing their datasets and annotation files, which made this work possible.

## Data and Code Availability

A permanent archival copy of BehaviorScope-X is available on Zenodo: Augustine, F. (2026). *BehaviorScope-X* (V1.0.0). Zenodo. **https://doi.org/10.5281/zenodo.21070316**. The latest codebase updates are available on GitHub: **https://github.com/farhanaugustine/BehaviorScope-X**. Video data are available via the original sources, including the MARS and Fly-v-Fly datasets (Segalin et al., 2021c; Eyjolfsdottir et al., 2014).

## CRediT authorship contribution statement

**Farhan Augustine:** Conceptualization, Methodology, Software, Formal analysis, Validation, Visualization, Supervision, Writing – original draft, Writing – review and editing. **Virginia Murray:** Validation, Visualization, Writing – original draft, Writing – review and editing.

## References

Abu Farha, Y., Gall, J., 2019. MS-TCN: Multi-stage temporal convolutional network for action segmentation, in: Proceedings of the IEEE/CVF Conference on Computer Vision and Pattern Recognition, pp. 3575–3584. doi:10.1109/CVPR.2019.00369.

Alain, G., Bengio, Y., 2016. Understanding intermediate layers using linear classifier probes. doi:10.48550/arXiv.1610.01644, arXiv:1610.01644.

Anderson, D.J., Perona, P., 2014. Toward a science of computational ethology. Neuron 84, 18–31. doi:10.1016/j.neuron.2014.09.005.

Arevalo, J., Solorio, T., Montes-y Gómez, M., González, F.A., 2017. Gated multimodal units for information fusion. arXiv doi:10.48550/arXiv.1702.01992, arXiv:1702.01992.

Augustine, F., O’Sullivan, S., Murray, V., Ogura, T., Lin, W., Singer, H.S., 2025. Integrapose: A unified framework for simultaneous pose estimation and behavior classification. Neuroscience 590, 1–22. doi:10.1016/j.neuroscience.2025.10.020.

Baltrušaitis, T., Ahuja, C., Morency, L.P., 2019. Multimodal machine learning: A survey and taxonomy. IEEE Transactions on Pattern Analysis and Machine Intelligence 41, 423–443. doi:10.1109/TPAMI.2018.2798607.

Bohnslav, J.P., Wimalasena, N.K., Clausing, K.J., Dai, Y.Y., et al., 2021. Deepethogram, a machine learning pipeline for supervised behavior classification from raw pixels. eLife 10, e63377. doi:10.7554/eLife.63377.

Breiman, L., 2001. Random forests. Machine Learning 45, 5–32. doi:10.1023/A:1010933404324.

Chen, T., Guestrin, C., 2016. XGBoost: A scalable tree boosting system, in: Proceedings of the 22nd ACM SIGKDD International Conference on Knowledge Discovery and Data Mining, pp. 785–794. doi:10.1145/2939672.2939785.

Choudhary, A., Geuther, B.Q., Sproule, T.J., Beane, G., Kohar, V., Trapszo, J., Kumar, V., 2025. Jax animal behavior system (jabs), a genetics-informed, end-to-end advanced behavioral phenotyping platform for the laboratory mouse. eLife 14, RP107259. doi:10.7554/eLife.107259. version of record published 2026.

Datta, S.R., Anderson, D.J., Branson, K., Perona, P., Leifer, A., 2019. Computational neuroethology: A call to action. Neuron 104, 11–24. doi:10.1016/j.neuron.2019.09.038.

Ding, G., Sener, F., Yao, A., 2023. Temporal action segmentation: An analysis of modern techniques. IEEE Transactions on Pattern Analysis and Machine Intelligence 45, 7169–7189. doi:10.1109/TPAMI.2022.3231064.

Efron, B., Tibshirani, R.J., 1993. An Introduction to the Bootstrap. Chapman & Hall/CRC. doi:10.1007/978-1-4899-4541-9.

Eyjolfsdottir, E., Branson, S., Burgos-Artizzu, X.P., Hoopfer, E.D., Schor, J., Anderson, D.J., Perona, P., 2014. Detecting social actions of fruit flies, in: European Conference on Computer Vision (ECCV), Springer. pp. 772–787. doi:10.1007/978-3-319-10605-2\_50.

Eyjolfsdottir, E.A., 2014. Detecting Actions of Fruit Flies. Master’s thesis. California Institute of Technology. URL: https://thesis.caltech.edu/8195/.

Goodwin, N.L., Choong, J.J., Hwang, S., Pitts, K., Bloom, L., et al., 2024. Simple behavioral analysis (simba) as a platform for explainable machine learning in behavioral neuroscience. Nature Neuroscience 27, 1411–1424. doi:10.1038/s41593-024-01649-9.

Harris, C., Finn, K.R., Kieseler, M.L., Maechler, M.R., Tse, P.U., 2023. Deepaction: a matlab toolbox for automated classification of animal behavior in video. Scientific Reports 13, 2688. doi:10.1038/s41598-023-29574-0.

Ho, D., Madden, S., 2025. DejaVid: Encoder-agnostic learned temporal matching for video classification. doi:10.48550/arXiv.2506.12585, arXiv:2506.12585. accepted to CVPR 2025.

Howard, A., Sandler, M., Chu, G., Chen, L.C., Chen, B., Tan, M., Wang, W., Zhu, Y., Pang, R., Vasudevan, V., Le, Q.V., Adam, H., 2019. Searching for MobileNetV3, in: Proceedings of the IEEE/CVF International Conference on Computer Vision, pp. 1314–1324.

Hsu, A.I., Yttri, E.A., 2021. B-soid, an open-source unsupervised algorithm for identification and fast prediction of behaviors. Nature Communications 12, 5188. doi:10.1038/s41467-021-25420-x.

Hu, Y., Ferrario, C.R., Maitland, A.D., et al., 2023. Labgym: Quantification of user-defined animal behaviors using learning-based holistic assessment. Cell Reports Methods 3, 100415. doi:10.1016/j.crmeth.2023.100415.

Huang, K., Han, Y., Chen, K., Pan, H., Zhao, G., Yi, W., Li, X., Liu, S., Wei, P., Wang, L., et al., 2021. A hierarchical 3D-motion learning framework for animal spontaneous behavior mapping. Nature Communications 12, 2784. doi:10.1038/s41467-021-22970-y.

Kabra, M., Robie, A.A., Rivera-Alba, M., Branson, S., Branson, K., 2013. Jaaba: interactive machine learning for automatic annotation of animal behavior. Nature Methods 10, 64–67. doi:10.1038/nmeth.2281.

Ke, J., Li, W., Pradhan, A., Markowitz, J.E., Wu, A., 2026. Behaviorvlm: Unified finetuning-free behavioral understanding with vision-language reasoning. arXiv arXiv:2603.12176. preprint.

Kornblith, S., Norouzi, M., Lee, H., Hinton, G., 2019a. Similarity of neural network representations revisited, in: Proceedings of the 36th International Conference on Machine Learning, PMLR. pp. 3519–3529. arXiv:1905.00414.

Kornblith, S., Shlens, J., Le, Q.V., 2019b. Do better ImageNet models transfer better?, in: Proceedings of the IEEE/CVF Conference on Computer Vision and Pattern Recognition (CVPR), pp. 2661–2671. doi:10.48550/arXiv.1805.08974, arXiv:1805.08974.

Lauer, J., Zhou, M., Ye, S., Menegas, W., Schneider, S., Nath, T., Rahman, M.M., et al., 2022. Multi-animal pose estimation, identification and tracking with deeplabcut. Nature Methods 19, 496–504. doi:10.1038/s41592-022-01443-0.

Lea, C., Flynn, M.D., Vidal, R., Reiter, A., Hager, G.D., 2017. Temporal convolutional networks for action segmentation and detection, in: Proceedings of the IEEE Conference on Computer Vision and Pattern Recognition, pp. 156–165. doi:10.1109/CVPR.2017.113.

Li, W., Ke, J., Wang, Y., Li, C., Wu, A., 2026. Learning when to look: On-demand keypoint-video fusion for animal behavior analysis. arXiv arXiv:2603.07279. preprint.

Luxem, K., Mocellin, P., Fuhrmann, F., et al., 2022. Identifying behavioral structure from deep variational embeddings of animal motion. Communications Biology 5, 1267. doi:10.1038/s42003-022-04080-7.

Mathis, A., Mamidanna, P., Cury, K.M., Abe, T., Murthy, V.N., Mathis, M.W., Bethge, M., 2018. Deeplabcut: markerless pose estimation of user-defined body parts with deep learning. Nature Neuroscience 21, 1281–1289. doi:10.1038/s41593-018-0209-y.

Miranda, L., Bordes, J., Pütz, B., Schmidt, M.V., Müller-Myhsok, B., 2023. DeepOF: a python package for supervised and unsupervised pattern recognition in mice motion tracking data. Journal of Open Source Software 8, 5394. doi:10.21105/joss.05394.

Pedregosa, F., Varoquaux, G., Gramfort, A., Michel, V., Thirion, B., Grisel, O., Blondel, M., Prettenhofer, P., Weiss, R., Dubourg, V., Vanderplas, J., Passos, A., Cournapeau, D., Brucher, M., Perrot, M., Duchesnay, E., 2011. Scikit-learn: Machine learning in Python. Journal of Machine Learning Research 12, 2825–2830.

Pereira, T.D., Shaevitz, J.W., Murthy, M., 2020. Quantifying behavior to understand the brain. Nature Neuroscience 23, 1537–1549. doi:10.1038/s41593-020-00734-z.

Pereira, T.D., Tabris, N., Matsliah, A., Turner, D.M., Li, J., Ravindranath, S., Papadoyannis, E.S., Normand, E., Deutsch, D.S., Wang, Z.Y., McKenzie-Smith, G.C., et al., 2022. Sleap: A deep learning system for multi-animal pose tracking. Nature Methods 19, 486–495. doi:10.1038/s41592-022-01426-1.

Segalin, C., Williams, J., Karigo, T., Hui, M., Zelikowski, M., Sun, J.J., Perona, P., Anderson, D.J., Kennedy, A., 2021a. The mouse action recognition system (mars): behavior annotation data. doi:10.22002/D1.2012.

Segalin, C., Williams, J., Karigo, T., Hui, M., Zelikowski, M., Sun, J.J., Perona, P., Anderson, D.J., Kennedy, A., 2021b. The mouse action recognition system (mars): pose annotation data. doi:10.22002/D1.2011.

Segalin, C., Williams, J., Karigo, T., Hui, M., Zelikowski, M., Sun, J.J., Perona, P., Anderson, D.J., Kennedy, A., 2021c. The mouse action recognition system (mars) software pipeline for automated analysis of social behaviors in mice. eLife 10, e63720. doi:10.7554/eLife.63720.

Shi, K., Zhang, G.W., Wang, Z., Zhang, S.K., Tao, H.W., Zhang, L.I., 2026. Trace: End-to-end temporal inference and annotation of animal behaviors from video. bioRxiv doi:10.64898/2026.04.14.718392. preprint.

Shin, S.J., Lee, J., Ko, C., Kim, D., 2026. BehaVERT: A transformer-based motion language model for decoding behavioral semantics in mice. International Journal of Computer Vision 134, 303. doi:10.1007/s11263-026-02834-y.

Skovorodnikov, P., Zhao, J., Buck, F., Kay, T., Frank, D., Koger, B., Costelloe, B., Couzin, I., Razzauti, J., 2025. Feral: A video-understanding system for direct video-to-behavior mapping. bioRxiv doi:10.1101/2025.11.16.688666. preprint.

Tillmann, J.F., Hsu, A.I., Schwarz, M.K., Yttri, E.A., 2024. A-soid, an active-learning platform for expert-guided, data-efficient discovery of behavior. Nature Methods 21, 703–711. doi:10.1038/s41592-024-02200-1.

Ultralytics, 2023. Ultralytics YOLO documentation. URL: https://docs.ultralytics.com/.softwaredocumentation. Accessed 2026-05-03.

Weinreb, C., Pearl, J.E., Lin, S., Osman, M.A.M., Zhang, L., et al., 2024. Keypoint-moseq: parsing behavior by linking point tracking to pose dynamics. Nature Methods 21, 1329–1339. doi:10.1038/s41592-024-02318-2.

Yosinski, J., Clune, J., Bengio, Y., Lipson, H., 2014. How transferable are features in deep neural networks?, in: Advances in Neural Information Processing Systems, pp. 3320–3328. doi:10.48550/arXiv.1411.1792, arXiv:1411.1792.

