## Supplemental Material for "BehaviorScope-X: reusing pose-trained visual representations for full-video ethology"

ABSTRACT

This document provides supplementary methods, figures, tables, and reproducibility details for BehaviorScope-X.

*Corresponding author
 ( Augustine)
ORCID(s): 0000-0002-8348-6039 ( Augustine); 0009-0006-0986-7657 ( Murray)

### S1. Dataset manifests, split definitions, and reproducibility provenance

The controlled comparisons used fixed dataset manifests rather than sampling videos independently for each model family. This provenance is part of the computational-biology claim: each behavior metric can be traced back to a specific biological video unit, pose checkpoint, feature cache, temporal model, and held-out evaluation mode. For MARS, the public resident-intruder held-out videos were used for the main behavior evaluation, with accepted human annotation files defining the frame-level ground truth. Training, validation, and held-out splits were recorded in the suite manifests and were reused across temporal sequence classifiers, controlled RF/XGBoost baselines, MobileNetV3 portability controls, DLC-HRNet portability controls, and ethogram-level analyses.

For Fly-v-Fly, behavior training used movies 1, 2, and 4, validation used movies 3 and 5, and held-out behavior evaluation used movies 6–10. The Fly pose checkpoint was trained on sampled pose frames from movies 1–10, but the held-out unit for Fly behavior results is the behavior annotation: behavior labels from movies 6–10 were excluded from behavior-model training and checkpoint selection. Fly annotation summary tables are stored with the analysis outputs to preserve the provenance of the held-out annotation analyses.

Each analysis records the command-line invocation, fixed configuration, resource measurements, model checkpoints, and tabular outputs. These files provide the provenance needed to reconstruct which checkpoints, feature caches, temporal heads, capacities, seeds, and evaluation modes contributed to each manuscript table or figure.

### S2. Temporal sequence classifier architecture

The temporal sequence classifiers consume fixed-length windows of cached input streams. Each window contains one or more visual streams, pose-derived geometry streams, and reliability scalars. Visual descriptors were extracted from frozen pose backbones and were not updated during behavior-classifier training. Pose and relational features were computed from the detected keypoints and bounding boxes.

Two head families were used in the main controlled temporal sequence classifier matrix. The LSTM head applies a recurrent encoder across the temporal dimension and uses the final hidden state for behavior classification. The temporal-attention head first projects each frame into a hidden representation and then learns attention weights over the window before classification. Both heads therefore receive the same cached input window but differ in how temporal evidence is pooled.

For MARS, the temporal sequence classifier matrix crossed input-stream condition, temporal head family, hidden dimension, and random seed. For MobileNetV3 portability and focused Fly-v-Fly experiments, the matrix was narrowed to the full-stream condition and hidden dimension 256 so that these analyses tested backbone or species transfer rather than repeating the full MARS capacity search.

**Table S1**
**Temporal classifier configuration and parameter ranges.** Rows summarize the manuscript-facing behavior-classifier families. Parameter counts include the temporal classifier and fusion layers used for behavior decoding; up-stream pose backbones were frozen and are tracked separately as pose-checkpoint provenance. The full configuration-level table is retained in `tables/production/model_parameters/`.

| Route | Seed configs | Architectures | Window | Visual dim/crop | Parameters |
|---|---|---|---|---|---|
| MARS YOLO/SPPF | 108 | 36 | 32 | 0 or 256 | 1.14M–14.65M |
| MARS MobileNetV3 | 6 | 2 | 32 | 960 | 4.48M–23.12M |
| DLC-HRNet top-down | 1 | 1 | 32 | 480 | 8.21M |
| Fly-v-Fly YOLO/SPPF | 2 | 2 | 8 or 16 | 256 | 4.97M |

Classifier training used train-split class weights, AdamW optimization, validation macro-F1 for checkpoint selection, and early stopping. The selected checkpoint for each run was evaluated only after training against the held-out video set.

### S3. Input streams and feature construction

BehaviorScope-X represents each social frame through four stream families: group visual context, per-animal visual descriptors, pose-self geometry, and pairwise relational geometry. The group visual stream is extracted from a crop containing both animals. Per-animal visual streams are extracted from animal-centered crops. Pose-self features summarize each animal's normalized keypoint geometry, while pairwise relational features summarize inter-animal distance, orientation, relative motion, and overlap.

The MARS YOLO/SPPF feature tap produces a 256-dimensional descriptor per crop. With one group crop and two animal-centered crops, the raw visual feature dimension is 768 before any stream ablation. The MARS pose-derived geometry vector contributes 132 dimensions. MobileNetV3 supplies a larger native visual descriptor, 960 dimensions per crop, and is therefore reported as a cross-backbone portability control rather than as a size-matched architecture competition. The DLC-HRNet portability analysis uses a documented HRNet-W32 pooled multi-branch feature tap. Four multi-resolution HRNet branches are global-average-pooled and concatenated, giving $32+64+128+256 = 480$ visual dimensions per crop and 1,440 visual dimensions per frame for one group crop and two animal-centered crops.

The relational overlap feature is defined as

$$f_{\text{overlap}} = \frac{\text{area}(\mathbf{b}_i^t \cap \mathbf{b}_j^t)}{\min(a_i^t, a_j^t)}. \tag{S1}$$

For valid boxes this quantity is bounded by 1.0. The implementation includes an epsilon-protected denominator for numerical stability, but no upper cap is part of the mathematical feature definition.

### S4. Feature caching and manifest format

The amortized-reuse design depends on caching pose-model outputs before training behavior classifiers. Each cache records the split, video identity, window start and end frames, crop-derived visual descriptors, pose-derived geometry, labels, and metadata needed to reconstruct the evaluation set.

For held-out evaluation, caches are built independently of training windows. This prevents downstream temporal heads from changing the pose checkpoint, crop definitions, split boundaries, or cached visual features. MARS held-out evaluation used the same held-out video manifests for all compared temporal heads. Fly-v-Fly held-out evaluation used movies 6–10 with stride 4 caches for both 16-frame and 8-frame window lengths.

The cache files preserve analysis provenance: table outputs record the model name, stream condition, head family, capacity, seed, manifest path, feature-cache path, and evaluation mode whenever these fields are needed to trace a reported metric back to the underlying run. The DLC held-out cache was stored separately from the YOLO/MobileNetV3 caches because it was built from full top-down DLC detector+pose inference. Its feature cache stores the 480-dimensional pooled HRNet descriptor per crop in float16 NPZ files.

### S5. Postprocessing of window predictions

Window-level class probabilities are converted into full-video frame predictions by assigning each frame the average probability over all windows covering that frame. The resulting per-frame probability trace is converted to labels by argmax or by the prespecified threshold decoder when such a decoder is enabled. The main controlled comparisons used argmax decoding for matched evaluation.

Frame labels are then optionally smoothed with a one-dimensional median filter and a minimum-duration filter. MARS used a median filter of width 5 and removed predicted behavior bouts shorter than 15 frames. Fly-v-Fly used a median filter of width 3 and minimum predicted bout duration of 3 frames, reflecting the much shorter duration of fly aggression bouts.

Raw and smoothed outputs are retained when diagnosing bout compression. This separates limitations arising from the model's predicted sequence from limitations introduced by postprocessing.

### S6. Frame-level and bout-level evaluation

Frame-level evaluation reports accuracy, macro precision, macro recall, and macro F1. Macro metrics are emphasized because the other/background class can dominate frame counts in full-video recordings.

**Table S2**
**MARS ground-truth bout counts and durations.** Ground-truth bouts were computed from dense frame labels after applying the same class-priority convention used for BehaviorScope-X held-out evaluation. The “all” scope includes all 132 annotated MARS videos (`train`, `validation`, `test_1`, and `test_2`); the held-out scope includes the 30 public held-out videos in `test_1` and `test_2`. Durations show that MARS behavior bouts, especially investigation, are often shorter than one second. IQR denotes the 25th–75th percentile range.

| Scope | Behavior | Bouts | Mean (s) | Median (s) | IQR (s) | $< 1$s (%) | $< 2$s (%) |
|---|---|---|---|---|---|---|---|
| All | All behavior | 7192 | 2.56 | 1.10 | 0.50–2.63 | 46.1 | 67.2 |
| All | Attack | 927 | 2.39 | 1.47 | 0.63–3.13 | 37.1 | 60.9 |
| All | Investigation | 5518 | 2.23 | 0.93 | 0.43–2.23 | 51.5 | 71.7 |
| All | Mount | 747 | 5.26 | 2.43 | 1.28–5.20 | 16.9 | 41.6 |
| Held-out | All behavior | 2070 | 2.17 | 0.88 | 0.40–2.23 | 53.5 | 71.5 |
| Held-out | Attack | 356 | 1.79 | 1.30 | 0.53–2.47 | 41.9 | 67.4 |
| Held-out | Investigation | 1499 | 1.75 | 0.73 | 0.33–1.77 | 61.0 | 77.5 |
| Held-out | Mount | 215 | 5.77 | 2.50 | 1.22–5.62 | 20.5 | 36.3 |

Bout-level evaluation converts each non-background frame sequence into maximal contiguous same-class bouts. A predicted bout is eligible to match a ground-truth bout only if the classes agree and their temporal intersection-over-union exceeds the specified threshold:

$$\mathrm{tIoU}(p, g) = \frac{|p \cap g|}{|p \cup g|}. \tag{S2}$$

Bout precision, recall, and F1 are then computed from matched predictions, unmatched predictions, and unmatched ground-truth bouts. MARS summary tables report tIoU thresholds 0.25 and 0.50. Fly-v-Fly also reports tIoU 0.10 because some fly aggression bouts last only a few frames.

The statistical unit for held-out model comparison is the video or the video-seed pair, not the overlapping window. Window counts describe training and prediction generation but do not define independent experimental samples.

### MARS ground-truth bout timing

To contextualize bout compression, MARS ground-truth bout timing was computed directly from the BENTO annotation files in `MARS-data/train`, `MARS-data/validation`, `MARS-data/test_1`, and `MARS-data/test_2`. The parser used the same class set and priority convention as the held-out evaluator: `attack`, `investigation`, `mount`, and `other`. Annotation spans were expanded to dense 30-Hz frame labels, and non-`other` frame runs were then converted back into maximal ground-truth behavior bouts. Latencies are reported in seconds. For adjacent non-`other` bouts, latency is the number of background frames between the end of one bout and the start of the next, divided by the annotation frame rate; zero indicates frame-adjacent or priority-resolved contact between successive behavior bouts. Same-class recurrence measures the interval to the next bout of the same behavior, allowing other behaviors to occur between the two same-class bouts.

**Table S3**

**Overall MARS ground-truth interbout latency.** Rows summarize different bout-to-bout timing definitions. "Any adjacent" uses every consecutive non-`other` bout pair in the dense ethogram. "Immediate same behavior" is restricted to adjacent pairs with the same class. "Immediate interbehavior" is restricted to adjacent pairs where the class changes. "Same-class recurrence" measures the gap to the next bout of the same behavior even if intervening behaviors occur. The held-out MARS ethograms contain many subsecond bout intervals, so bout-boundary preservation is intrinsically stringent even before model error or postprocessing is considered.

| Scope | Timing definition | Pairs | Mean (s) | Median (s) | IQR (s) | $< 1$s (%) | $< 2$s (%) | $< 5$s (%) |
|---|---|---|---|---|---|---|---|---|
| All | Any adjacent | 7060 | 4.21 | 1.23 | 0.03–5.07 | 46.2 | 58.3 | 74.6 |
| All | Same adjacent | 4067 | 6.25 | 3.23 | 1.23–7.88 | 19.6 | 37.6 | 62.2 |
| All | Interbehavior | 2993 | 1.44 | 0.03 | 0.03–0.07 | 82.2 | 86.5 | 91.4 |
| All | Same-class recurrence | 6927 | 8.65 | 3.43 | 1.30–9.20 | 18.1 | 36.0 | 59.9 |
| Held-out | Any adjacent | 2040 | 3.58 | 0.77 | 0.03–4.20 | 52.7 | 63.8 | 78.1 |
| Held-out | Same adjacent | 960 | 6.06 | 3.27 | 1.13–7.83 | 21.1 | 38.0 | 62.1 |
| Held-out | Interbehavior | 1080 | 1.37 | 0.03 | 0.03–0.33 | 80.8 | 86.7 | 92.3 |
| Held-out | Same-class recurrence | 2007 | 8.08 | 3.27 | 1.23–8.52 | 19.6 | 36.5 | 61.9 |

**Table S4**

**Same-behavior ground-truth latency by class.** Immediate same-behavior latency counts only adjacent bout pairs with the same class. Same-class recurrence measures the gap to the next bout of the same class regardless of whether other behaviors intervene. The recurrence rows are therefore the more general measure of how rapidly the same annotated behavior returns within the ethogram, while the immediate rows describe literal same-label fragmentation in the adjacent bout stream.

| Scope | Definition | Behavior | Pairs | Mean (s) | Median (s) | IQR (s) | $< 1$s (%) | $< 2$s (%) |
|---|---|---|---|---|---|---|---|---|
| All | Immediate | Attack | 137 | 1.35 | 0.60 | 0.40–1.23 | 70.1 | 86.1 |
| All | Immediate | Investigation | 3917 | 6.44 | 3.47 | 1.33–8.13 | 17.9 | 35.8 |
| All | Immediate | Mount | 13 | 1.75 | 1.67 | 1.60–2.13 | 7.7 | 69.2 |
| All | Recurrence | Attack | 876 | 10.38 | 1.87 | 0.83–5.67 | 29.7 | 51.8 |
| All | Recurrence | Investigation | 5386 | 6.60 | 3.32 | 1.33–8.10 | 17.5 | 36.2 |
| All | Recurrence | Mount | 665 | 22.96 | 11.33 | 4.40–27.93 | 7.7 | 14.3 |
| Held-out | Immediate | Attack | 24 | 1.05 | 0.47 | 0.29–1.27 | 66.7 | 95.8 |
| Held-out | Immediate | Investigation | 935 | 6.19 | 3.40 | 1.20–7.90 | 20.0 | 36.5 |
| Held-out | Immediate | Mount | 1 | 1.67 | 1.67 | 1.67–1.67 | 0.0 | 100.0 |
| Held-out | Recurrence | Attack | 342 | 9.30 | 1.93 | 0.81–5.68 | 29.5 | 50.9 |
| Held-out | Recurrence | Investigation | 1469 | 6.21 | 3.23 | 1.23–7.30 | 19.1 | 36.6 |
| Held-out | Recurrence | Mount | 196 | 19.90 | 11.23 | 4.93–25.15 | 6.1 | 11.2 |

### S7. Ethogram-level summaries

Ethogram analyses compare model predictions and human annotations as ordered behavior sequences over complete videos. The main summaries include behavior budgets, bout counts, bout-duration ratios, bout-to-bout transition matrices, and n-gram recovery. Before transition analysis, contiguous bouts of the same class are collapsed so that the transition matrix reflects behavior changes rather than frame persistence.

For sequence-structure analyses, the ordered non-background bout sequence is converted into behavior n-grams. Model n-gram recovery is interpreted against null sequences that preserve composition or first-order transition statistics. These comparisons separate recovery of local ethogram grammar from simple base-rate matching.

**Table S5**
**Held-out ordered interbehavior latency in MARS ground truth.** Rows report immediate adjacent non-`other` bout pairs in `test_1` and `test_2` where the behavior class changes. The ordered direction is retained because transitions are asymmetric. Many investigation-to-attack and investigation-to-mount transitions occur with one-frame or zero-frame gaps, showing that the held-out annotations often place distinct behavior bouts in direct temporal succession. Rare transition types are retained for completeness but should not be overinterpreted.

| Transition | Pairs | Mean (s) | Median (s) | IQR (s) | < 1s (%) | < 2s (%) | < 5s (%) |
|---|---|---|---|---|---|---|---|
| Attack → Investigation | 330 | 2.00 | 0.48 | 0.03–1.43 | 65.2 | 80.9 | 91.2 |
| Investigation → Attack | 328 | 0.07 | 0.03 | 0.03–0.03 | 99.4 | 99.4 | 99.4 |
| Investigation → Mount | 212 | 0.08 | 0.03 | 0.03–0.03 | 99.5 | 99.5 | 99.5 |
| Mount → Investigation | 207 | 3.76 | 0.03 | 0.03–4.57 | 57.0 | 62.3 | 75.4 |
| Attack → Mount | 2 | 0.03 | 0.03 | 0.03–0.03 | 100.0 | 100.0 | 100.0 |
| Mount → Attack | 1 | 0.03 | 0.03 | 0.03–0.03 | 100.0 | 100.0 | 100.0 |

Bout-compression diagnostics are retained in the supplement because a model can achieve strong frame-level performance while under-counting dense short-bout structure. This is particularly important for investigation in MARS and for short Fly-v-Fly aggression behaviors.

### S8. YOLO26n-pose backbone and SPPF feature tap

The primary BehaviorScope-X representation source is a YOLO26n-pose checkpoint trained for MARS keypoint localization. During behavior training, this checkpoint is frozen. It supplies detections, keypoints, crop coordinates, pose-derived geometry, and intermediate visual descriptors.

The YOLO/SPPF feature tap uses the frozen backbone through the SPPF-containing layer boundary, followed by adaptive average pooling. For a 224-by-224 crop, this yields one 256-dimensional descriptor per crop. The downstream behavior classifier therefore receives a compact learned visual representation without training a second video backbone for behavior recognition.

SPPF aggregates multi-scale spatial context before the feature vector is pooled. This is useful for behavior analysis because crop-local evidence can span both small local features and whole-body or social configuration cues. The C2PSA module participates in full pose training but is not the feature extraction endpoint used in the primary YOLO/SPPF behavior representation.

### S9. Multi-scale feature extraction comparison

The multi-scale feature extraction analysis compared feature taps from different backbone depths and aggregation strategies. Its purpose was to test whether the SPPF-level descriptor was a reasonable feature source for amortized reuse, balancing behavior performance, feature dimensionality, cache size, and runtime cost.

In matched feature-tap selection trials, replacing the SPPF descriptor with a multi-scale tap that concatenated features from multiple backbone depths produced only a marginal change in validation macro-F1 (SPPF 0.814 and

0.818 versus multi-scale 0.816 and 0.828 at LSTM hidden dimensions 256 and 896, respectively). Because the multi-scale tap increased descriptor dimensionality and cache size without a corresponding behavior gain, the compact SPPF descriptor was retained as the default feature source.

The final manuscript promotes the SPPF-tap route because it provides a compact 256-dimensional descriptor per crop while retaining late-backbone spatial context from the trained pose model. Earlier multi-scale trials were used as feature-tap selection checks rather than as independent biological experiments. For the release supplement, the reportable provenance is therefore the chosen tap location, descriptor dimensionality, frozen-checkpoint identity, and the downstream controlled matrix built from that fixed cache. The controlled YOLO/SPPF stream-ablation and training-throughput figures below serve as the manuscript-facing validation that the selected SPPF descriptor supports the downstream behavior-decoding workflow.

#### Additional YOLO/SPPF analyses

The following supplemental figures provide methodological context for the main YOLO/SPPF results. The full matrix figure shows the controlled comparison underlying the streamlined main-text stream-ablation panels. The ethogram and bout-structure figures clarify why strong frame-level decoding can coexist with compressed bout counts and altered transition matrices in dense short-bout regimes. The training-throughput figure summarizes the resource profile of model fitting; end-to-end deployment throughput is reported separately in Section S14.

### S10. Controlled RF/XGBoost baseline details

The RF/XGBoost baselines test whether tabular classifiers can exploit the same pose-derived and frozen visual evidence when temporal access is controlled. Pose-frame models receive single-frame pose features. Pose-window models receive temporally concatenated pose features over the same local context used by the temporal sequence classifiers. Visual descriptor baselines use train-split PCA before RF/XGBoost training so that dimensionality reduction does not leak held-out information.

These models are interpreted as controlled classical baselines rather than as direct replacements for temporal sequence classifiers. Their role is to test the practical limit of tabular classifiers under matched temporal access and train-only dimensionality reduction.

### S11. MobileNetV3 portability details

The MobileNetV3 analysis tests whether the pose-reuse principle is restricted to the YOLO/SPPF feature tap. The MobileNetV3 checkpoint is a native pose backbone with a 960-dimensional descriptor per crop, yielding 2,880 raw visual dimensions per frame for one group crop and two animal-centered crops.

Since MobileNetV3 differs from YOLO/SPPF in architecture, feature dimension, pose quality, and extraction layer, the comparison is interpreted as a portability control. It asks whether a structurally different pose-trained backbone can support the same full-video behavior decoding workflow, not whether two size-matched architectures are equivalent.

Seed-level MobileNetV3 behavior metrics, cache provenance, pose metrics, and resource details are retained in the manuscript-facing production tables so the main text can focus on the cross-backbone finding.

The final descriptor-provenance analysis further qualifies this portability result by showing that the larger MobileNetV3 descriptor retained strong video-identity structure. The MobileNetV3 route should therefore be read as a successful cache-and-classify portability test, not as evidence that larger frozen visual descriptors are automatically better behavioral measurements.

### S12. DeepLabCut SuperAnimal HRNet-W32 portability details

The DLC-HRNet analysis tests whether amortized pose vision can be implemented through a widely used external pose-estimation framework. The run used DeepLabCut 3.0.0rc10 with the SuperAnimal TopViewMouse HRNet-W32 model in a top-down detector-plus-pose workflow. The pose model was fine-tuned on the MARS DLC project and selected by validation pose mAP. Epoch 27 produced the best validation pose mAP during fine-tuning (91.768) and was therefore used for the behavior cache. The detector was fine-tuned within the same DLC project; epoch 4 produced the best validation detector mAP50–95 (71.022) and was therefore used for top-down held-out inference.

Held-out DLC evaluation used the same 28 accepted MARS videos used by the corresponding YOLO/SPPF and MobileNetV3 held-out analyses. The full-video DLC top-down NPZ cache contained 11,960 windows and occupied 39.08 GiB. The corresponding HRNet feature cache occupied 1.04 GiB. This separation preserves two distinct provenance layers: the full top-down detector+pose cache can be audited independently of the smaller behavior-classifier feature cache.

For visual feature reuse, features were extracted from the HRNet feature module inside the DLC pose model immediately before the keypoint heads. Four HRNet-W32 branches are globally average-pooled and concatenated, giving $32 + 64 + 128 + 256 = 480$ visual dimensions per crop. With one group crop and two animal-centered crops, this yields 1,440 visual dimensions per frame before temporal windowing. The held-out behavior classifier used the same attention-256 temporal head and held-out evaluation metrics as the primary MARS held-out comparison.

### S13. Visual descriptor provenance details

The descriptor-provenance analysis compares the frozen visual evidence exposed by the YOLO/SPPF, MobileNetV3, and DLC-HRNet pose routes before temporal classification. Each route used the same 28 accepted held-out

**Table S6**
**Route-level visual descriptor provenance summary.** Fisher ratios and linear-probe metrics were computed from balanced held-out descriptor samples before temporal classification. Effective dimensionality is the participation ratio over the top 128 principal components.

| Route | Features | Fisher ratio | Eff. dim. | PC50 | Probe macro-F1 | Probe acc. |
|---|---|---|---|---|---|---|
| YOLO/SPPF | 768 | 54.0 | 14.6 | 6 | $0.849 \pm 0.015$ | $0.859 \pm 0.018$ |
| MobileNetV3 | 2880 | 26.1 | 7.6 | 4 | $0.757 \pm 0.028$ | $0.767 \pm 0.033$ |
| DLC-HRNet | 1440 | 63.3 | 13.3 | 5 | $0.704 \pm 0.059$ | $0.728 \pm 0.046$ |

**Table S7**
**Descriptor component probes and video-identity structure.** Behavior macro-F1 was computed from visual descriptors alone. Video balanced accuracy uses the same descriptor components to predict held-out video identity across 28 videos; chance is 0.036.

| Route | Component | Features | Behavior macro-F1 | Video bal. acc. | Behavior–video acc. |
|---|---|---|---|---|---|
| YOLO/SPPF | Group | 256 | 0.663 | 0.507 | 0.159 |
| YOLO/SPPF | Animal pair | 512 | 0.824 | 0.603 | 0.221 |
| YOLO/SPPF | Combined | 768 | 0.828 | 0.594 | 0.233 |
| MobileNetV3 | Group | 960 | 0.684 | 0.709 | -0.021 |
| MobileNetV3 | Animal pair | 1920 | 0.715 | 0.773 | -0.053 |
| MobileNetV3 | Combined | 2880 | 0.750 | 0.746 | 0.006 |
| DLC-HRNet | Group | 480 | 0.621 | 0.278 | 0.341 |
| DLC-HRNet | Animal pair | 960 | 0.677 | 0.321 | 0.356 |
| DLC-HRNet | Combined | 1440 | 0.730 | 0.338 | 0.390 |

MARS videos and the same four behavior classes. For each sampled 32-frame window, group and animal-centered visual descriptors were averaged across time to form a window-level descriptor vector. Route-level analyses used 2,000 balanced windows, with 500 windows per behavior class. Component and video-identity probes used 1,200 balanced windows, with 300 windows per behavior class.

The route-level probe asks how much behavior-label structure is linearly accessible from frozen descriptors alone. The video-identity probe uses the same descriptors to estimate how much recording-specific structure remains in the frozen visual space. This distinction matters for measurement provenance: a descriptor can support behavior decoding while also preserving nuisance structure tied to a particular recording, pose route, or visual background.

Linear centered-kernel alignment showed route-specific descriptor geometry: YOLO/SPPF and MobileNetV3 had the highest pairwise alignment (CKA = 0.779), whereas DLC-HRNet aligned more weakly with YOLO/SPPF (CKA = 0.376) and MobileNetV3 (CKA = 0.218). These results support the main-text conclusion that descriptor reuse is a provenance-sensitive measurement choice rather than a framework-independent black box.

### S14. End-to-end deployment benchmark implementation and diagnostics

We evaluated the BehaviorScope-X inference path using the benchmark package in `end-to-end_pipeline_benchma`[illegible] The bundle was designed to be moved between computers without editing absolute paths. It contains three held-out MARS MP4 videos, their matching `.annot` files for traceability, the MARS YOLO-pose checkpoint, and two selected full-stream temporal sequence classifiers: LSTM-256 and attention-256, both from seed 42. Each benchmark run loads the pose model and one temporal classifier, processes a full MP4 recording, writes frame/window behavior predictions, and records runtime measurements.

The measured path is the same Python inference route used for deployment: video loading, YOLO-pose detection and tracking, crop construction, frozen YOLO/SPPF feature extraction, temporal sequence classification, postprocessing, CSV export, and resource monitoring. The default decoding path uses OpenCV/Ultralytics on the original MP4 files. An optional `ffmpeg-predecode` mode first writes a local cached MP4 and then runs the same Python/PyTorch inference path on that cached file. This mode tests whether video preparation contributes to throughput limits, but it is not a zero-copy NVDEC, NVENC, or VideoToolbox frame provider. Cache construction time is therefore reported separately from inference wall time.

Across the six model-video runs, the standard OpenCV path processed 125,152 frames at 60.7 frames/s on Apple Silicon MPS, 25.4 frames/s on an RTX2080 workstation, and 32.6 frames/s on the RTX4080 workstation (Supplementary Fig. S9; Supplementary Table S8). Thus, the implementation was near-real-time on the RTX2080 and real-time or faster on the RTX4080 and Apple Silicon systems.

The RTX4080 FFmpeg-predecode/cache diagnostic processed the same 125,152 frames at 31.8 frames/s during inference, or 31.6 frames/s when the unique per-video cache construction time was included once. This did not improve over the standard RTX4080 OpenCV path; per-video/model throughput changed by -1.6% to -2.9%. The result indicates that MP4 predecode/cache alone does not explain the discrete-GPU throughput gap in this implementation.

Stage timing in the RTX2080 OpenCV run and RTX4080 diagnostic localized the remaining cost upstream of temporal sequence classification (Supplementary Fig. S10). The combined pose/tracking/crop stage accounted for 62.5% of interval wall time on the RTX2080 OpenCV run and 73.8% on the RTX4080 FFmpeg-cache diagnostic. Frozen YOLO/SPPF feature extraction accounted for 30.1% and 19.1%, respectively; window assembly accounted for approximately 6%; and temporal classification accounted for 0.62–0.64%. The reported pose/tracking/crop category includes video-frame retrieval through the local inference path, Ultralytics pose/tracking latency, transfer of pose results to host memory, and crop construction. These timings identify the practical optimization target as the frame, pose, crop, and feature-extraction path rather than the LSTM or attention classifier.

**Table S8**
**End-to-end benchmark summary.** Aggregate FPS is computed as total frames divided by total inference wall time. For FFmpeg-predecode/cache, cache construction time is reported separately and included once per unique source video in the final column.

| Condition | Runs | Frames | Inference FPS | FPS incl. cache | Mean GPU util. |
|---|---|---|---|---|---|
| MPS OpenCV | 6 | 125152 | 60.7 | 60.7 | – |
| RTX2080 OpenCV | 6 | 125152 | 25.4 | 25.4 | 23.7% |
| RTX4080 OpenCV | 6 | 125152 | 32.6 | 32.6 | 31.4% |
| RTX4080 FFmpeg cache | 6 | 125152 | 31.8 | 31.6 | 32.1% |

**Table S9**
**Focused Fly-v-Fly primary-annotation held-out evaluation.** Metrics summarize movies 6–10 under primary annotations. Raw-window metrics are reported for frame macro precision, recall, F1, and bout F1 at TIoU 0.25. Smoothed metrics report bout F1 at TIoU 0.10, 0.25, and 0.50.

| Window | Frame P | Frame R | Frame F1 | Raw bout F1@0.25 | Smooth F1@0.10 | Smooth F1@0.25 | Smooth F1@0.50 |
|---|---|---|---|---|---|---|---|
| 16 frames / stride 8 | 0.386 | 0.555 | 0.376 | 0.237 | 0.320 | 0.237 | 0.163 |
| 8 frames / stride 4 | 0.412 | 0.627 | 0.441 | 0.374 | 0.446 | 0.374 | 0.214 |

Together, these measurements characterize the tractability of the current deployment path and identify the dominant runtime costs that remain available for future optimization.

### S15. Fly-v-Fly annotation-aware evaluation details

Fly-v-Fly held-out evaluation uses primary annotations for movies 6–10 and secondary-annotation analyses where a second annotation file is available. In the local Aggression subset, the held-out secondary annotation is `movie6_second_actions.mat`. Videos without secondary annotation are retained in the annotation-availability summaries rather than omitted.

For videos with secondary annotation, the evaluator reports primary, secondary, union, intersection, and padded-primary ground-truth modes. The union mode treats a behavior frame as present if either annotator marked it. The intersection mode retains only same-class agreement. The padded-primary mode expands primary bouts by an empirical boundary margin estimated from primary-versus-secondary bout-boundary offsets.

The final focused Fly-v-Fly evaluation compared two full-stream attention-256 YOLO/SPPF temporal recipes. The 8-frame window with stride 4 improved held-out primary-annotation frame macro-F1 and bout F1 relative to the 16-frame window, consistent with the short duration of the held-out aggression bouts (Supplementary Table S9). The median held-out lunge bout was four frames at 30 Hz, and charge events could be one to four frames, making strict frame-exact event recovery an intentionally severe test.

Annotation-aware scoring was retained to separate model error from boundary ambiguity. For the held-out video with secondary annotation, the same predictions were scored against primary, secondary, union, intersection, and padded-primary ground-truth modes. The 8-frame recipe remained the stronger short-window configuration under

**Table S10**
**Fly-v-Fly annotation-mode sensitivity.** Rows summarize raw frame macro-F1 and raw bout F1@0.25 for the two focused window recipes under alternative ground-truth modes. Primary and padded-primary rows use movies 6–10; secondary, union, and intersection rows use the held-out movie with secondary annotations available.

| Window | Ground-truth mode | Frame macro-F1 | Bout F1@0.25 |
|---|---|---|---|
| 16 frames / stride 8 | Primary | 0.376 | 0.237 |
| 16 frames / stride 8 | Padded primary | 0.393 | 0.304 |
| 16 frames / stride 8 | Secondary | 0.312 | 0.146 |
| 16 frames / stride 8 | Union | 0.376 | 0.145 |
| 16 frames / stride 8 | Intersection | 0.340 | 0.143 |
| 8 frames / stride 4 | Primary | 0.441 | 0.374 |
| 8 frames / stride 4 | Padded primary | 0.335 | 0.172 |
| 8 frames / stride 4 | Secondary | 0.360 | 0.243 |
| 8 frames / stride 4 | Union | 0.442 | 0.301 |
| 8 frames / stride 4 | Intersection | 0.380 | 0.308 |

primary labels and also improved the intersection and union annotation modes relative to the 16-frame recipe (Supplementary Table S10). The padded-primary mode expands primary bouts by empirically estimated boundary uncertainty and therefore tests boundary tolerance rather than strict primary-label recovery.

### S16. GUI application and annotation workflow

The BehaviorScope-X application provides a graphical route through the same command-line workflow used for the manuscript analyses. The application organizes project setup, video import, pose tracking, annotation, dataset export, training, evaluation, and full-video inference into sequential tabs. Each GUI action constructs the corresponding command-line invocation so that runs launched through the interface remain reproducible.

The annotation interface stores labeled spans with review states. Only approved spans are exported for training. The application supports class-folder clip export for clip-based workflows and full-video BENTO-compatible `.annot` export for complete ethogram review. After training, the application can export a bundled model containing the temporal classifier and the associated pose checkpoint, simplifying deployment on new recordings.

The following screenshots show the user-facing workflow used to construct a reproducible BehaviorScope-X analysis from tutorial annotations through single-video and batch inference. The numbered callouts in each panel identify the fields that determine the corresponding command-line invocation.

### CRediT authorship contribution statement

**Farhan Augustine:** Conceptualization, Methodology, Software, Formal analysis, Validation, Visualization, Supervision, Writing – original draft, Writing – review and editing. **Virginia Murray:** Validation, Visualization, Writing – original draft, Writing – review and editing.

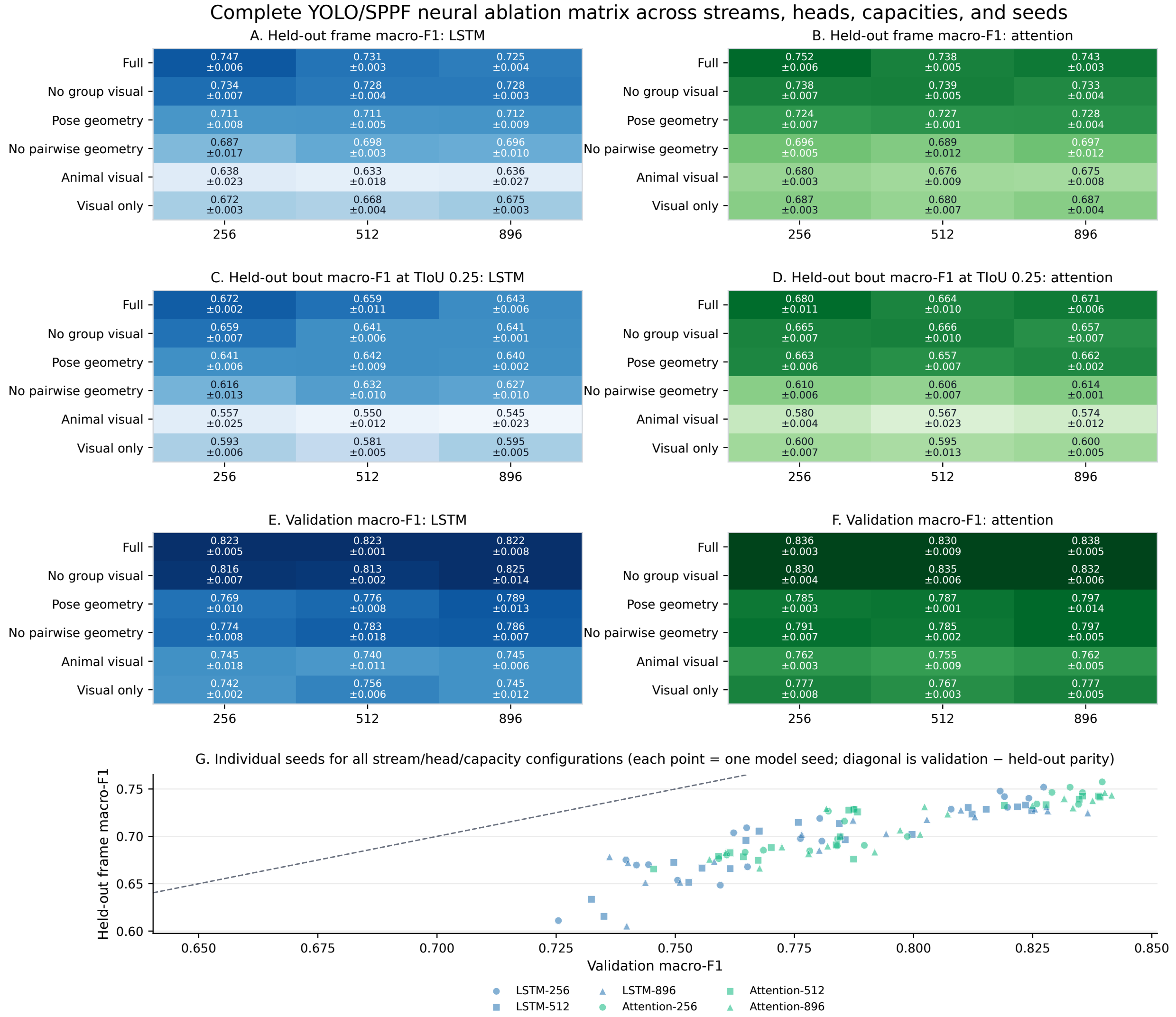

**Figure S1: Full YOLO/SPPF stream-ablation matrix.** Held-out and validation metrics are shown across stream conditions, temporal head families, capacities, and seeds. The main text reports the streamlined stream-ablation and temporal-head findings; this figure shows the full controlled matrix, including validation-to-held-out behavior across individual seed runs.

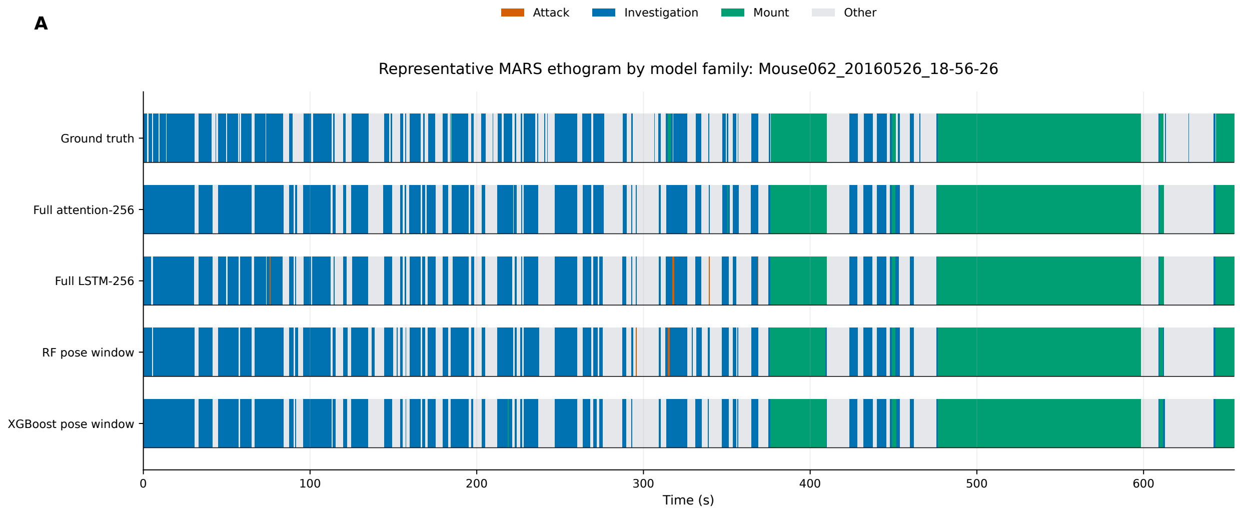

**Figure S2: Representative MARS ethograms by model type.** Example full-video predictions compare model families and stream conditions against human annotation. The purpose is qualitative but specific: to show how model-family and stream choices appear in continuous ethograms after the quantitative held-out metrics have identified the relevant model comparisons.

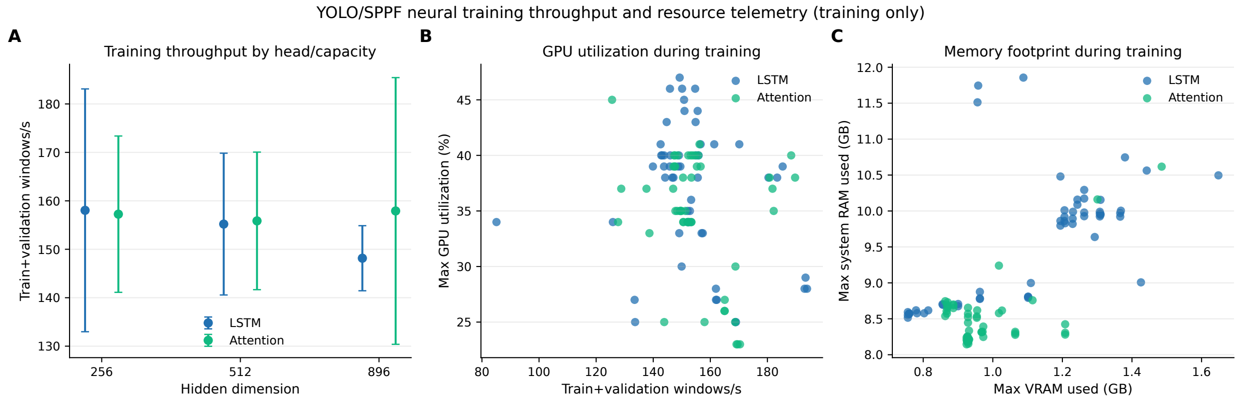

**Figure S3: Training-stage throughput and resource telemetry.** Training and validation throughput, GPU utilization, memory use, and wall time summarize the computational profile of the controlled YOLO/SPPF temporal sequence classifier runs. These values describe training-stage resource use only; deployment throughput is measured separately with the end-to-end benchmark in Section S14.

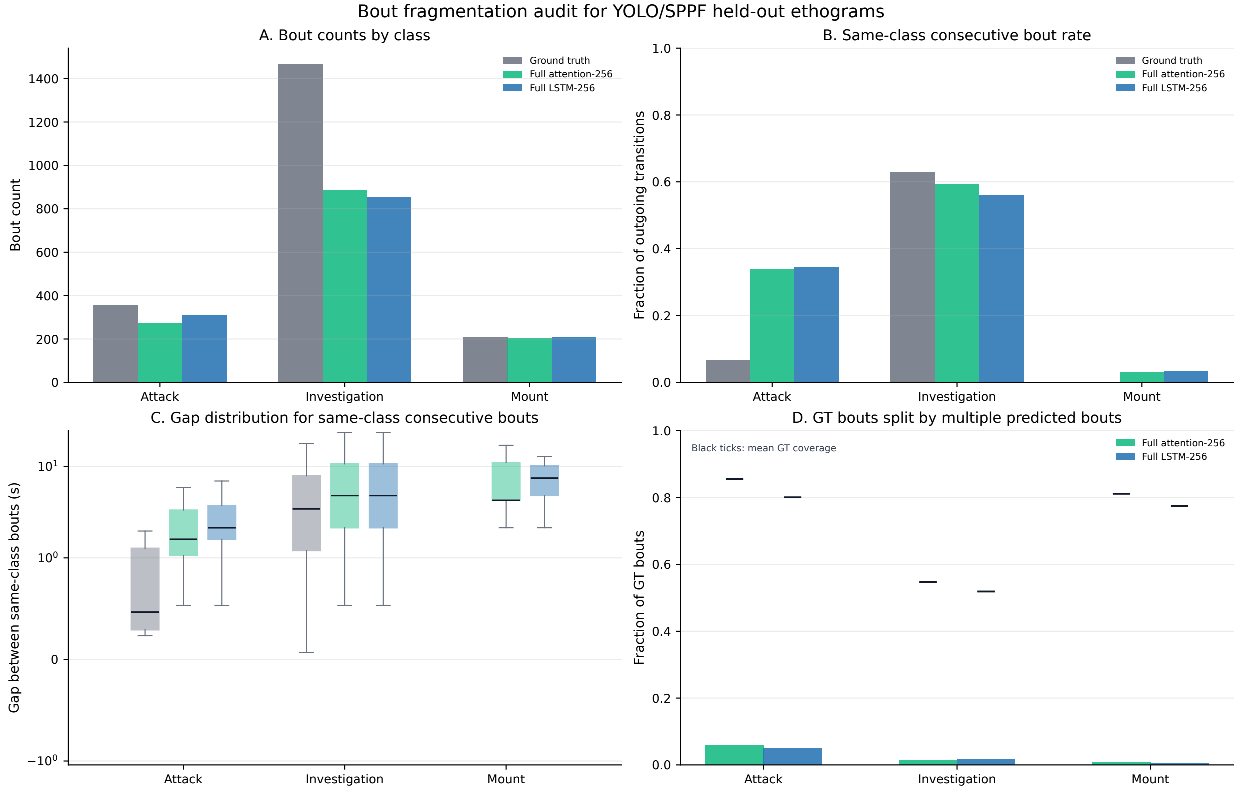

**Figure S4: Bout fragmentation and compression analysis.** Raw and postprocessed predictions are compared with ground-truth bout structure to test whether smoothing and minimum-duration filtering explain the observed bout-count differences. The diagnostic supports the main-text interpretation that the MARS limitation is primarily bout-density and under-segmentation, especially for investigation, rather than an artifact introduced only by postprocessing.

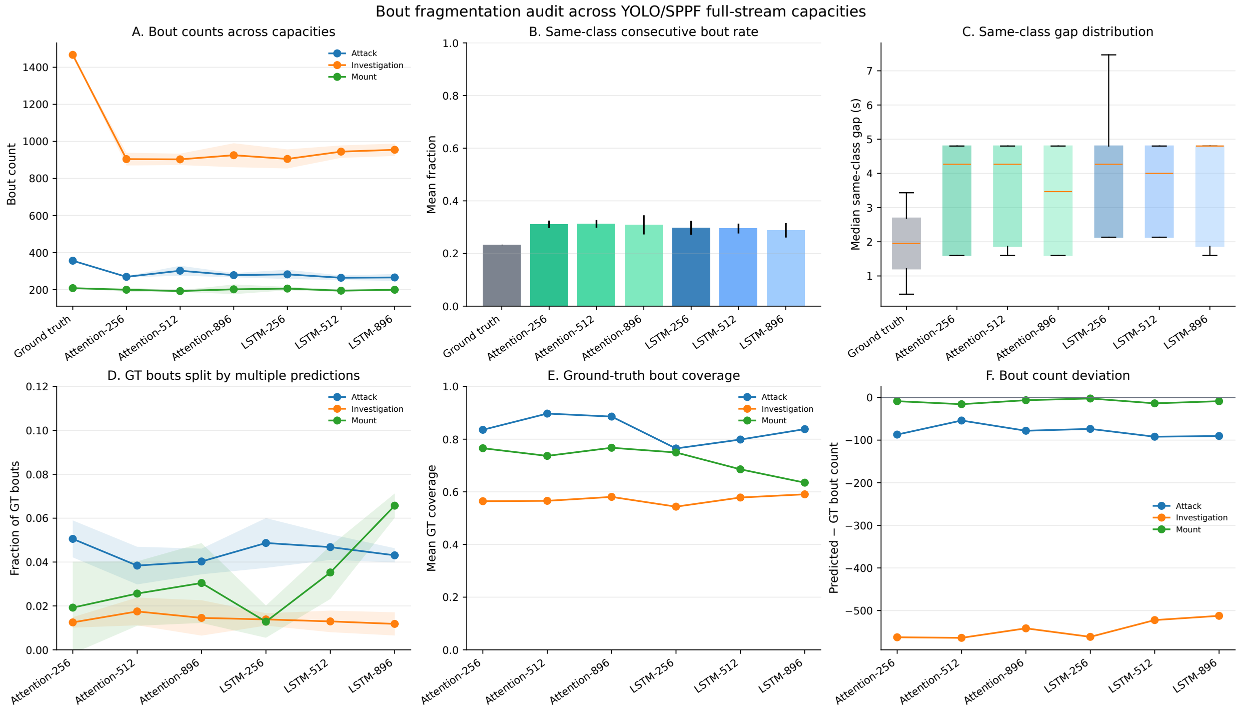

**Figure S5: Bout-structure analysis across full-stream capacities.** Full-stream LSTM and attention models are compared across capacities and seeds for bout-count error, same-class transitions, ground-truth splitting, and bout coverage. This figure extends the transition analysis beyond the selected 256-capacity models and shows that transition-matrix differences are not explained only by short-gap fragmentation of individual ground-truth bouts.

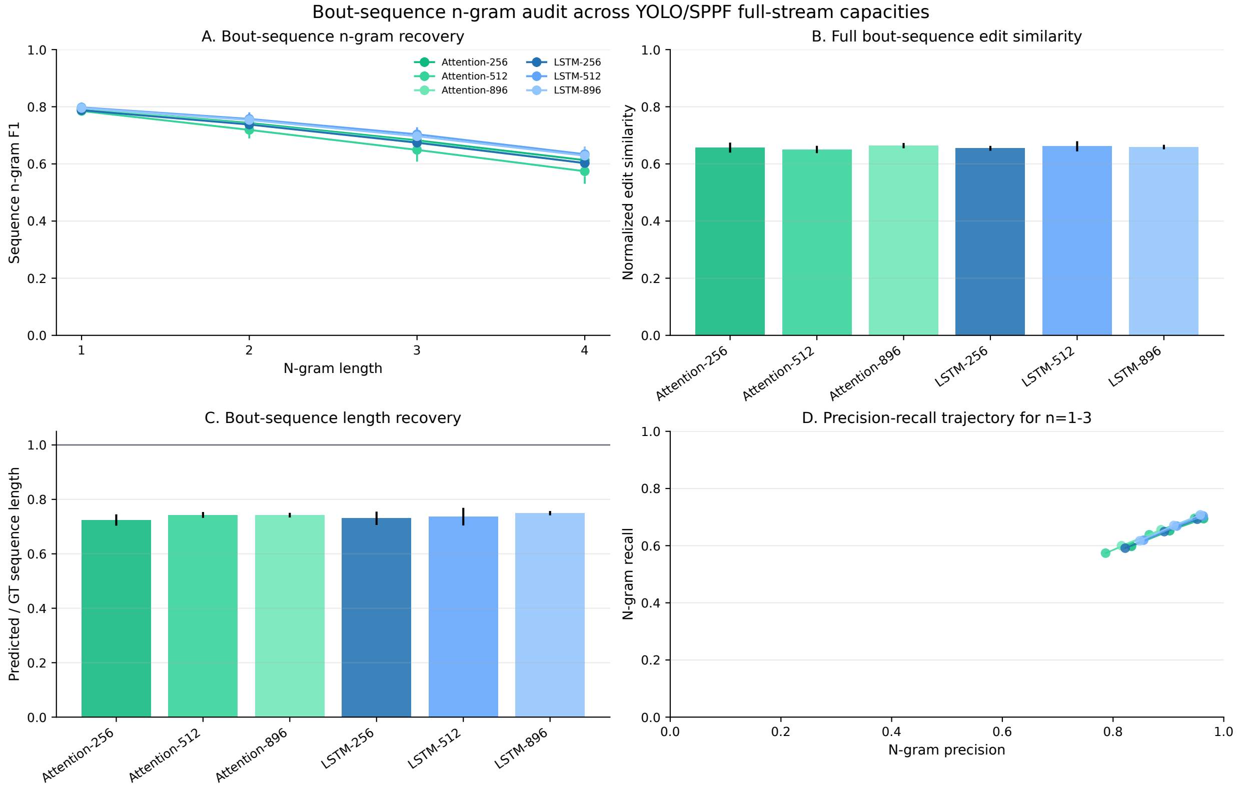

**Figure S6: Bout-sequence n-gram recovery.** Predicted and ground-truth ethograms are compared as ordered bout sequences, with n-gram recovery evaluated against composition-shuffled and Markov sequence controls. This diagnostic asks whether predicted ethograms retain behavior-order structure despite imperfect bout boundaries and compressed bout counts.

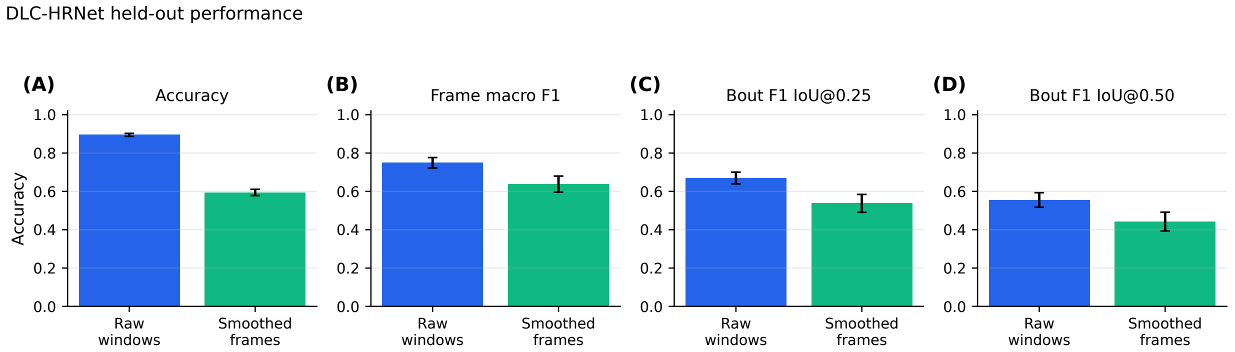

**Figure S7: DLC-HRNet held-out raw-window and smoothed-frame performance.** (A) Accuracy, (B) frame macro-F1, (C) bout F1 at TIoU 0.25, and (D) bout F1 at TIoU 0.50 are summarized across the 28 accepted held-out MARS videos. Raw-window decoding is retained because smoothing reduced bout-level performance for this DLC-HRNet run.

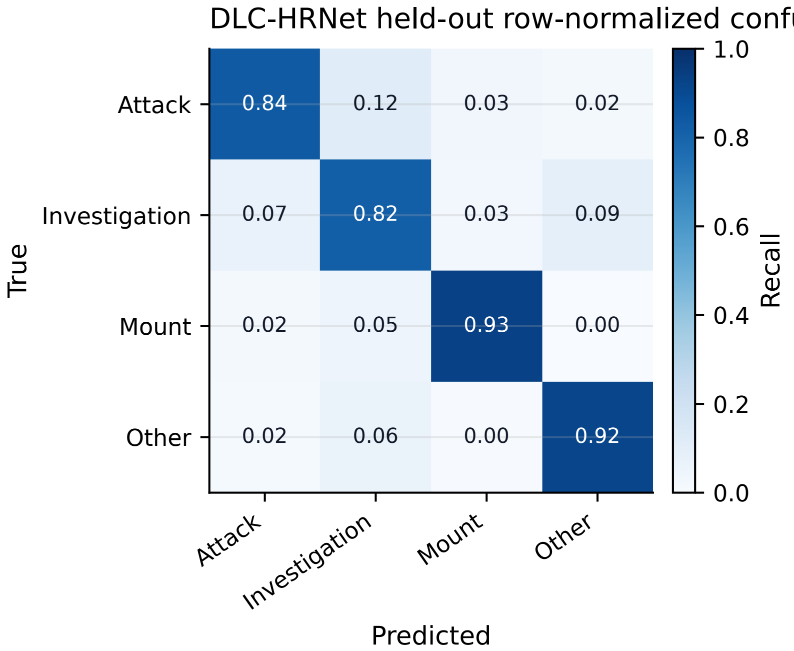

**Figure S8: DLC-HRNet held-out row-normalized confusion matrix.** Rows are ground-truth classes and columns are predicted classes for the raw-window DLC-HRNet held-out evaluation. The diagonal summarizes class-specific recall across attack, investigation, mount, and other.

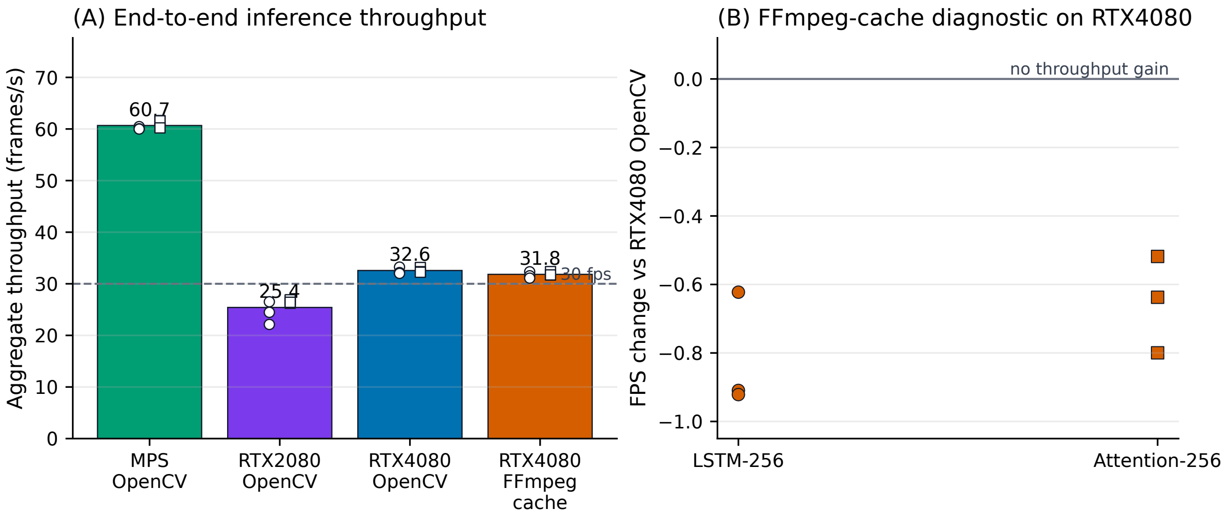

**Figure S9: End-to-end benchmark throughput.** (A) Aggregate throughput across six model-video runs for Apple Silicon MPS, RTX2080, and RTX4080 systems using the standard OpenCV path, plus an RTX4080 FFmpeg predecode/cache diagnostic. Points show individual model-video runs for LSTM-256 and attention-256. The dashed line marks real-time 30 frames/s video analysis. (B) The FFmpeg-cache diagnostic did not increase RTX4080 throughput relative to the standard OpenCV path for the matched model-video runs.

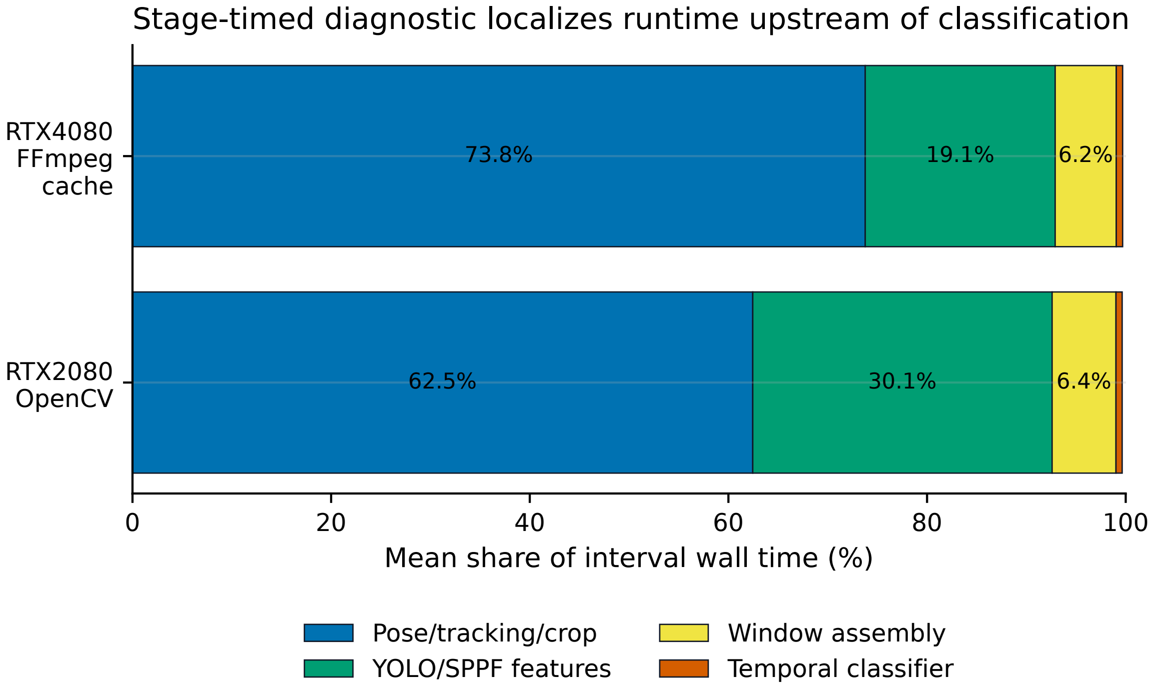

**Figure S10: Stage-timed deployment diagnostics.** Mean interval wall-time shares are shown for the stage-timed RTX2080 OpenCV run and RTX4080 FFmpeg-cache diagnostic. The combined pose, tracking, result-transfer, and crop-construction path dominated runtime, followed by frozen YOLO/SPPF feature extraction. Temporal sequence classification accounted for less than 1% of interval wall time.

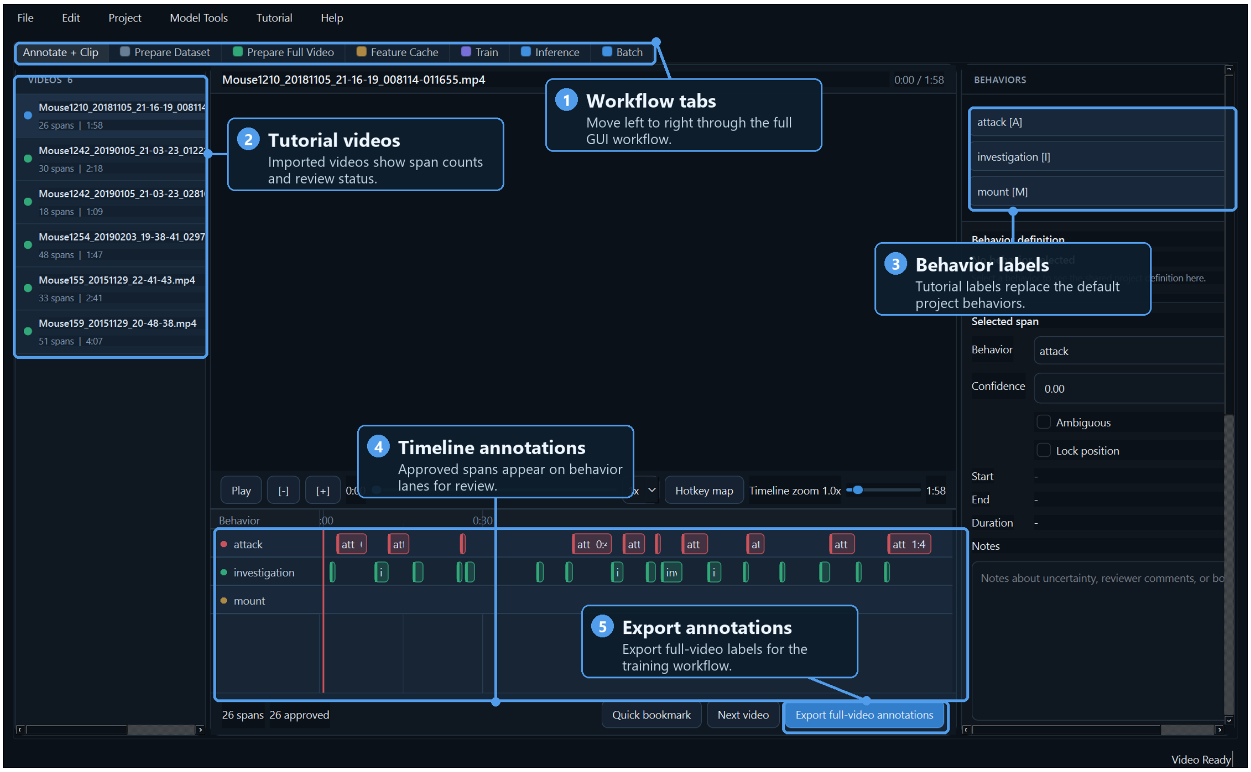

**Figure S11: GUI annotation and tutorial review workflow.** The annotation tab provides the left-to-right workflow tabs, imported tutorial videos, project behavior labels, timeline-level annotation spans, and full-video annotation export. Approved spans can be reviewed on behavior lanes before export, allowing the same GUI project to produce BENTO-compatible labels for full-video BehaviorScope-X training.

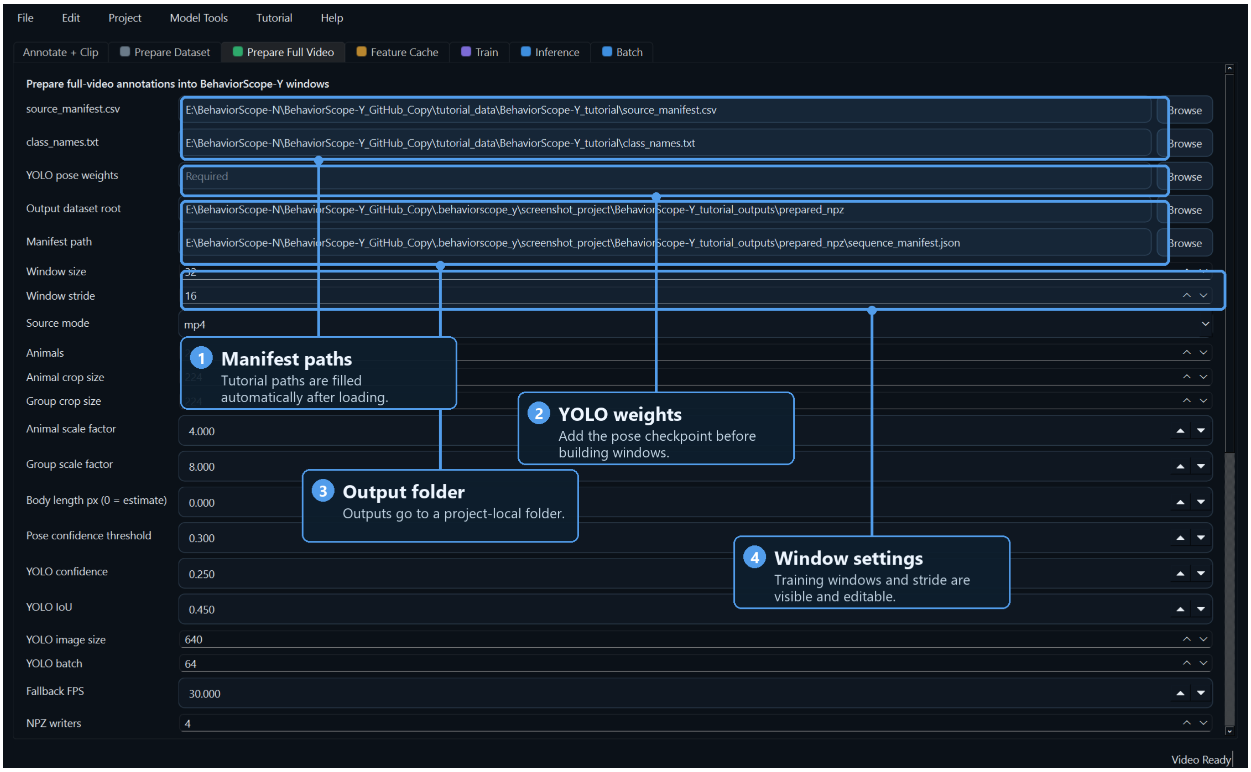

**Figure S12: GUI preparation of full-video training windows.** The full-video preparation tab collects the source manifest, class-name file, YOLO-pose checkpoint, output dataset root, manifest path, animal count, crop geometry, window length, and stride. These fields map to the full-video NPZ preparation command and define the cached sequence windows used for downstream classifier training.

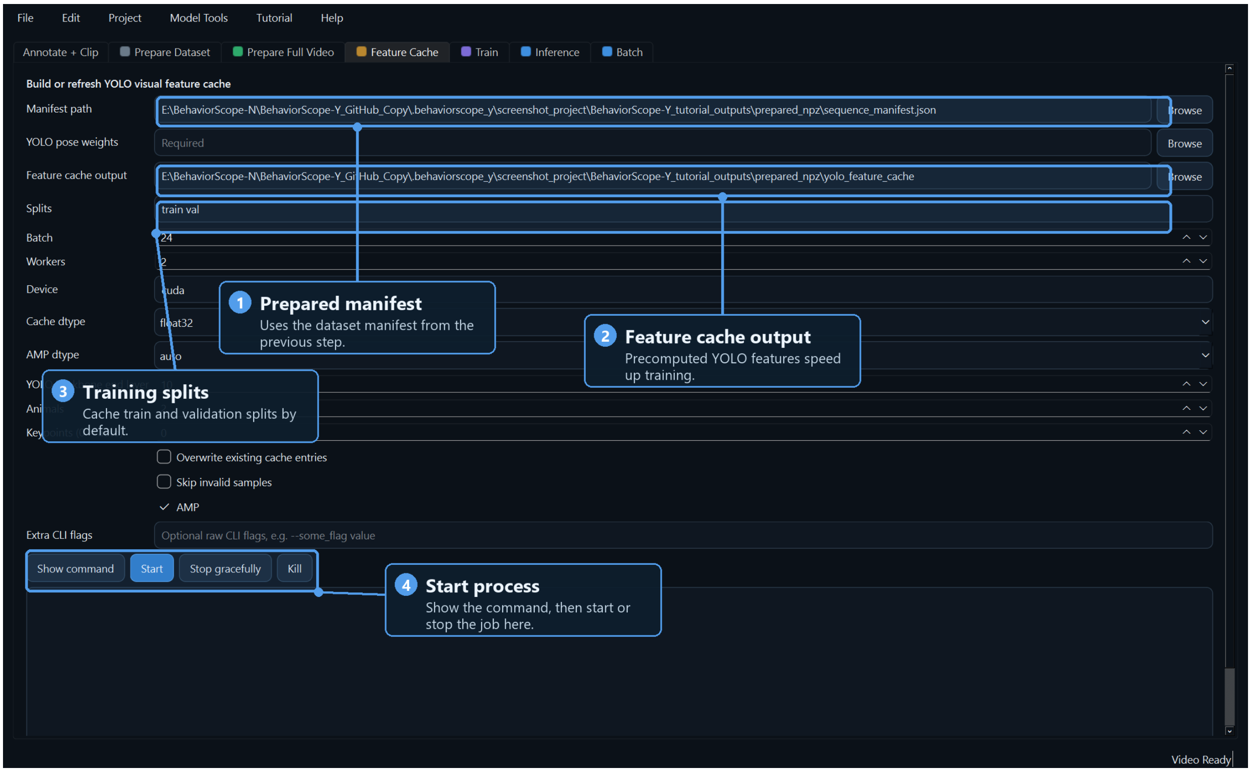

**Figure S13: GUI construction of the YOLO visual feature cache.** The feature-cache tab reuses the prepared sequence manifest and selected YOLO-pose checkpoint to precompute frozen visual descriptors for the train and validation splits. The output directory, batch size, worker count, device, cache precision, and AMP controls are exposed before launch so the cached representation can be rebuilt or reused reproducibly.

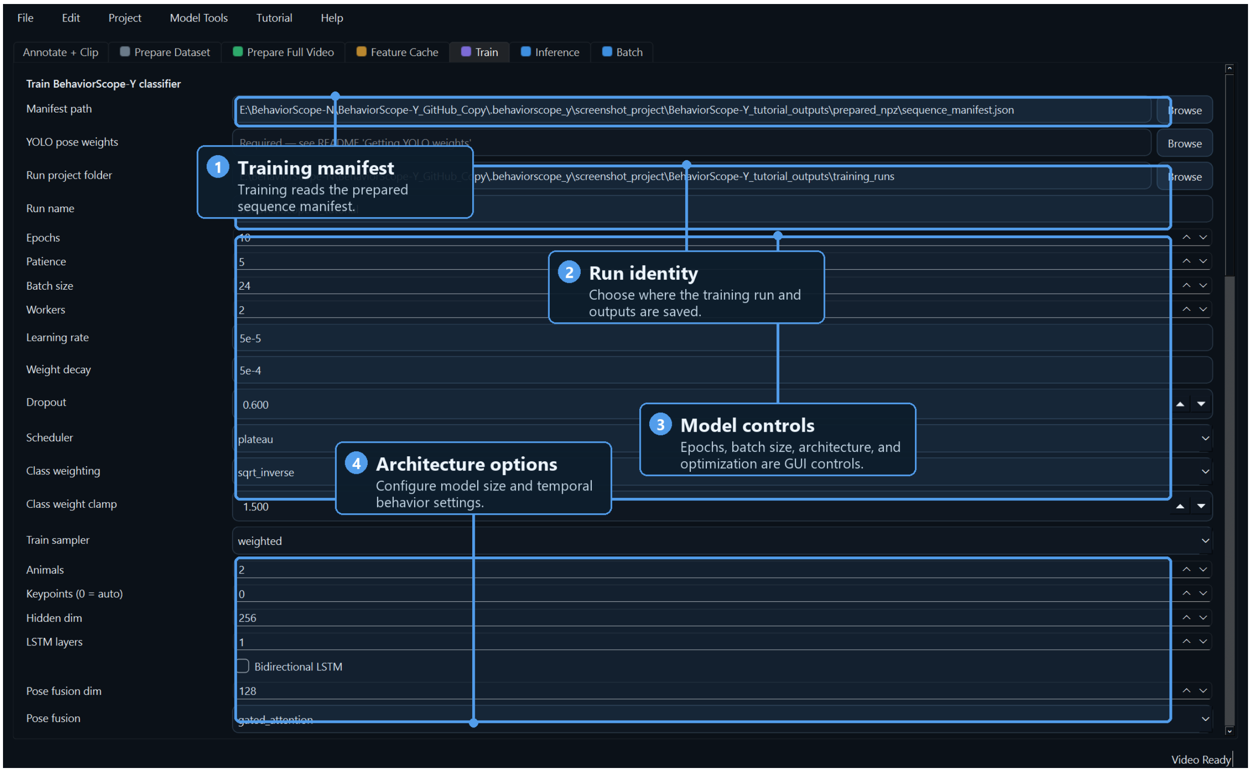

**Figure S14: GUI temporal classifier training controls.** The training tab reads the prepared sequence manifest, records the run folder and run name, and exposes optimization, class-weighting, sampler, animal-count, keypoint-count, hidden-dimension, recurrent-layer, and pose-fusion controls. These settings define the temporal classifier run while preserving the pose checkpoint and cached input manifest used to generate the training data.

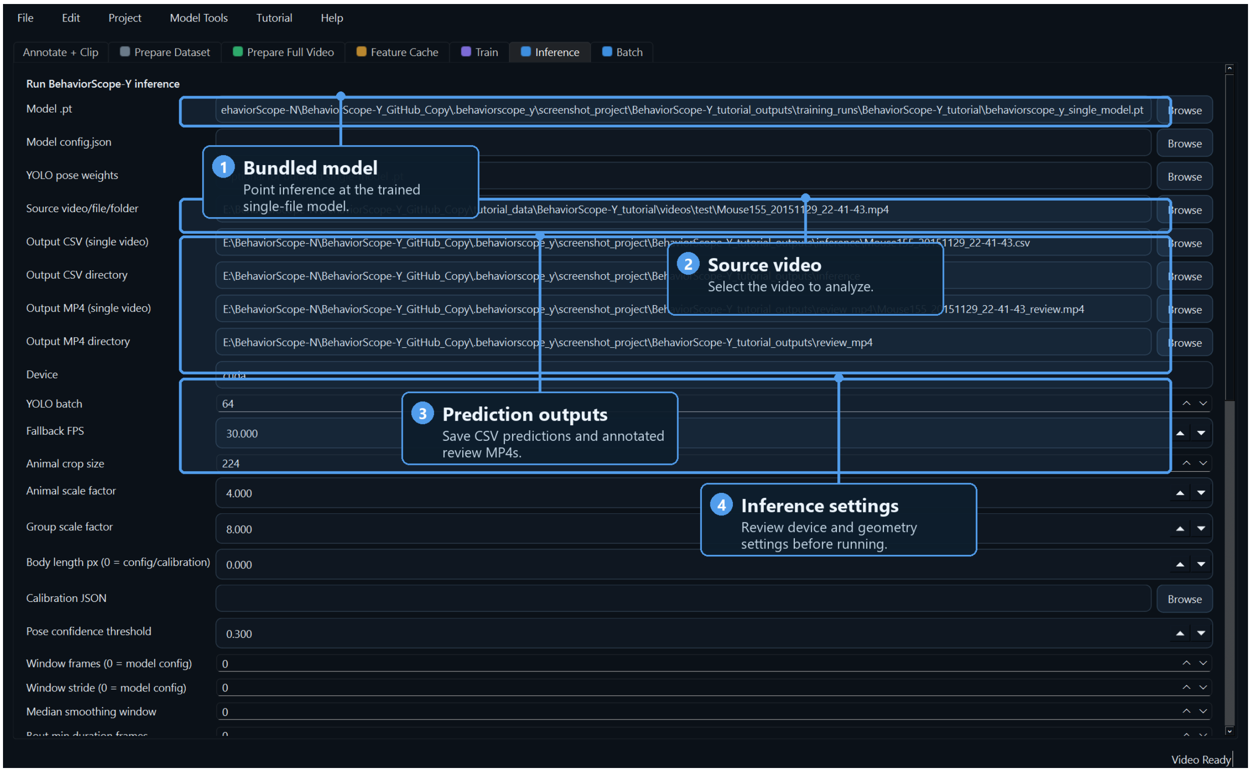

**Figure S15: GUI single-video inference workflow.** The inference tab accepts a bundled BehaviorScope-X model, an optional external model configuration or YOLO-pose checkpoint, a source video, and output paths for prediction CSV files and annotated review videos. Geometry, device, batch, frame-rate, smoothing, and bout-filter controls remain visible so deployment-time prediction settings can be checked before running a new video.

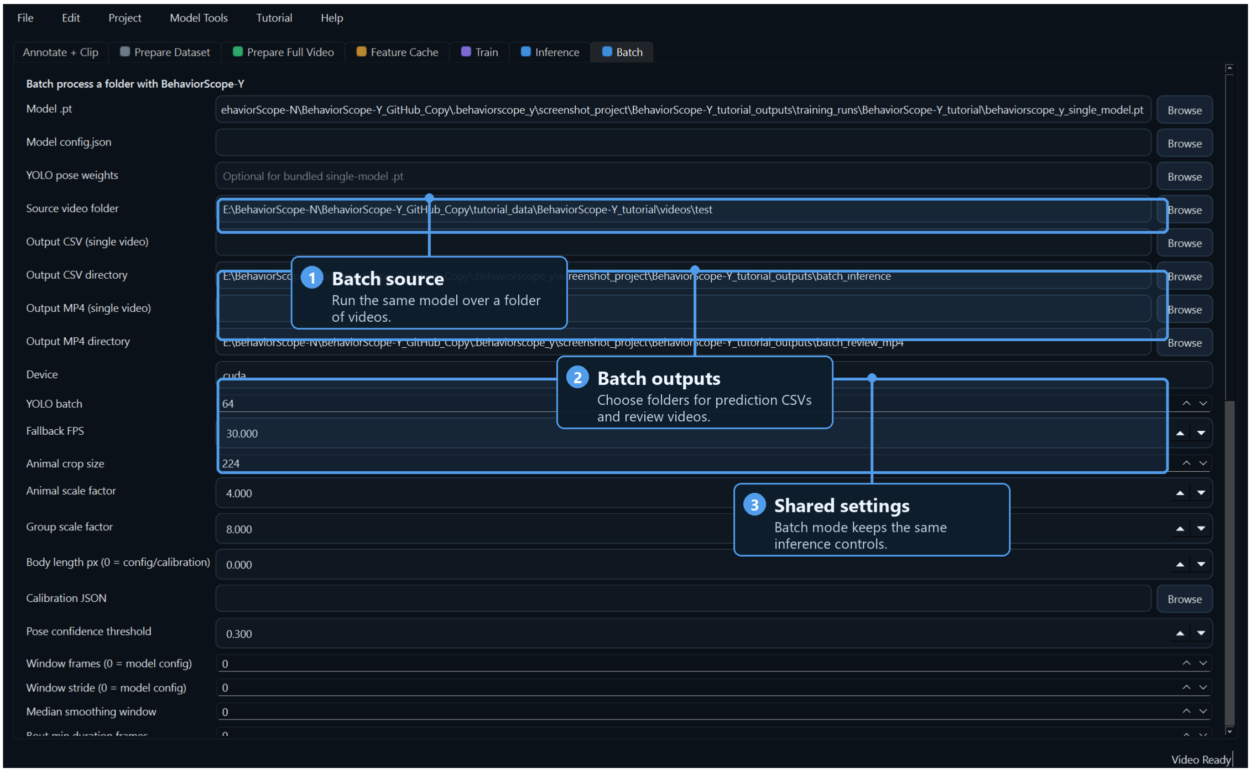

**Figure S16: GUI batch-inference workflow.** The batch tab applies the same bundled model and inference controls to a folder of source videos. Separate output directories for prediction CSVs and annotated review MP4s allow the same model to be deployed across many videos while retaining the device, crop geometry, calibration, smoothing, and bout-filter settings used for the single-video path.
